# Nanocluster-antibody-drug conjugates (NADC) as an intravesical precision theranostic agent for interstitial cystitis

**DOI:** 10.1101/2024.04.20.590432

**Authors:** Zhijun Lin, Wanyan Wang, Dingxin Liu, Qiwei Liu, Chenyixin Wang, Zhifeng Xu, Zhiming Wu, Xiangfu Zhou, Xiaodong Zhang, Yong Huang, Qi Zhao, Jiang Yang

## Abstract

Interstitial cystitis (IC) is a chronic inflammatory bladder disorder with recurring severe pain, lacking timely diagnostic and therapeutic options. Here, we propose a unitary theranostic nanocluster-antibody-drug conjugate (NADC) by covalently placing dihydroorotate dehydrogenase inhibitors (DHODHi) and ultrasmall gold quantum clusters (AuQCs) on a nerve growth factor (NGF) antagonistic antibody with simultaneous X-ray computed tomographic and near-infrared fluorescence imaging contrasts. Combining anti-inflammatory effects from all individual components, intravesical NADC specifically homed to mucosal lesions with NGF overexpression in voided bladder and capably alleviated inflammation in chronic, acute, and prophylactic IC models of rats, as revealed by behavioral and pathological evaluations. Transcriptomics unveiled cytokine modulation and concomitant inhibition of perturbed IL-17, NF-κB, TNF, and JAK-STAT signaling pathways. Notably, NADC indirectly remodeled the host bladder microbiota by differentially varying anti-inflammatory and pro-inflammatory bacteria diversities. Distinct from conventional nanoparticles conjugated with antibodies and drugs, NADC relies on the antibody framework, outperforms clinical standard-of-care agents, and represents an emerging category of precision theranostic agents with translational potential for IC theranostics in clinical practice.

## Introduction

Interstitial cystitis/bladder pain syndrome (IC/BPS) is a prevalent and chronic disease with overloaded healthcare expenses but without a cure at present(1). IC patients suffer from debilitating pelvic pain, bladder pressure, and urinary frequency and urgency resulting from bladder lining inflammation. Lacking definitive biomarkers, the diagnosis of IC is rather complex, mandating a combination of tests including but not limited to bladder diary reference, pelvic exams, cystoscopy, potassium sensitivity tests, urine cytology, and even invasive biopsy(2). However, with unelucidated etiology and symptoms overlapping with other urinary disorders, IC is frequently left with limited therapeutic options. Represented by clinically approved Rimso-50 (50% DMSO), localized intravesical therapy has been proven to be advantageous over oral medications, nerve stimulation, and surgery due to reduced systemic side effects, enriched local concentrations of drugs, high response rates, and minimal invasiveness(3). Nevertheless, current intravesical therapies, such as guideline-recommended lidocaine, heparin, and DMSO, only provide temporary symptom relief and risk systemic toxicity through leaking into circulation. Moreover, these intravesical agents demand frequent instillations owing to low bladder retention, urinary washout, and poor urothelium entry, which significantly reduce therapeutic efficacy(4). Therefore, there is a pressing need to develop IC-specific targeted agents to achieve sustained efficacy, preferably with imaging features to reliably diagnose IC, visualize drug distribution patterns, and reduce unnecessary examinations.

Ideal IC management should bilaterally direct anti-inflammation and relieve pain. NGF is expressed as a pain mediator in bladder tissues through sensitizing afferent nerve pathways associated with IC(5). NGF is first synthesized as a glycosylated precursor with retained receptor binding affinities and then proteolytically processed into mature NGF, both of which can induce autocrine/paracrine loops(6, 7). Hinged on readily known NGF receptors of TrkA, p75NTR, and SorCS, NGF inhibition can disrupt autocrine signaling as an alternative strategy for blocking only one receptor, presumably yielding reduced normal tissue toxicity and reserved functions of other ligands(8–10). NGF inhibitors like tanezumab, fulranumab, and fasinumab have demonstrated clinically meaningful benefits in pain resolution in IC patients(11). However, as observed in clinical studies, intravenous administration of these monoclonal antibodies inevitably arouses severe systemic toxicity, such as headaches, upper respiratory tract infection, increased joint damage, and peripheral neuropathy, which bring about safety concerns and jeopardize regulatory approvals but could be circumvented via intravesical routes(11). Meanwhile, non-invasive intravesical therapy can preserve high local concentrations of drugs by direct exposure. DHODH is required to produce immunomodulatory cytokines like IL-6 and IL-17A(12). Recent studies revealed the anti-inflammatory and immunomodulatory effects of selective DHODHi via suppressing mitochondrial pyrimidine synthesis, exemplified by clinically investigational vidofludimus (IMU-838)(13). While elevated levels of IL-17A are found in the bladder tissues and urine of IC patients, IL-17A triggers the release of other pro-inflammatory cytokines(14). Immunosuppressive vidofludimus can downregulate the expression of IL-17A by inhibiting the downstream NF-κB signaling pathway to attenuate chronic inflammation and autoimmune tissue damage(14). To our knowledge, the indication of vidofludimus has not been expanded to treat IC.

The clinical success of antibody-drug conjugates (ADCs) as a targeted therapy has gone beyond oncology, with investigational examples like ABBV-3373 and DSTA4637S(15–18). Even though ADCs and derivatives, including immune-stimulating antibody conjugates (ISAC) and immune-modulating antibody-drug conjugates (IM-ADC), can enhance efficacy and minimize off-target side effects on normal tissues by precision targeting, they lack the visualization for therapeutic surveillance. We hypothesize that anti-NGF in NADC can transiently associate with cell surface receptors during autocrine/paracrine signaling, followed by internalization for lysosomal payload release. In this study, we covalently engineered an intravesical theranostic NADC, comprised of an NGF-targeting monoclonal antibody, immunosuppressing DHODHi, and AuQCs (Fig. 1). Unlike conventional nanoparticles (NPs), ultrasmall AuQCs are comparatively insignificant in size with reference to antibodies and exhibit anti-inflammatory effects and imaging contrasts of non-invasive near-infrared fluorescence (NIRF) and spectral CT. NADC unifies the otherwise differential pharmacological behaviors of individual drugs to achieve image-guided therapeutic synergy against IC. The combined locoregional bladder instillation and specific NGF targeting of NADCs can maximize drug precision, minimize systemic toxicity, and spare normal tissues, improving somatic pain symptoms and functional voiding in rat models of chronic, acute, and prophylactic IC. The transcriptome and microbiome analyses identify multifaceted inhibition of inflammatory signaling pathways of IL-17, NF-κB, TNF, and JAK-STAT and differential modulation of dysregulated microbiota populations in IC. The versatile NADC framework represents theranostic convergence of synergistic precision medicine that can safely and effectively treat IC with high NGF expressions in bladder mucosae under image guidance.

**Figure 1.**
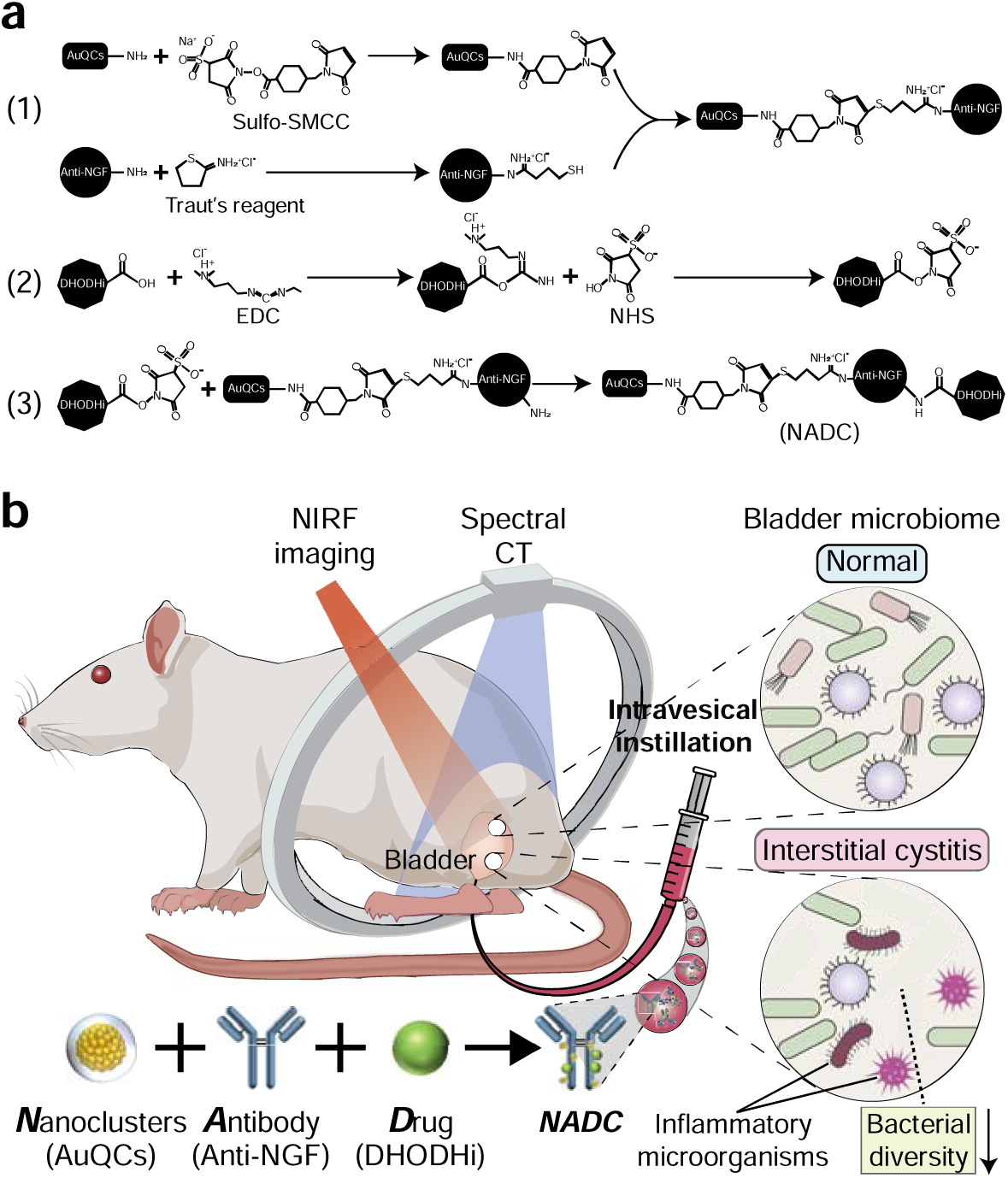
Synthesis pathways of NADC and concept of intravesical theranostic NADC against interstitial cystitis (IC). (a) Schematic diagram of the synthesis process of NADC. (1) Solvent-accessible primary amino groups of AuQCs (black square) react with sulfo-NHS ester of sulfo-SMCC crosslinker to acquire maleimide-terminated derivatives reactive towards sulfhydryls. Concurrently, primary amines from the lysine side chains of the anti-NGF tanezumab antibody (black circle) react with the cyclic thioimidate compound, Traut’s Reagent, to introduce a reactive sulfhydryl group in one step that is successively crosslinked to maleimide-activated AuQCs. (2) The benzoic acid group of vidofludimus (black octagon) is activated by EDC to form the O-acylisourea ester intermediate, which is in turn stabilized by sulfo-NHS to derive the amine-reactive DHODHi molecule. (3) The remaining unreacted amines on tanezumab are covalently crosslinked with the DHODHi NHS ester to yield NADC. (b) Theranostic AuQCs and the immunosuppressive DHODHi are conjugated through covalent linkage to the humanized NGF antagonistic monoclonal IgG2 tanezumab that targets overexpressed NGF from IC. AuQCs, vidofludimus, and tanezumab synergize in NADC and encompass concomitant regulation of inflammation-associated signaling pathways and remodeling of the host bladder microbiota. Intravesical NADC allows bilateral management of IC by simultaneously alleviating inflammation and pain under dual-modality NIRF and CT image guidance.

## Results

### NGF is upregulated in human bladders with IC

First, we conducted a comprehensive single-cell RNA sequencing (scRNA-seq) analysis to differentiate specific cell types within human IC tissues and accurately investigate their NGF expression levels. Based on gene expression profiles and corroborating evidence from existing literature, our analysis identified 20 distinct cellular clusters at a resolution of 0.2 (Fig. 2a,b), which were then classified into 12 distinct cell types. The overall bladder NGF expression level is markedly higher in IC patients than in healthy individuals (Fig. 2c). Among others, epithelial cells emerge as the predominant cell population in bladder tissues of IC patients, with unique subpopulations overexpressing NGF compared to the healthy control group (Fig. 2d,e), particularly in epithelial cell-2 characterized by KRT7, SNCG, and KRT13 expression. Although NGF expression in epithelial cell-1 co-expressing KRT19, KRT18, and KRT17 did not reach statistical significance, the mean level was notably higher in IC patients (Fig. 2d). Gene Ontology (GO) enrichment analysis suggests the clinical relevance of the epithelial cell-2 subpopulation with vital bladder barrier functions in terms of cell junction integrity, adhesion, and tight junction organization (Fig. 2f). This IC-related epithelial cell-2 subpopulation is also intricately linked to multiple inflammation-related pathways (Supplementary Fig. 1). To further validate our findings, we chemically induced a recognized rat model of IC using gradient doses of cyclophosphamide (CYP) to correlate the severity of inflammation and bladder damage with in situ NGF expression. Immunofluorescence revealed a dose dependence of NGF expression primarily in the urothelium and epithelial cells, corresponding to induction doses (Fig. 2g and Supplementary Fig. 2). Immunoblotting of homogenized bladder tissues corroborated a similar dose-dependent pattern of NGF expression (Fig. 2h), underscoring its pivotal role in disease onset and exacerbation. The scRNA-seq highlights the dual functions of subpopulated bladder epithelial cells as both a barrier and a primary source of NGF expression, thereby positioning NGF as an effective therapeutic target for treating IC. Building on these insights, we have designed NADC with integrated imaging and therapeutic multimodalities to halt bladder inflammation and pain symptomatology.

**Figure 2.**
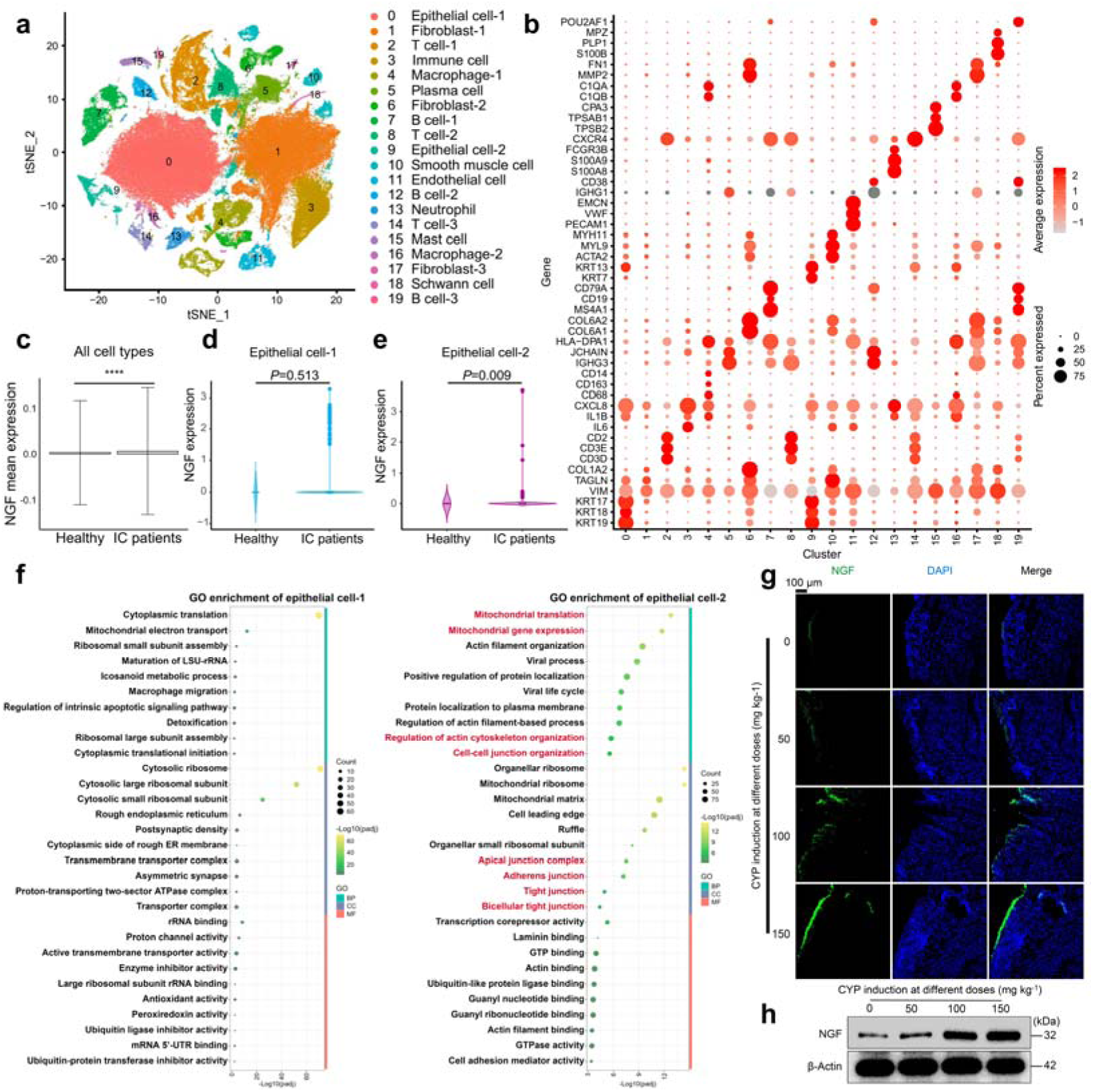
NGF is upregulated in the bladder of humans with IC. (a) A t-SNE plot displayed 20 clusters at a resolution of 0.2, with the epithelial cell population being notably distinct among all cell types. (b) A dot plot highlighted marker genes specific to different cell types. (c) Mean NGF expression across all cell types was significantly elevated in IC patients compared to healthy individuals (p ≤ 0.001 from *t*-test). (d,e) Violin plots depicted NGF expression specifically in epithelial cells between healthy and IC groups. (f) GO enrichment analysis of marker genes for epithelial cell-1 and epithelial cell-2 based on single-cell data, showing pathways colored from green to yellow indicating increasing -log(p.adjust) values, with circle size denoting the number of enriched genes. (g) Immunofluorescence images of bladder tissues from IC rat models, induced with varying CYP doses, demonstrated increasing NGF expression with higher induction doses. (h) Corresponding western blot images confirmed NGF expression levels in bladder tissues.

### Preparation and characterization of NADCs

To synthesize NADCs, we utilize a two-step conjugation strategy (Fig. 1a). First, a proportion of amines on tanezumab were sulfhydryl-activated by Traut’s reagent, with an increased negative zeta potential (Supplementary Fig. 3). In parallel, AuQCs were covalently coupled to the amine-reactive *N*-hydroxysuccinimide (NHS) ester of a hetero-bifunctional sulfo-SMCC to form stable amides. Next, the thiolated anti-NGF antibody reacted with maleimide-tethered AuQCs to derive thioether bonds. Last, the remaining primary amines of the complex were covalently crosslinked to the benzoic acid of vidofludimus with NHS-EDC to attain NADC. Accompanied by sequential modifications of tanezumab by AuQCs and DHODHi, we observed increases in the hydrodynamic size from 7.5±0.5 to 28.2±4.5 nm and the zeta potential from −7.6 ± 0.7 mV to −16.7 ± 0.7 mV (Fig. 3a,b). While AuQCs were ultrasmall under 3 nm as visualized by transmission electron microscopy (TEM), NADC reached 25 nm in diameter due to the relatively large size of immunoglobulins (Fig. 3c,d and Supplementary Fig. 4). NADC well-preserved characteristic absorption peaks of DHODHi at 260 and 290 nm, as well as the anti-NGF peak at 280 nm that was broadened by AuQCs conjugation (Fig. 3e). Furthermore, a minor hypsochromic shift in the NIRF emission of AuQCs was perceived following covalent modifications into AuQCs-anti-NGF or NADC (Fig. 3f). We then calculated the molar nanocluster-to-drug-to-antibody ratio (NDAR) to be 4.7:3.4:1, which warrants dual DHODHi and theranostic AuQCs payloads to concurrently alleviate overactive immune reactions and navigate diagnostic imaging (Fig. 3g). The molecular weight (MW) of NADC was studied by non-reducing native polyacrylamide gel electrophoresis (PAGE), resulting in a band over 180 kDa (Supplementary Fig. 5). The exact MW and compositions of NADC were further confirmed with MALDI-TOF-MS (Supplementary Fig. 6). DHODHi is in the sodium adduct form, and the MALDI spectrum of AuQCs is dominated by a major peak of α-LA ligands at 14.1 kDa, with clusters calculated to be Au_38_ dimerizing from Au_19_(19). Tanezumab shows a typical IgG MW around 148 kDa and turns 217.5 kDa after covalent conjugation with AuQCs. The NDAR ratio in NADC was quantified through m/z shifts in consistency with experimentally measured results. In conclusion, given the relative dimensions of individual constituents in the complex, the NDAC is fundamentally dissimilar from conventional antibody-and-drug-conjugated NPs (Fig. 3h).

**Figure 3.**
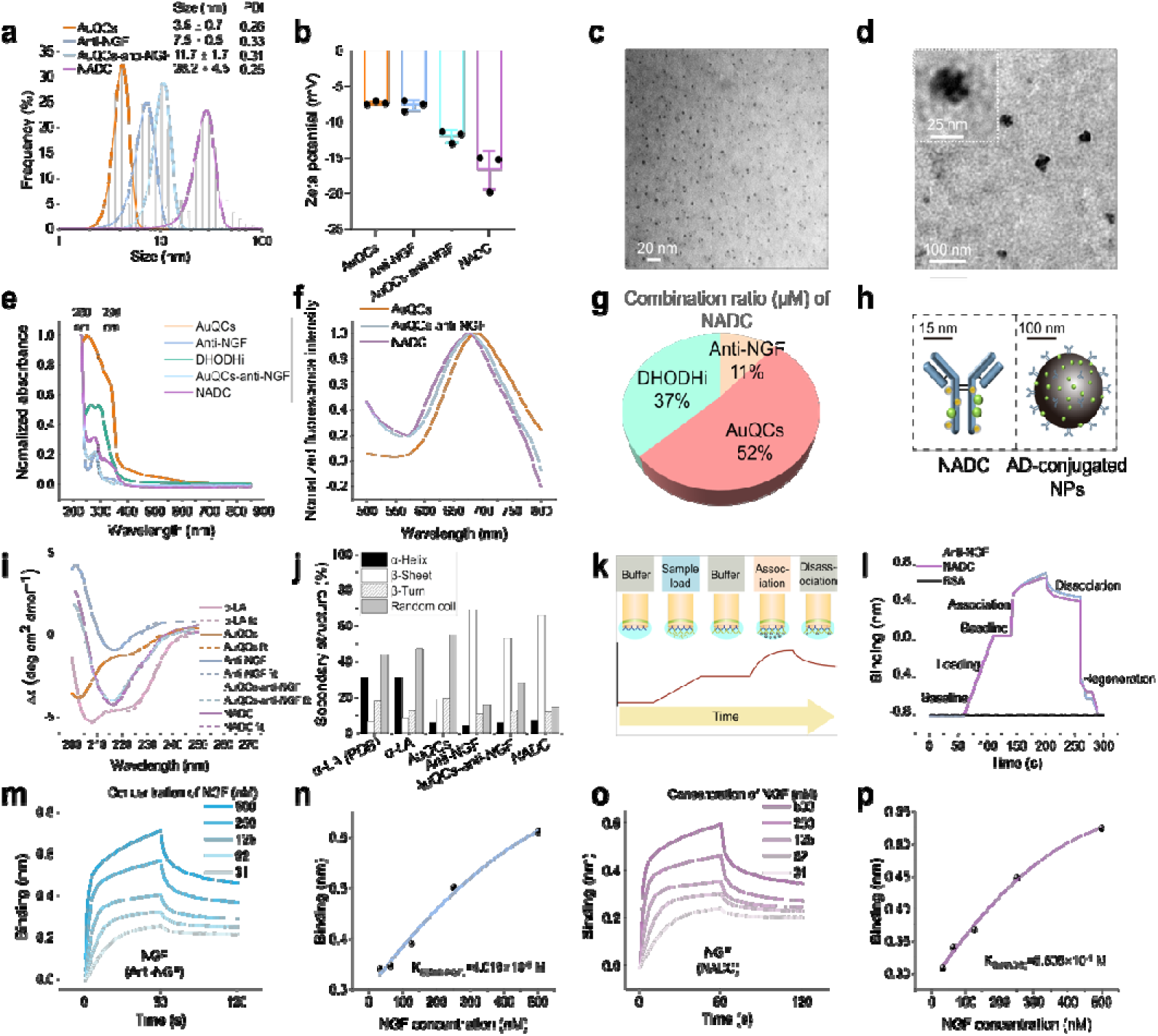
Characterization of NADC. (a) DLS analysis for the hydrodynamic size distribution and (b) zeta potential of NADC and individual constituents (n=3). Representative TEM images of (c) ultrasmall-sized AuQCs and (d) NADC. (e) Normalized UV-vis absorption spectra showing distinct feature peaks of DHODHi at 260 and 290 nm (dotted lines) and a broadened peak at 280 nm of tanezumab in NADC. (f) Normalized fluorescence emission spectra of NADC at the excitation wavelength of 360 nm. (g) The molar NDAR of each constituent in NADC. (h) A schematic comparison between NADC and conventional antibody-drug-conjugated NPs. (i) Far-UV CD spectra of NADC and individual constituents. The dotted lines indicate fitted curves by applying the BeStSel algorithm to data for secondary structure determination and folding recognition. (j) Calculated secondary structure compositions of NADC, with structural components predominantly contributed by the anti-NGF antibody as the framework. The structure of apo-state bovine α -LA retrieved from RSCB (PDB ID: 1F6R) was quantified to be consistent with experimental data. (k) Schematic representation of BLI sensorgrams showing interferometric binding signal changes with assay processes. (l) BLI sensorgrams of the loading, binding, association, and disassociation steps of anti-NGF antibody and NADC, with BSA as a control. Tanezumab and NADC were immobilized on the AHC biosensor surface by anti-human IgG Fc captures in similar binding kinetics. (m and n) Real-time binding kinetics of the anti-NGF antibody and (o and p) NADC by BLI for interactions with gradient NGF concentrations at 2-fold increments. (n and p) Steady-state analyses for K_D_ determination. NADC retained the high binding affinity of tanezumab at the same order of magnitude at nanomolar levels.

### Structures and binding affinities of anti-NGF antibodies are retained in NADC

Far-ultraviolet circular dichroism (CD) spectroscopy revealed the dominant contribution and rigid preservation of antibody secondary structures in NADC, with largely unaltered β-sheets at 216 nm, β-turn broad bands at 220-230 nm, and flexible random coils at 200 nm (Fig. 3i). Notwithstanding the dominance of tanezumab in CD signatures of NADC, α-LA in AuQCs still contributed to an elevated α-helix proportion, with ambiguous but distinguishable negative bands at 222 and 208 nm in NADC (Fig. 3i,j). To assess whether antigen recognition and binding are still viable in the NADC scaffold after conjugation with AuQCs and DHODHi, we performed label-free biolayer interferometry (BLI) to analyze interference patterns upon interactions with NGF (Fig. 3k). Real-time BLI sensorgrams of the anti-NGF antibody and NADC rendered coinciding steps in the loading, association, and dissociation phases, suggesting resembling probe immobilization and antigen-antibody binding kinetics (Fig. 3l). In the meantime, minimal interactions were observed in all phases for bovine serum albumin (BSA). After probe immobilization on the sensor surface, affinity binding dynamics were measured for anti-NGF and NADC at gradient NGF concentrations ranging from 31 to 500 nM (Fig. 3m-p). Fitting of maximum wavelength shifts by Michaelis-Menten substrate-binding kinetics yielded equilibrium dissociation constants (K_D_) of 4.019×10^-9^ M for anti-NGF and 8.505×10^-9^ M for NADC at equivalent orders of magnitude (Fig 3n,p). Collectively, these results elaborated on the successful preparation of NADC with the rigid antibody conformations and NGF-targeting affinities sufficiently retained as the prerequisite for subsequent IC theranostics.

### NADC as a dual-modality imaging contrast agent for IC

To scrutinize the diagnostic features of NADC, we administered the NADC agent by intravesical instillation to rats with CYP-induced urinary bladder inflammation, a well-established preclinical IC model. AuQCs and NADC could readily delineate bladder contours with fast contrast enhancement in a trice (Fig. 4a,b). Time-course NIRF imaging revealed a classic one-compartment open model, where administered contrast agents were instantaneously distributed in the inflamed bladder as a uniform body compartment. In rats with IC, AuQCs were rapidly cleared with only minor NIRF signals seen after 3 h, whereas NADC was dynamically sustained with prolonged retention and identifiable signals even after 12 h (Fig. 4a). The bladder retention half-life of NADC was experimentally determined to be 4.6±0.3 h, much higher than 1.2±0.1 h of AuQCs (Fig. 4b). In contrast, the NIRF intensities of both NADC and AuQCs instantly and similarly decayed in healthy rats within 3 h post-instillation, with comparable half-lives of 1.5± 0.2 h and 1.6 ± 0.2 h, respectively (Fig. 4c,d). Next, to confirm that the extended half-life of NADC is attributable to active NGF targeting in inflamed bladder tissues, we conducted comparative NIRF imaging in rat models using NADC and CF647-labeled anti-NGF antibodies. The successful covalent conjugation of CF647 to anti-NGF antibodies with retained NIRF was characterized and confirmed, with the particle size and zeta potential close to unmodified antibodies (Supplementary Fig. 7). Probe-specific fluorescence of NADC and anti-NGF-CF647 at identical excitations can be easily differentiated in phantoms by multispectral unmixing (Supplementary Fig. 8). The bladder retention half-lives of both probes were comparable to each other and found larger in IC rats than in healthy rats (Supplementary Fig. 9,10). Meanwhile, NGF was markedly overexpressed in the inflamed urothelial lesions at levels positively linked to induction doses (Supplementary Fig. 11). The NADC distribution was correlated and overlapped with NGF in high degrees. The results unequivocally authenticated that the antibody-directed NGF-targeting of NADC could prolong intravesical retention in the IC-irritated bladder with an improved diagnostic window and drug action.

**Figure 4.**
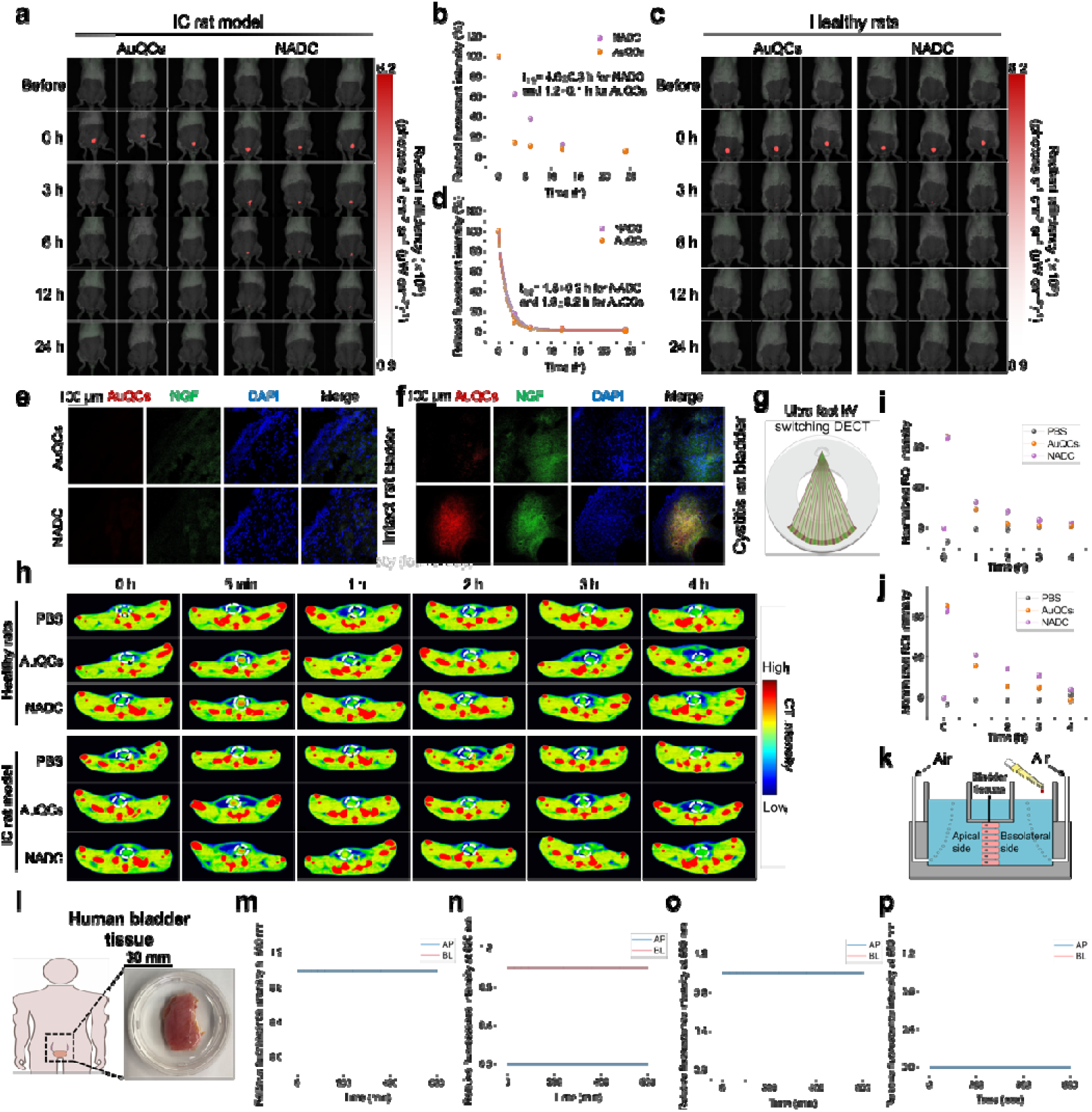
NADC as a dual-modality contrast agent for imaging IC. Time-course in vivo NIRF imaging of (a, b) rats with CYP-induced cystitis and (c, d) in healthy conditions receiving intravesical AuQCs or NADC (n=3 animals per group). Confocal laser scanning microscopy (CLSM) of bladder tissue sections from (e) intact healthy and (f) IC rats following intravesical AuQCs or NADC stained by immunofluorescence. (g) A schematic diagram showing the setup of a dual-energy CT scanner. (h) Planar spectral CT images of the bladder at various time points after intravesical administration of indicated agents to healthy rats or rats with IC (n=3 animals per group). Corresponding calculated intensities for the bladder ROIs (white dotted circles) of (i) healthy rats and (j) rats with IC. Evaluation of mucosal drug transport of NADC across human bladder tissues by Ussing chamber tests (k-p). (k) A schematic illustration of the Ussing chamber test using human bladder tissues. The chamber was perfused with gas, and NADC was introduced on one side to assess the transcellular and paracellular transports to the other side. (l) A photograph of a human bladder tissue piece surgically resected from a patient. AuQCs or NADC were collected at predetermined intervals to measure fluorescence spectra on both sides. NIRF intensities of AuQCs (m-n) and NADC (o-p) on both sides as a function of time.

As in human IC, local NGF at the inflammation sites in the preclinical IC rat model was overly expressed with higher signals from immunofluorescence than in normal bladders(20) (Fig. 4e,f). 12 h after intravesical injection, the retained NIRF signal of NADC in bladder tissues was much higher than that of AuQCs, in extensive and precise colocalization with NADC-targeted NGF. Three-dimensional (3D) volumetric imaging by Z stacks visually showed stronger, broader, and more penetrative NIRF signals of NADC, dictating long bladder residence time and partial penetration into deep bladder urothelia by actively targeting NGF (Supplementary Fig. 12). Further, the targeting capability and internalization of NADC against the inflammatory bladder mucosal epithelium at cellular levels were verified using an induced SV-HUC-1 cell model by fluorescence imaging (Supplementary Fig. 13).

On top of NIRF, the intrinsic CT contrast of NADC overcomes the limited tissue penetration depth as a complementary imaging modality, providing cross-sectional information and reinforced anatomical microstructures (Fig. 4g). Taking advantage of multi-energy spectral CT, phantom images of AuQCs and NADC in equivalent contrast can be clearly distinguished from the clinical iodine agent, Ioversol, with only minor crosstalk bleeding through iodine and non-iodine mapping (Supplementary Fig. 14a,b). The monochromatic energy attenuation curves of AuQCs and NADC are almost identical within 40-140 keV, with considerable differences in keV energy separation to discriminate Au from PBS and iodine (Supplementary Fig. 14c). The comparable CT signal intensities of AuQCs (R^2^=0.99) and NADC (R^2^=0.98) could be decomposed, linearly depending on concentrations of Au material densities with low interference from Ioversol in non-iodine projections (Supplementary Fig. 14d,e). After intravesical instillation in healthy and IC rats, bladders were immediately delineated by contrast-enhanced spectral CT using AuQCs and NADC (Fig. 4h). Time-lapse CT imaging affirmed a more deferred bladder voiding in IC rats administered with NADC than AuQCs, while expeditious emptying of both agents was found in healthy rats (Fig. 4i,j). Combining the bimodalities of NIRF and spectral CT with contrast enhancement from NADC allows us to simultaneously register temporal and spatial resolutions to diagnose bladder disorders such as IC.

### Limited bladder lining permeability of NADC into systemic circulation towards SV-HUC-1 mimicry, rat, and human urothelia

As probed earlier by 3D volumetric imaging, NADC capably pervaded the inner bladder lining and reached the suburothelium at a concrete penetration depth (Supplementary Fig. 12). We adapted the Ussing Chamber model to ascertain whether NADC possesses impermeability traits without traversing bladder tissues and triggering unintended side effects from systemic entry and absorption. First, we constructed an in vitro bladder mucosa mimicry using the immortalized human normal urothelial cell line SV-HUC-1 (Supplementary Fig. 15a). SV-HUC-1 was continuously cultured until a monolayer of high confluence and tight junctions formed and reached the saturated transepithelial/transendothelial electrical resistance (TEER) to mimic urothelium (Supplementary Fig. 15b). Next, we added AuQCs and NADC from the apical or basolateral side and then periodically monitored NIRF of the opposite sides to scrutinize whether the agents were transported across urothelium from transcytosis or paracellular space (Supplementary Fig. 16). Noticeably, NIRF signals were only observed at the ipsilateral application side but not at the contralateral side, confirming the physiological impermeability of SV-HUC-1 urothelial mimicry for both AuQCs and NADC. Likewise, we collected bladder tissues from healthy and IC rats and established ex vivo urothelia in the Ussing Chamber to evaluate the permeability (Supplementary Fig. 17,18). Surprisingly, besides the intact bladder mucosa from healthy rats, the irritated bladder urothelium from IC rats undergoing barrier disruption was neither penetrable to AuQCs nor NADC. Last, we set up a similar transepithelial ex vivo model using surgically dissected human bladder tissues with histological and microenvironmental organ characteristics of clinical relevance (Fig. 4k-p, Supplementary Fig. 19). Consistently, AuQCs and NADC were retained at the ipsilateral side and unable to cross the bladder barrier. Under image guidance, we also monitored NADC in bladder lesions and assessed the drug penetration depth. Intravesical NADC was perceived to rapidly target mucosal lesions of IC in 15 min and gradually penetrated in depth over time, reaching the submucosal layer to a limited extent of 60 μm in 12 h without entering the systemic circulation (Supplementary Fig. 20).

### NADC alleviates inflammatory responses in vitro

Before initiating in vivo studies, we verified if NADC could alleviate in vitro inflammatory responses in cells. We activated NOD-like receptor protein 3 (NLRP3) inflammasomes in SV-HUC-1 bladder epithelial cells to mimic inflammatory responses synergized by lipopolysaccharide (LPS) and adenosine triphosphate (ATP). Measured by quantitative reverse transcription polymerase chain reaction (RT-qPCR), LPS/ATP co-induction upregulated mRNA expression of pro-inflammatory cytokines of IL-1β, TNF-α, and IL-6 and downregulated anti-inflammatory cytokines of IL-4, IL-10, and IL-13 in the stimulated SV-HUC-1 model (Supplementary Fig. 21). Fortunately, the inflammation-induced differential cytokine expressions were effectively ameliorated by NADC, with each NADC constituent of tanezumab, DHODHi, and AuQCs also actively imposing therapeutic mitigation to some extent.

Oxidative stress, which produces reactive oxygen species (ROS), is widely implicated in the pathogenesis of IC/BPS(21). Ultrasmall-sized metal nanoclusters, especially protein-protected ones, usually possess stronger biocatalytic activities than large NPs, thereby scavenging ROS and mitigating inflammation(22). To scrutinize the anti-inflammation contribution of AuQCs in NADC, we conducted a series of biochemical assays and electron spin resonance (ESR) spectroscopy (Supplementary Fig. 22). AuQCs display potent radical scavenging against ABTS^•+^ that even outperforms Trolox, alongside the enzyme-mimicking activities of catalase (CAT), superoxide dismutase (SOD), and glutathione peroxidase (GPx) involved in multiple ROS conversion processes. These antioxidation properties are also identified in NADC, but neither vidofludimus nor anti-NGF. ESR spectra confirmed the scavenging of hydroxyl radicals (·OH) and superoxide anion (O_2_ ^·-^) by the AuQCs constituent in NADC. Further, the enzyme inhibitory effect on DHODH by vidofludimus was retained in NADC (Supplementary Fig. 23).

### NADC as a therapeutic agent to treat chronic interstitial cystitis in rats

Next, we estimated the in vivo therapeutic efficacy of NADC in a standard chronic IC rat model established by repeated intraperitoneal (i.p.) CYP induction (Fig. 5a). During the entire therapeutic process, we applied the Von Frey rodent test with a calibrated filament to determine the mechanical sensitivity threshold from lower abdomen withdrawals in response to a punctate stimulus (Fig. 5b). Meanwhile, we carried out time-course bladder urodynamics to meticulously track urinary voiding functions (Fig. 5c). After IC onset by repetitive low-dose CYP stimulations, the mechanical withdrawal threshold in the IC rat models was reduced and consistently maintained at a low level with a prominent reflex over the entire period, indicative of persistent, severe lower abdominal pain (Fig. 5d). Nonetheless, intravesical therapy with NADC effectively attenuated pain symptoms and almost reversed the mechanical allodynia threshold of IC to normal levels, with individual NADC components exerting partial yet significant efficacy (Fig. 5d). FDA-approved intravesical instillation of DMSO and hyaluronic acid (HA) has been recommended by the American Urological Association (AUA) guidelines as the clinical standard of care for IC.

**Figure 5.**
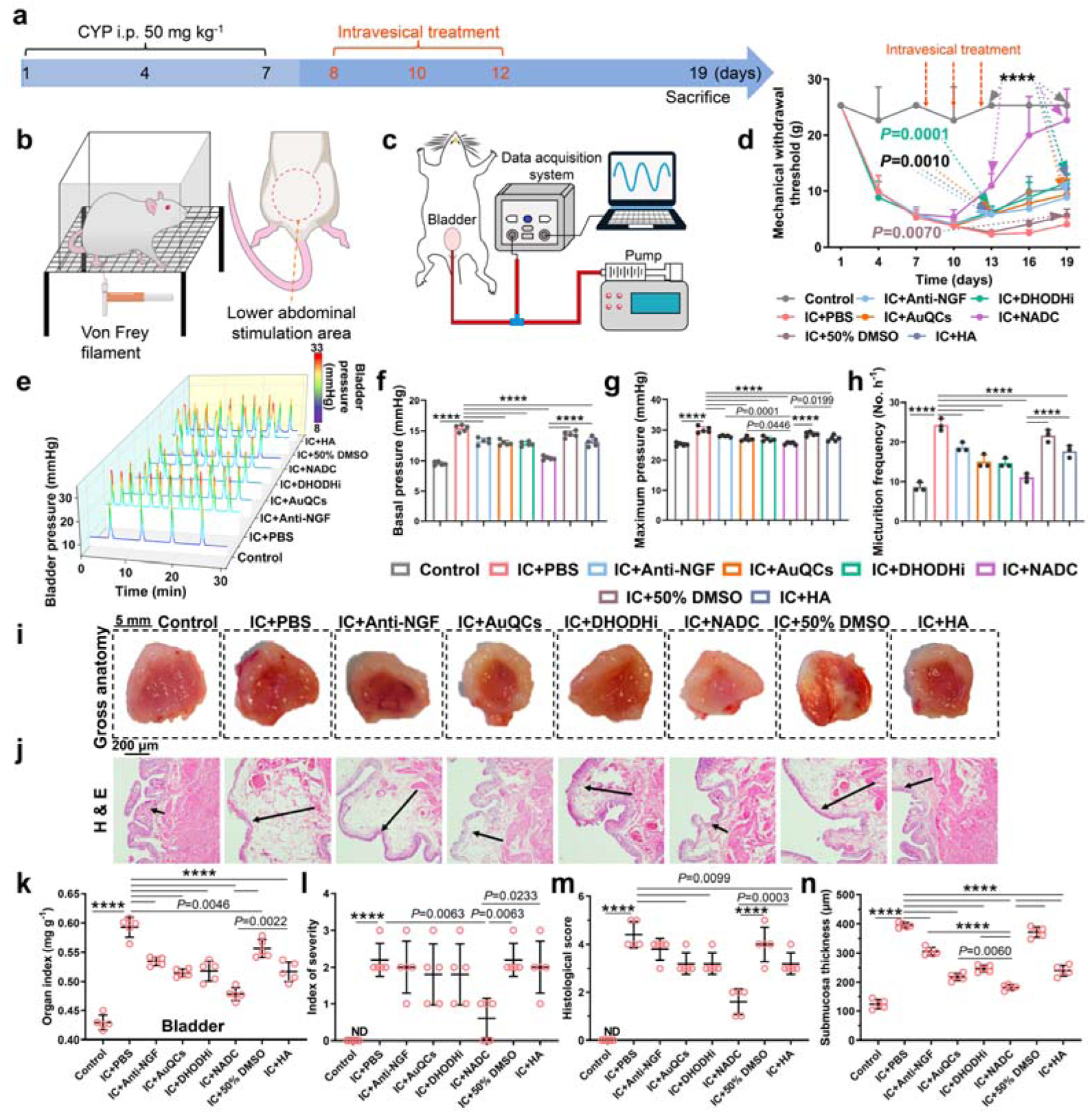
NADC as a therapeutic agent to treat chronic IC in rats. (a) Flowchart showing the chronic IC induction and therapeutic regimens in rats. Schematic illustrations of the procedures for (b) the Von Frey filament test and (c) urodynamics. (d) Changes in the mechanical allodynia threshold over time in chronic IC rat models receiving indicated interventions. All statistical analyses between groups in *P* values were compared with the IC+PBS group (n=5 animals per group). (e) Real-time bladder urodynamic curves of rats with chronic IC in different groups. (f) The basal pressure, (g) maximum pressure, and (h) micturition frequency of bladder urodynamics in different rat groups (n=3 to 5 animals per group). (i) Gross anatomy of rat bladders harvested at the therapeutic endpoint of intravesical infusion. NADC significantly reduced symptoms of IC-induced congestion, bleeding, and edema. (j) Histopathology of resected bladders in indicated treatment groups by H&E staining. The length of the arrow indicates the bladder’s mucosal edema zone. (k) Bladder organ index defined by the bladder weight/body weight ratio, (l) index of severity, (m) histological scores, and (n) submucosa thickness (n=5). Statistical significance levels: \*\*\*\**P* ≤ 0.0001.

NADC outperformed intravesical DMSO and HA in pain relief and therapeutic efficacy. Furthermore, urodynamic assessment informed markedly heightened basal and maximum intravesical pressures alongside increased micturition frequency (Fig. 5e-h), mirroring the clinical presentation of IC patients who experience recurrent urination and urgency. After intravesical therapy, the abnormal urinary symptoms were all rehabilitated partly by tanezumab, DHODHi, AuQCs, and HA but restored to healthy levels with uninfluenced voiding by NADC.

In addition to soothing IC symptoms, NADC also improved bladder macroscopic manifestations through anatomical examination of gross pathology, significantly diminishing IC-triggered congestion, bleeding, and edema (Fig. 5i). Accordingly, the bladder organ index elevated by IC was mitigated by NADC and each constituent (Fig. 5j). Compared to the intact bladder anatomy with high integrity in the normal bladder, inflamed bladder lesions in IC rats featured a disrupted structure with urothelium denudation, submucosal edema, inflammatory infiltration, lamina propria hemorrhage, and granulation, which were adequately attenuated by NADC (Fig. 5k). Further histopathological analyses unveiled palliated outcomes on the index of severity, histological scores, and submucosal thickness by NADC, which were all raised in IC (Fig. 5l-n). The results confirmed intravesical NADC’s effectiveness in restoring cystitis-associated pathological damage to bladder tissues. In addition, while both anti-NGF-DHODHi and anti-NGF-AuQCs conjugates showed appreciable anti-inflammatory responses and symptom attenuations in vivo, the overall therapeutic outcome was inferior to that of NADC (Supplementary Fig. 24). The therapeutic efficacy of intravesical NADC was highly durable with no recurrence for over a month (Supplementary Fig. 25).

### NADC as a therapeutic agent to treat acute interstitial cystitis (AIC) in rats

Although IC is typically characterized as a persistent and chronic condition, part of patients may experience acute exacerbations of flare-ups with intensified symptoms. To evaluate the NADC efficacy against the exacerbated condition(23), we constructed an AIC rat model induced with a high dose of CYP (Fig. 6a). As measured by the Von Frey test, the 50% mechanical withdrawal threshold of AIC rats was rapidly curtailed within 4 days, reflecting severe hyperalgesia (Fig. 6b). Meanwhile, the body weight of AIC rats sharply dropped in short periods (Fig. 6c). To explore the influence of AIC on locomotor activities and anxiety-like behaviors of rats, we performed the open field test (OFT) and tracked the exploration footage using an overhead camera (Fig. 6d). Healthy rats moved freely in all zones of the arena with abundant footage, but rats suffering from AIC-induced pain were crouched with deficient total traveled distance, low max velocity, and restricted locomotor areas (Fig. 6e-h). The willingness, capability, and rate of motor activities were affected by the visceral pain from AIC. By comparison, intravesical therapy of NADC substantially recovered the mechanical allodynia, body weight loss, and diminished locomotor parameters, with individual NADC constituents also being moderately effective in cystitis remission (Fig. 6). Additionally, the urinary function of AIC rats was verified by urodynamic testing (Fig. 6i). Notably, the aggravated basal and maximum bladder pressures and urination frequency in AIC rats were significantly alleviated with intravesical instillation, with the most pronounced therapeutic effect observed for NADC (Fig. 6j-l). Gross pathology unveiled more hemorrhagic lesions and more prominent edema in the bladder mucosa of AIC rats (Supplementary Fig. 26a) than in chronic IC (Fig. 5i), which were effectively mitigated by intravesical therapy. Correspondingly, the bladder submucosal edematous zone markedly thickened and hemorrhaged by AIC was microscopically visualized to be improved by the NADC intravesical intervention (Supplementary Fig. 26b). Aligned with earlier results on chronic IC, the elevated organ index, severity, and histological score in AIC were suppressed (Supplementary Fig. 26c-e).

**Figure 6.**
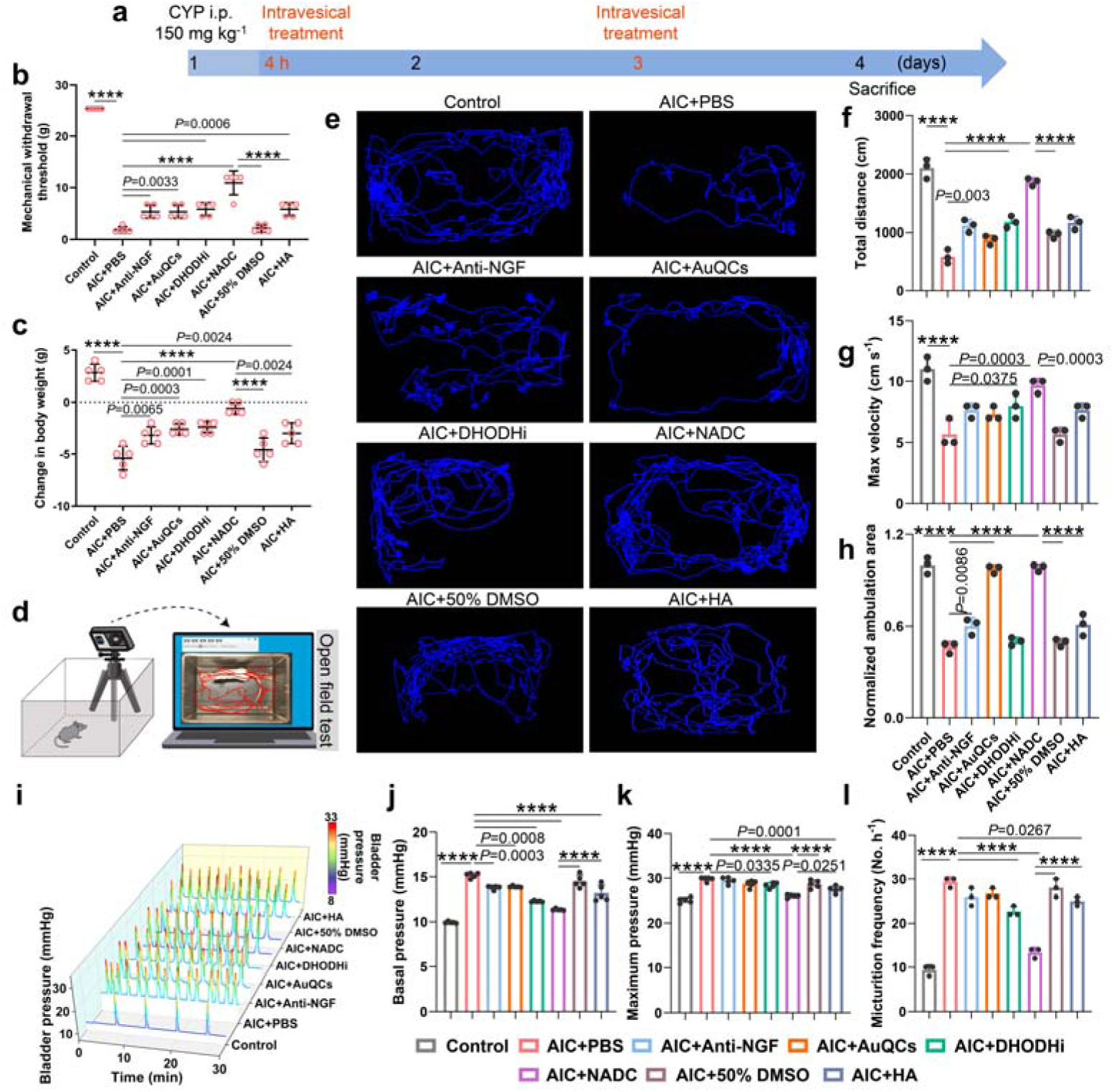
NADC as a therapeutic agent to treat AIC in rats. (a) Flowchart showing the AIC induction and therapeutic regimens in rats. (b) Mechanical withdrawal thresholds of rats receiving indicated intravesical interventions (n=5 animals per group). (c) Body weight changes of rats after treatment at the study endpoint (n=5 animals per group). (d) Schematic illustration showing the setup of open field tests (OFT) in the study. (e) Travel trajectories of the exploration footage of rats with AIC receiving various treatments. (f) The corresponding total traveled distance, (g) maximum velocity, and (h) normalized restricted locomotor area of rats from different groups (n=3). (i) Real-time bladder urodynamic curves of rats with AIC in different groups. (j) The basal pressure, (k) maximum pressure, and (l) micturition frequency of bladder urodynamics in different rat groups (n =3 to 5 animals per group). Statistical significance levels: \*\*\*\**P* ≤ 0.0001.

### NADC as prophylactic therapy for IC in rats

Gleaned from anti-microbial agents for prophylactic therapy of bacteria-associated cystitis, we aim to study the revelation of intravesical NADC as non-surgical and minimally invasive prophylaxis for IC. Before the high-dose CYP induction to recapitulate exacerbated cystitis in rats on the 6^th^ day, we perfused the bladder with intermittent prophylactic NADC at pre-defined dosing (Fig. 7a). Coinciding with its therapeutic effects in chronic IC and acute cystitis, pre-dosed NADC exhibited appreciable preventative efficacy on the mechanical pain threshold, body weight changes, urodynamics, bladder pressures, and micturition frequency deteriorated from cystitis onset (Fig. 7b,c, Supplementary Fig. 27a,b). Specifically, healthy rats urinated 4 times in 30 min, whereas rats with acute cystitis urinated at a higher emiction frequency of 15 times. Following NADC prophylaxis, the frequency was reduced to 5 times, compared to 14 and 12 times by prophylaxis of 50% DMSO and HA, respectively (Supplementary Fig. 27b). Macro- and microscopic histopathological analysis of the bladder further verified the prevention of cystitis-related bladder injuries by NADC prophylaxis, with insignificantly altered organ index, severity, and histological score concerning healthy rats (Fig. 7d, Supplementary Fig. 27c-g). Notably, each NADC unit, i.e., vidofludimus, tanezumab, and AuQCs, was competent in counteracting acute cystitis induction.

**Figure 7.**
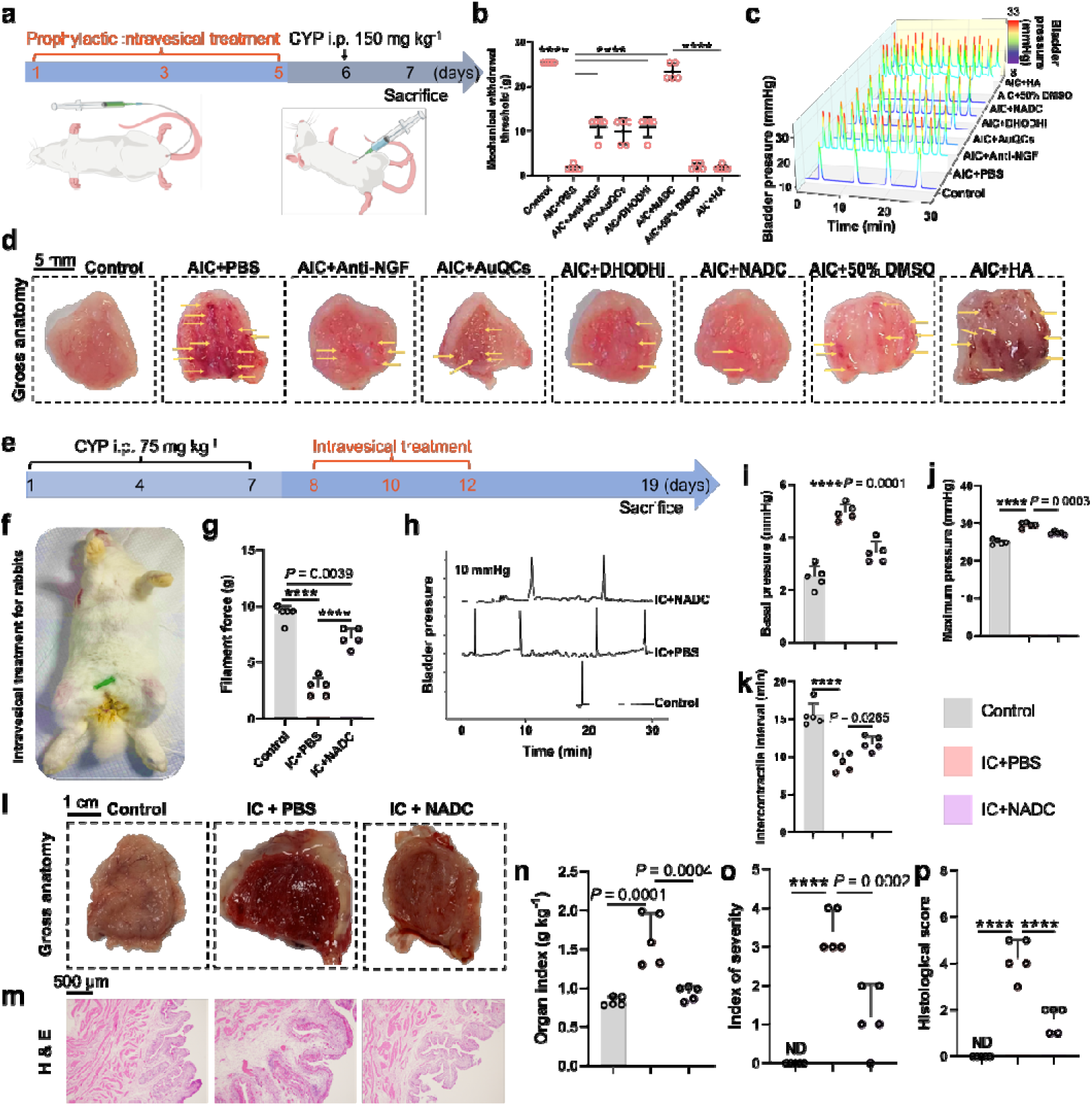
Evaluation of intravesical NADC as prophylaxis in rats and intervention in rabbits for IC. (a) Flowchart showing the therapeutic regimens of rats followed by AIC induction. (b) Mechanical withdrawal thresholds of rats receiving indicated intravesical prophylaxis before AIC induction (n=5). (c) Real-time bladder urodynamic curves of AIC rats with prophylactic treatment. (d) Gross anatomy of bladders harvested from rats receiving intravesical prophylaxis before AIC induction. Yellow arrows indicate bleeding lesions in bladder tissues. (e) Flowchart showing IC induction and therapeutic regimens in rabbits (n=5). (f) A photograph showing intravesical procedures in rabbits. (g) Mechanical withdrawal thresholds of rabbits from various groups measured by von Frey filaments. (h) Real-time bladder urodynamic curves, (i) basal pressure, (j) maximum pressure, and (k) intercontractile interval of rabbits from various groups. (l) Gross anatomy of rabbit bladders harvested at the therapeutic endpoint. (m) Bladder histopathology, (n) bladder organ index, (o) index of severity, and (p) histological scores of rabbits in indicated groups. Statistical significance levels: \*\*\*\**P* ≤ 0.0001.

### Therapeutic efficacy of NADC in non-rodent rabbit models

To further strengthen preclinical validity with multispecies advocacy, we evaluated the therapeutic efficacy in rabbit IC models, as rabbits possess a larger bladder volume, stratified urothelium, and a glycosaminoglycan layer composition more akin to humans. IC was induced in rabbits via i.p. injection of CYP, and intravesical NADC was administered by urethral catheterization (Fig. 7e,f). As the hallmark of neuropathic pain, the mechanical allodynia of rabbits was significantly alleviated by NADC alongside improved bladder urodynamics and pressure parameters (Fig. 7g-k). Like in rodents, NADC mitigated the bladder manifestation defects of hemorrhage and edema from IC with microscopically ameliorated histopathology (Fig. 7l-p).

### Coordinative molecular mechanisms of NADC

Although the mechanism of action for NADC is straightforward via dual selective inhibition of DHODH and NGF, the downstream and synergistic pathways remain underexplored. In addition, AuQCs manifested theranostic features with comparable therapeutic and prophylactic efficacies (Fig. 5-7). To inspect the underlying molecular mechanisms of the NADC complex against IC, we profiled transcriptomes of bladders dissected from IC rats treated with PBS or NADC by RNA sequencing (RNA-seq) with reference to healthy controls. The high-dimensionality RNA-seq data were dimensionally reduced by principal component analysis (PCA), distinctly separating the IC group by differentially expressed genes (DEGs) from the converged healthy control and IC rats receiving intravesical NADC treatment (Fig. 8a). The number of DEGs altered by IC among all genes increased in both up- and down-regulated directions, which were extensively restored by intravesical NADC instillation (Fig. 8b). In particular, the plentiful DEGs downregulated by IC were overturned into augmented expressions by NADC. We then mapped the DEGs for enrichment analyses of gene ontology (GO) and Kyoto Encyclopedia of Genes and Genomes (KEGG) (Fig. 8c, d). GO informatics highlighted enriched slim annotations of IC for response to external stimulus, generation of ROS, response to chemical, and ROS-modulated enzyme activities, while NADC assimilated GO terms of cytokine and cytokine receptor activities, cytokine production, growth factor receptor binding and activity, inflammatory response, immune response, cellular response to TNF, IL-6 production, and ROS metabolism (Fig. 8d). Next, miscellaneous KEGG cascades noted for cytokine regulation, immune modulation, neuroactive ligand-receptor interaction, glycolysis/gluconeogenesis, and inflammation were enriched (Fig. 8c), comprehensively covering the combinational effects of NGF and DHODH dual inhibitions and immune-modulating AuQCs. Compared to their larger-sized counterparts, ultrasmall nanoclusters AuQCs have been widely reported to scavenge ROS and attenuate inflammation with enhanced catalytic activities(24). Importantly, we highlighted coordinative admitted inflammation signaling pathways of IL-17, NF-κB, and JAK-STAT that were dysregulated by IC but effectively mitigated by NADC with opposite regulations (Fig. 8e). Key quality-control passed gene panels of these inflammation pathogenic pathways in the bladder transcriptome significantly altered by IC were rescued by NADC and almost restored to control levels (Fig. 8f and Supplementary Fig. 28), suggestive of IC relief. Protein-protein interaction network analysis modeled the complex pathways and biological functions, which distinctly disclosed the interconnected signaling pathways shared with interacting nodes (Supplementary Fig. 29, 30).

**Figure 8.**
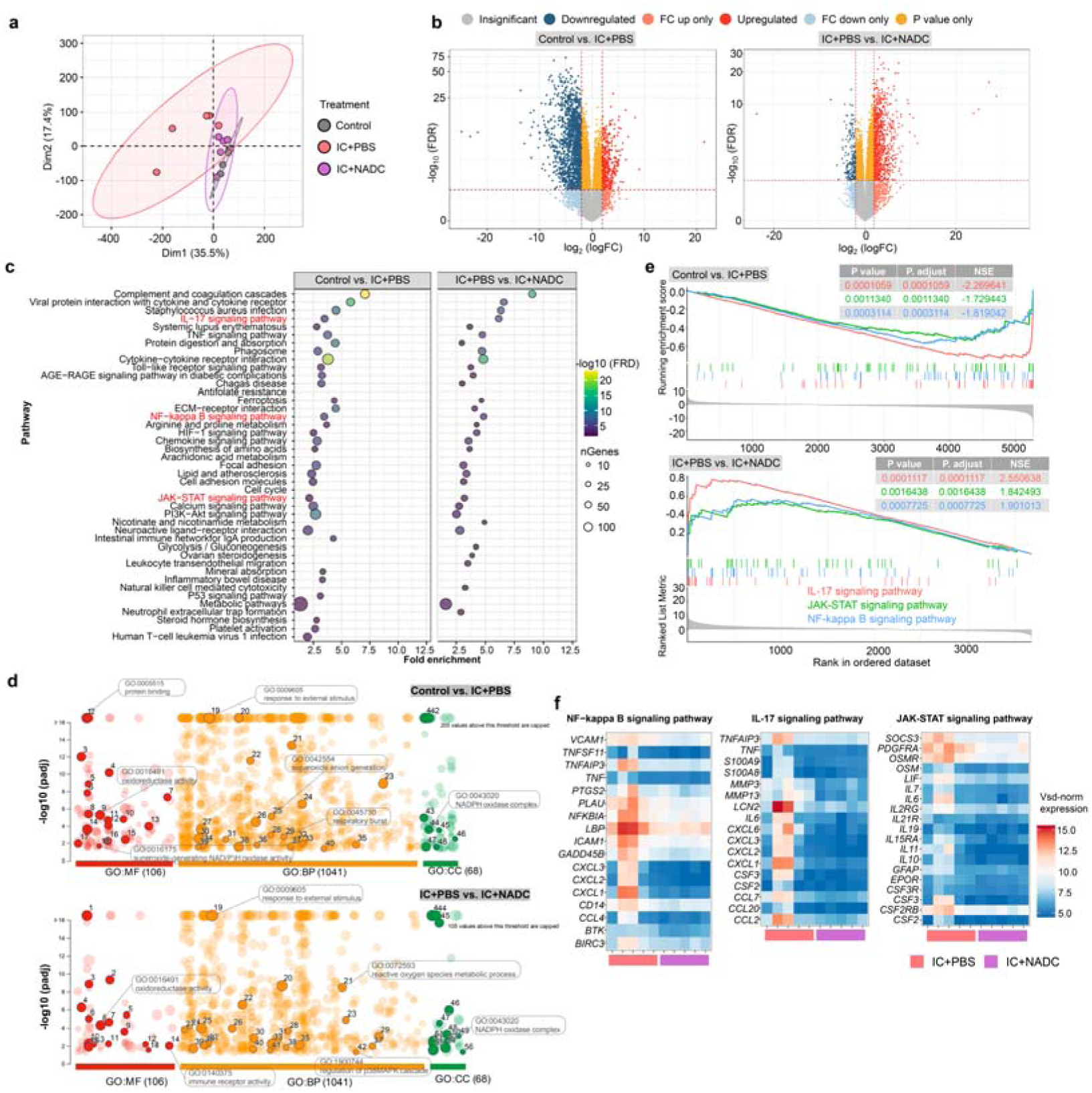
Coordinative molecular mechanisms of NADC. (a) PCA of the RNA-seq raw data indicates that IC is the primary factor distinguishing sample populations (n=5). Dim1 and Dim2 represent the first and second dimensions, respectively. (b) Volcano plots illustrate the pattern distribution of DEGs between compared groups by RNA-seq. IC alters an elevated number of DEGs in upregulated and downregulated categories but is substantially reverted following intravesical NADC instillation. (c) KEGG pathways derived from DEGs post-NADC treatment with low and high q-values denoted in blue and green. The circle size reflects enriched gene counts. (d) GO enrichment mapping specifically relevant to IC occurrence and follow-up intravesical NADC. (e) GSEA of intravesical NADC therapy predominantly correlated with the NF-κB, IL-17, and JAK-STAT signaling pathways. (f) Heatmaps of significantly altered key effector genes in the NF-κB, IL-17, and JAK-STAT pathways in IC rat bladder tissues with or without NADC treatment. Expression values from high to low are displayed in color gradients ranging from red to blue.

To complement transcriptional analysis, we further interrogated protein expressions of key effectors in the pinpointed pathways by immunoblotting. In consistency with IF results (Fig. 3e,f), NGF was overly expressed with CYP induction in IC bladder tissues of rats to provide a viable precision target for NADC (Supplementary Fig. 31). Interestingly, none of the agents suppressed the aberrant NGF expression, including NGF-targeting tanezumab and NADC, corroborating a sustained targeting efficiency. In contrast, the phosphorylation of the transmembrane NGF catalytic receptor and pain sensory mediator, tropomyosin receptor kinase A (TrkA), was incremented in IC but inhibited by tanezumab and NADC, with total TrkA remaining identical in healthy and IC rats and independent of therapeutic interventions. Next, we probed major pro-inflammatory cytokines and observed potent inhibition of IL-17A, IL-6, and IL-1β by DHODHi and TNF-α by AuQCs in NADC, with negligible effects from anti-NGF (Supplementary Fig. 31). As the upstream regulator, IL-17A is a known target of vidofludimus that impairs NF-κB and JAK-STAT pathways(25).

After that, we comprehensively screened pivotal mediators in NF-κB and JAK-STAT pathways at protein levels (Supplementary Fig. 32). Defectively elevated phosphorylation of the two IκB kinase (IKK) catalytic subunits, IKKα and IKKβ, and RelA/p65 resulted in activated canonical NF-κB in IC models, which was adequately inhibited by NADC to limit degradation of the NF-κB inhibitor IκBα(26), yet with minimal influence on the total protein expressions. Likewise, IC-deregulated JAK-STAT members broadly involved in cytokine signaling, including JAK1, STAT1, and STAT2, were appreciably attenuated by NADC with potent synergy, which concurrently prohibited phosphorylation and expression.

### Alleviation of IC through bladder microbiota modulation by NADC

Although the bladder microbiome is less diverse and traditionally overlooked, emerging clinical evidence has established a plausible correlation between urinary microbiomes and IC/BPS(27). To investigate whether NADC mitigated IC through leveraging bladder microbiota, we isolated bacterial DNA from rat bladder tissues and ran 16S ribosomal RNA (rRNA) gene amplicon sequencing. Using non-metric multidimensional scaling (NMDS) analysis of Bray-Curtis dissimilarity distances, we identified a distinct bladder microbiota in IC rats significantly differing from all the other treatment groups (*P*<0.05) that shared high similarities with the healthy control (Fig. 9a). Comparatively, DHODHi monotherapy induced a more extensive segregation of microbiome alteration. At genus levels, we found significant divergence of the IC microbiome from that of healthy rats in the composition and abundance (permutational multivariate ANOVA testing, Pr(>F)=0.02), with overrepresented genera like *Escherichia*, *Shigella*, *Enterococcus*, *Staphylococcus*, and *Enterobacter* and underrepresented genera like *Acinetobacter*, *Burkholderia*, *Caballeronia*, and *Paraburkholderia* (Fig. 9b). The number of observed operational taxonomic units (OTUs), Chao1, and ACE indices in the alpha-diversity metrics were significantly reduced in IC and anti-NGF treatment groups but were rescued to normal levels with treatment of AuQCs, DHODHi, or NADC (Fig. 9c). Further taxonomic analysis of microbial community compositions of OTUs in bladder tissues validated the conservation of core genera of *Rhodococcus* among all groups, along with decreased abundance in high-prevalence *Acinetobacter* and increased abundance in less-prevalence *Escherichia*-*Shigella* in IC (Fig. 9d). With linear discriminant analysis (LDA) effect size (LEfSe), cladograms highlighting key bacterial clades portrayed spatial phylogenetic correlations of bacterial lineages in taxonomic hierarchies (Fig. 9e). Noticeably, the bacterial taxa at phylum, class, order, family, and genus levels were all significantly varied by IC to some extents (LDA scores ≥2.0) yet were inversely regulated in a similar clade landscape by intravesical NADC therapy. The relative abundance of *Gammaproteobacteria* (class) and *JG30-KF-CM45* (family to species) with LDA scores ≥3.0 by LEfSe were significantly higher in IC, contributing to pathogenesis (Fig. 9f). In contrast, *Actinobacteriota* (phylum and class), *Nocardiceae* (family), *Rhodococcus* (genus and species), and *Corynebacteriales* (order) were enriched following NADC therapy, which depreciated *Gammaproteobacteria* (class) along with other virulent populations of *Enterobacteriaceae* (family), *Escherichia*-*Shigella* (genus to species), *Bacilli* (class), and *faecalis* (*Enterococcus* species) (Fig. 9f). Therefore, the bladder microbiome remodeling by NADC plays an indispensable role in IC mitigation.

**Figure 9.**
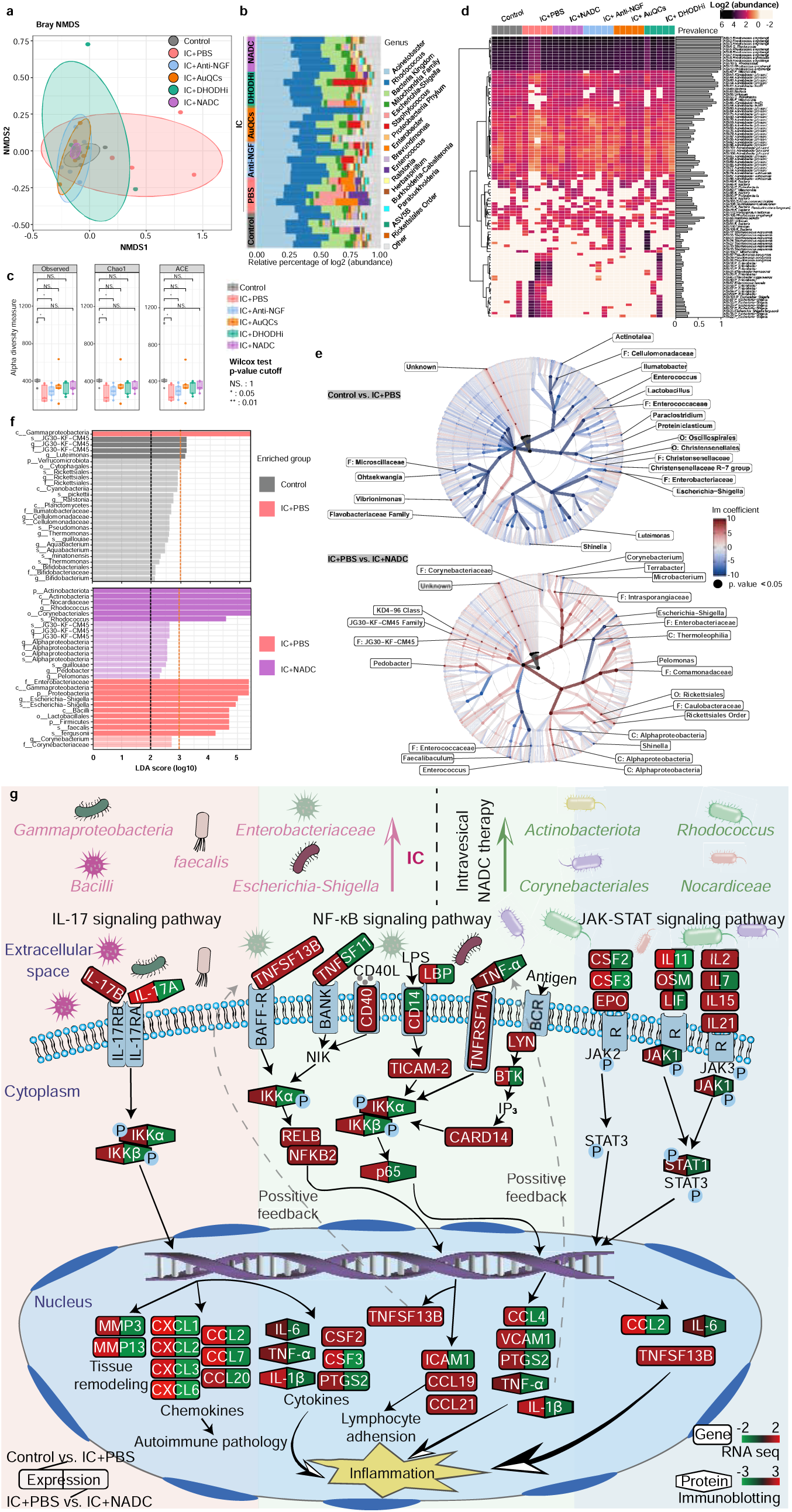
Alleviation of IC through bladder microbiota modulation by NADC. (a) NMDS analysis of Bray-Curtis distances across bladder tissue samples (n=5 animals per group) revealed distinct microbiota compositions in IC-afflicted rats, markedly divergent from other treatment cohorts. Each point symbolizes the bladder microbiota of an individual rat, with NMDS data highlighting significant microbial distinctions. (b) A bar graph illustrated the genus-level relative abundance of gut microbiota. (c) Variations in alpha diversity among different groups were presented, including observed counts, species richness (Chao1), ACE, Shannon, and Simpson indices. Statistical significance is denoted by * (p < 0.05) and ** (p < 0.001), as determined by Wilcoxon rank sum tests. (d) A heatmap displayed the relative abundance (above a 0.001 cutoff) and prevalence (above a 75% cutoff) of prokaryotic OTUs in the examined bladder tissue samples. (e) A clade diagram illustrated the phylogenetic distribution of microbial lineages pertinent to these two groups, with each smaller circle representing a taxonomic level’s taxon and the diameter corresponding to relative abundance (layers from inner to outer: genus, family, order, class, and phylum). (f) Statistical analyses employing the LDA effect size method revealed significant taxa between groups, identified by an alpha value of 0.05 and an LDA (log10) absolute value greater than 2.0. The Gammaproteobacteria class and the JG30-KF-CM45 lineage (from family to species) were significantly more abundant in IC, as indicated by LDA scores ≥3.0 using LEfSe, implicating them in disease pathogenesis. (g) Mechanistic synergy of intravesical NADC therapy against IC by coordinating NF-κB, IL-17, and JAK-STAT signal transduction pathways. Effectors in mRNA and protein expressions are mapped from immunoblotting and RNA-seq. Left half: IC before treatment; right half: IC after NADC treatment.

### Safety profiles of NADC

As evidenced by earlier transmembrane and imaging studies (Fig. 4k-p and Supplementary Fig. 15-19), catheter-directed intravesical NADC was refrained from penetrating bladder tissues and entering the systemic circulation, theoretically with minimized systemic toxicity. To verify such claims, we scrutinized toxicity profiles of intravesical NADC and individual drug payloads by comprehensively screening panels of clinical chemistry and hematology. Electrolyte balances of bicarbonate (CO_2_), K^+^, Na^+^, Cl^-^, and Ca^2+^ were well maintained (Supplementary Fig.33). Additionally, indicators of liver and kidney functions, such as alanine transaminase (ALT), aspartate transaminase (AST), alkaline phosphatase (ALP), lactate dehydrogenase (LDH), blood urea (UREA), and creatinine (CREA), all fell within reference ranges without significant changes from the control. Blood proteins of albumin (ALB), globulin (GLB), and total protein (TP) stayed normal. The complete blood count (CBC) covering red blood cells (RBC), white blood cells (WBC), and platelets (PLT) in all groups lay in biologically non-significant reference ranges, despite a minor elevation of basophils (BAS) observed with NADC treatment (Supplementary Fig. 34).

## Discussion

The psychological impact of IC is pervasive, long-lasting, and debilitating, with frequent urination and pain negatively interfering with social intimacy, work, and daily activities. Nonetheless, practical, accessible, and affordable therapy for IC is somewhat limited in clinical practice. As a relatively emerging therapeutic modality, ADC combines the specificity of monoclonal antibodies and the cytotoxic potency of drug payloads, necessitating precise delivery and selective killing to tackle otherwise unsolvable clinical challenges. While there are fifteen ADCs approved by the US Food and Drug Administration (including recent ones like Tivdak and Elahere) and more in clinical phases poised to enter the market(28, 29), indications are predominantly limited to hematologic malignancies and solid tumors, with autoimmune conditions primarily overlooked. Cytotoxic payloads of ADCs are unsuitable for treating IC, where the therapeutic objective is to direct immune modulation or suppression rather than causing cell death. Besides, all current ADC drugs lack a companion imaging modality to inform their pharmacokinetics and drug distribution patterns. As an imaging facilitator, AuQCs afford intrinsic NIRF with a significant Stokes shift arising from band gap discretization and optical quantum confinement, aside from the high atomic number (*Z*=79) and *K*-edge (80.7 keV) of Au as superior CT contrast(30). Besides whole-body in vivo imaging (Fig. 4), the NADC is also liable for microscopical visualization of drug distribution patterns in high resolution. Interestingly, despite the weaker signal, the distribution of AuQCs without antibody-mediated targeting was also confined in regions with NGF expression, probably by passive targeting (Fig. 4e-f).

Therefore, intravesical NADC polytherapy enables visualizable precision medicine for IC, combining catheter-guided direct dosing of high drug concentrations, NGF-specific targeting, selective DHODH modulation, and theranostic AuQCs. Compared to systemic administration, intravesical instillation delivers a fixed dose relatively independent of body weight, raising local drug concentrations in the dysfunctional bladder more rapidly. The stringently controlled infusion rate can reduce infusion-related reactions and hypersensitivity. A non-cleavable covalent linkage of stable amide and thioether bonds in NADC prevents the premature release of active DHODHi and AuQCs and warrants cytosolic efficacy after antibody degradation. It has been reported that ADC with a thioether covalent linker is more effective in proliferating cancer cells with higher activities than a disulfide cleavable counterpart(31). Therefore, such a structural design can improve in vivo stability, therapeutic window, and pharmacokinetics of payloads. In the study, the multifold precision of NADC by intravesical drug dosing spared normal bladder tissues with low urothelial NGF expression and immunologically modulated inflammatory cells with non-cytotoxic payloads of vidofludimus and AuQCs. Although the bladder mucosal urothelium is a multifaceted tight-junction barrier known as one of the most impermeable membranes of the body, patients with IC conditions may experience increased bladder permeabilities(32). As verified by whole-body imaging and Ussing chamber assays (Fig. 4 and Supplementary Fig. 15-19), the limited bladder permeability of NADC serves as a prerequisite of locoregional intravesical therapy without entering systemic circulation or compromising urothelial integrity and detrusor muscles. Consequently, the direct relief of inflamed bladder linings by intravesical NADC can bypass drug overdose and improve patient tolerance, resulting in minimized off-target toxicity (Supplementary Fig. 33-34).

In the current IC management, AUA clinical guidelines recommend systemic treatment as the first and second line, but the overall effectiveness is limited. Intravesical DMSO has long been approved by the FDA as the standard third-line treatment for IC, and more recently, HA that repairs damaged GAG barriers has been clinically recommended. Our study showed that the two guideline-endorsed treatments underperformed NADC in chronic, acute, and prophylactic rat models (Fig. 5-7) and rabbit models (Fig. 7e-p). Together with the imaging modalities, NADC might be proposed for potential guideline revisions. Meanwhile, the drug antibody ratios (DAR) of AuQCs and DHODHi fall around optimal DAR values between two and four for most ADCs with balanced safety and efficacy(33), ensuring appreciable therapeutic efficacy on the mechanical threshold and micturition function against chronic IC and AIC (Fig. 5-6). Gleaned from anti-microbial agents for the prophylactic therapy of bacteria-associated cystitis, we also studied the revelation of intravesical NADC as non-surgical and minimally invasive prophylaxis for IC. The intravesical prophylactic therapy with NADC prevented pain symptoms and bladder dysfunction from acute cystitis to limit its occurrence and recurrence (Fig. 7). Following receptor-mediated internalization to release AuQCs and vidofludimus payloads, NADC concurrently inhibits multiple inflammatory pathways such as NF-κB, JAK-STAT, IL-17, and TNF as combinatorial precision therapy (Fig. 8c-f), which was well-documented with clinical success(34, 35). As a promising transformative therapeutic, NADC also provides alternative mechanisms toward IC alleviation by reshaping the bladder microbiome in composition and abundance (Fig. 9). The altered bacteria with endosymbiosis are widely involved in bioremediation, biosynthesis, and regulation of inflammation(36, 37).

In summary, we report for the first time a theranostic intravesical NADC with immunomodulating drug payloads in non-cleavable covalent linkage, serving as a unified multifaceted precision agent against chronic, acute, and prophylactic IC symptoms through catheter guidance. Besides the ability to be tracked by imaging, the NADC simultaneously regulates multiple perturbed inflammation-related signaling pathways and remodels the host bladder microbiota. Our findings here provide new insights into the development and in vivo visualization of theranostic ADCs, which may be further extended to a broader category of ADCs, peptide-drug conjugates, aptamer-drug conjugates, and other relevant conjugates to treat autoimmune diseases in general.

## Methods

### Synthesis of NADC

AuQCs with optimal NIR fluorescence were synthesized as previously reported(19). Briefly, 40 mg mL^-1^ α-LA and 10 mM HAuCl_4_ were mixed at equivalent volumes. After that, 1 M NaOH at 15% (v/v) was added to the mixture and sufficiently vortexed until the precipitates formed from the prior step were completely dissolved. The reaction then proceeded with stirring at 40 °C for 20 h and was dialyzed against pH 7.4 PBS at 10 kDa molecular weight cutoff for 2 days. The dialysis was sealed from light with fresh buffers substituted every 8 h. The concentration of AuQCs was quantitatively measured by nanoparticle tracking with a NanoSight NS500 system (Malvern Panalytical).

To prepare the NADC, we first modified tanezumab with ultrasmall AuQCs(38). Typically, 20 μg tanezumab in 100 μL PBS reacted with 26.6 μM Traut’s reagent at room temperature for 1 h to modify primary amines into reactive sulfhydryl groups. In parallel, 30 μM water-soluble sulfo-SMCC was added to 6 μM AuQCs in PBS and maintained for 30 min at room temperature to drive amide bond formation between NHS esters and primary amines of AuQCs. After washing with PBS by 3 kDa ultrafiltration (Millipore), the sulfhydryl-activated tanezumab reacted with maleimide-modified AuQCs by the hetero-bifunctional sulfo-SMCC for 2 h in the dark at 4 °C, resulting in the stable formation of thioether bonds (Fig. 1a). To remove unreacted AuQCs, we extensively washed the modified complex with PBS via 100 kDa ultrafiltration (Millipore). Last, the remaining unreacted amines in the complex were covalently conjugated to the benzoic acid of vidofludimus to acquire NADC. 20 μM equimolar NHS and EDC were appended into 25 μM vidofludimus and kept for 15 min at room temperature(39). Afterward, activated vidofludimus was added to AuQCs-modified tanezumab and incubated for 2 hours at room temperature from light. Excessive vidofludimus small molecules in the reaction were eliminated by 3 kDa ultracentrifugation.

### Single-cell RNA-seq analysis

The scRNA-seq data for IC were obtained from the GEO database (GSE175526) and processed using the Seurat package. The quality control criteria applied included: (1) a minimum of 300 genes detected per cell and (2) less than 15% mitochondrial gene content. After quality control, batch effects were corrected, and the samples were integrated using the Harmony package. Cell clustering was conducted with the FindClusters function (resolution = 0.2) and visualized using the t-SNE method. Clusters were annotated based on marker gene expression in Supplementary Table 1. We conducted enrichment analysis using the clusterProfiler package (version 4.12.3) to evaluate GO and KEGG pathways for all marker genes from clusters 0 and 9. For GO enrichment analysis, we identified significant GO terms specific to each cluster by excluding overlapping terms between the two clusters. We then selected the top 10 terms for Biological Process (BP), 10 terms for Cellular Component (CC), and 10 terms for Molecular Function (MF), based on adjusted p-values. In cluster 9, 8 genes of particular interest were highlighted in red. We ranked the pathways based on adjusted p-values for KEGG analysis and selected the top 20 pathways with the lowest p-values for each cluster. All visualizations were created using the ggplot2 package version 3.5.1 to present the enrichment results.

### Biolayer interferometry assays

To confirm the binding affinity between NADC and NGF after conjugation, we performed label-free BLI analysis using the Octet R8 system (Sartorius)(40). First, the anti-human IgG Fc capture (AHC) biosensor (Sartorius) was conditioned in the detergent-containing Kinetic Buffer (KB, PBS with 0.02% v/v Tween-20) for 60 s. Next, 75 μg mL^-1^ NADC or tanezumab on an equivalent antibody basis was immobilized onto the AHC biosensor surface for 60 s. Gradient NGF concentrations from 31 to 500 nM were bound to the modified biosensor during the association process, followed by dissociation in KB. The equilibrium dissociation constant (K_D_) was calculated from the binding curves in the Octet Analysis Studio software (Sartorius).

### In vivo spectral CT imaging

Dual-energy CT imaging was performed using a GE Revolution CT scanner with GSI Xtream (GE Healthcare). Monochromatic images were captured in a 40–140 keV fast-switching energy range and processed in the GSI Viewer software (GE Healthcare) to reconstruct composite images following noise reduction(41, 42). Before imaging, SD rats were anesthetized with 7% chloral hydrate (350 mg kg^-1^) by intraperitoneal injection and immobilized in the supine position. Intravesical administration of 500 μL PBS, AuQCs (15 μM), or NADC (5 μM) was initiated via the urethra using a 20-gauge indwelling catheter needle. CT scans were acquired in the axial mode, and three-dimensional multiplanar reconstruction images were generated. CT numerical values in Hounsfield Units (HU) and corresponding standard deviations (SD) were retrieved from the average of three individual ROIs(43).

### In vivo therapy in rat and rabbit IC models using intravesical NADC

SD rats were supplied by the Institute of Experimental Animals at Sun Yat-sen University. The therapeutic efficacy of intravesical NADC was evaluated in chronic, acute, and prophylactic IC rat models in eight groups: healthy control, IC + PBS, IC + anti-NGF, IC + AuQCs, IC + DHODHi, IC + NADC, IC + 50% DMSO, and IC + HA, where IC was induced by CYP. Non-invasive procedures for intravesical treatment were described as we reported(44). In brief, a 20-gauge indwelling intravenous catheter was inserted into the bladder of rats through the urethra, and 500 µL of different agents of tanezumab (266 nM), AuQCs (1.25 µM), DHODHi (900 nM), or NADC (266 nM) was then instilled. For chronic IC, 50 mg kg^-1^ CYP was intraperitoneally administrated on days 1, 4, and 7, with intravesical therapy performed on days 8, 10, and 12. Rats were sacrificed on day 19, and bladder tissues were collected for further analysis. The AIC rat model was established using a high 150 mg kg^-1^ CYP dose(23). On day 4, rats were sacrificed, with bladders collected. Therapeutic measures were conducted at 4 h and on day 3 post-injection. The prophylactic effect of NADC was estimated by intravesical drug administration to rat models on days 1, 3, and 5, followed by 150 mg kg^-1^ intraperitoneal CYP induction of cystitis on day 6. Rats were sacrificed on day 7, with therapeutic effects assessed by conscious cystometry and Von Frey tests for bladder pain at the endpoint.

New England white rabbits at ∼2.5 kg body weight were acquired from the Guangzhou Huadong Xinhua Experimental Animal Farm. Fifteen rabbits were assigned into healthy control, IC + PBS, and IC + NADC groups. To induce IC, we injected 75 mg kg^-1^ CYP via i.p. on days 1, 4, and 7. Bladders were emptied using a 6-Fr catheter before treatment. 6 mL intravesical PBS or 266 nM NADC was instilled on days 8, 10, and 12 under anesthesia of an intramuscular Zoletil 20/xylazine hydrochloride mixture (15/5 mg kg^-1^). Rabbits were euthanized on day 19.

### mRNA sequencing

Bladder specimens were excised with autoclaved scissors, placed in a 5 mL sterile Eppendorf tube, and stored at −80 °C before sequencing. Paired-end sequencing reads of 2 x 150 bp in length were firstly trimmed and cleaned by fastp and aligned to *Rattus norvegicus* genome mRatBN7.2 (http://asia.ensembl.org/Rattus_norvegicus/Info/Index) by the HISAT2 aligner(45). PCA was performed to identify similarities among samples and replicates. RNA-seq data were investigated for differentially expressed genes (DEGs) and gene set enrichment analysis (GSEA). The R package DESeq2 v1.38.3 was used to identify the significant DEGs with false discovery rate (FDR) < 0.001 and an absolute value of log_2_ ratio ≥ 2(46). Gene annotations were downloaded from org.Rn.eg.db (Genome wide annotation for *Rattus noregicus* version 3.8.2.), and all ensemble gene IDs were mapped to symbol IDs using the R package AnnotationDbi v1.60.2 for further analysis(47). Next, a universal GSEA was conducted using the R package clusterProfiler (gseKEGG function v 4.0.5), for which the DEG set pre-ranked by their log2 fold changes was compared with the KEGG pathway dataset as a reference to identify significantly enriched pathways in DEGs(48). GO term enrichment analysis was conducted using g:Profiler portal (function g:GOSt; https://biit.cs.ut.ee/gprofiler/gost). The association of genes and KEGG pathways was depicted as a network using the R package clusterProfile (function cnetplot v4.8.2)(49).

### 16S rRNA sequencing and metrics calculation

16S rRNA amplicon sequencing was performed using probes listed in Supplementary Table 3. Paired-end sequencing reads of 2 x 251 bp in length were pre-processed using the DADA2 v1.26.0 pipeline in R, including quality filtering and trimming, paired reads merging, and chimeras removing(50). The number and percentage of reads that passed each step can be found in Supplementary Table 4. DADA2 was then used to assign the amplicon sequence variants (ASVs) to the processed reads, and taxonomy was assigned using the Silva database v138.1 to each sequence variant(51). Raw sequencing data were archived under NCBI Bioproject PRJNA1085057, and codes are available at https://github.com/WanyanW/NADC-Microbiome-Yang. Richness (observed ASVs), Chao1, and ACE indexes were calculated to represent α diversity for each sample from rarefied sequence counts. Differences in the α-diversity index between any two groups were analyzed by the Wilcoxon test. β-Diversity metrics were calculated using the Bray-Curtis distance and represented by non-metric multidimensional scaling (NMDS) using the phyloseq R package v1.42.0(52). ASVs with significant differences in abundance between groups were evaluated by the linear discriminant analysis (LDA) effect size (LEfSe) method using the R package microbiomeMarker v1.4.0, and any ASVs with a *P* value < 0.05 and an LDA score (log10) > 2 were featured as microbiome biomarkers(53). Linear regression modeling of the abundances of individual taxa against the treatment was conducted using the R package Microviz v0.11.0, and the results were plotted on the taxonomic tree generated from the taxonomic table(54). The regression coefficient estimates, test statistics, and corresponding *P* values from all these regression models can be found in Supplementary Table 5.

### Statistics

Error bars refer to standard deviation (SD), with all data presented as mean ± SD unless specified otherwise. All statistical analyses were performed with GraphPad Prism 9 (Version 9.5.1). Two-tailed Student’s *t*-tests were applied to compare two normally distributed data sets. Nonparametric tests were used for non-normally distributed data. For comparisons involving more than two means in normally distributed data, one-way or two-way ANOVA followed by appropriate post hoc tests were conducted, while nonparametric tests were applied in other cases. Unless noted, differences were considered statistically significant if the *P* value was less than 0.05.

### Study approval

All animal procedures were reviewed and authorized by the Institutional Animal Care and Use Committee (IACUC) at the Sun Yat-sen University Cancer Center (#SYSU-IACUC-2021-000986 for rats and #SYSU-IACUC-2023-000517 for rabbits). Our study exclusively examined female rats, and whether the findings are relevant to male rats is not studied. All animals in the study were randomly assigned to experimental groups. Human bladder tissues were surgically resected from para-cancerous tissues of a bladder cancer patient receiving radical cystectomy at the Sun Yat-sen University Cancer Center under a protocol approved by the ethical committee (#SL-B2023-305-01).

## Data availability

The data supporting the work’s findings are deposited and available in the *Research Data Deposit* (RDD) public medical database, with the designated identifier # (assigned upon gallery proof).

## Acknowledgments

We thank Agropur for providing the high-purity α-LA. The authors further acknowledge the National Natural Science Foundation of China (82472041, 82071978, and 52271196), the Guangdong Provincial Joint Fund for Corporate Innovation and Development (2024A1515220058), the National Key Research and Development Program of China (2021YFF1200700), the Young Talents Program of Sun Yat-sen University Cancer Center (YTP-SYSUCC-0024), and the Fundamental Research Funds for the Central Universities at Sun Yat-sen University (31610026) for supporting the study.

## Author contributions

J.Y. and Z.L. conceived and designed the study. Z.L., W.W., and D.L. conducted experiments and collected, managed, and visualized the data. W.W., Q.L., and C.W. performed bioinformatic analyses, coding, and visualization with technical help and advice from Q.Z. Further, Z.L. and Z.X. jointly performed spectral CT imaging, and Y.H. and X.Z assisted in urodynamic analyses. Z.W. helped with human tissue collection and analysis. J.Y. supervised the project. X.Z. provided resources and technical advice for the project. J.Y. and Z.L. co-wrote the manuscript, which was revised, proofread, and approved by all authors.

