## Supplemental material for "Nanocluster-antibody-drug conjugates (NADC) as an intravesical precision theranostic agent for interstitial cystitis"

*Affiliations:*
[1]State Key Laboratory of Oncology in South China, Guangdong Provincial Clinical Research Center for Cancer, Sun Yat-sen University Cancer Center, Guangzhou, 510060 China
[2]Department of Ecosystem Science and Management, Pennsylvania State University, University Park, PA, 16802 USA
[3]Department of Intensive Care Unit, Sun Yat-sen University Cancer Center, Guangzhou 510060, China
[4]Department of Urology, The Second Affiliated Hospital of Guangzhou University of Chinese Medicine, Guangzhou, 510060 China
[5]Department of Radiology, The First People's Hospital of Foshan, Foshan, 528041 China
[6]Department of Urology, The Third Affiliated Hospital of Sun Yat-sen University, Guangzhou, 510080 China
[7]Tianjin Key Laboratory of Brain Science and Neural Engineering, Academy of Medical Engineering and Translational Medicine, Tianjin University, Tianjin, 300072 China

[#]These authors contributed equally to the first-authorship

**Supplementary information**

Supplementary Methods……………………………..…………………………….………………..4

Supplementary references………………..…..……………………………..…………………11

Supplementary Table 1. Marker genes of the 20 clusters in the bladder dataset…........12

Supplementary Table 2. List of primers for RT-qPCR analysis…………….………..……..…13

Supplementary Table 3. Bacterial 16S rRNA-targeting PCR primers for sequencing…...13

Supplementary Table 4. Analysis of 16S rRNA sequencing data quality…….…………..…14

Supplementary Table 5. Linear regression modeling of the abundance of individual taxa………………………………………………………………………………………16

Supplementary Fig. S1. KEGG pathway analysis of epithelial cell-1 and epithelial cell-2.........................................................................................................................17

Supplementary Fig. S2. Signal quantification of bladder NGF expression from confocal fluorescence microscopic images……………………………………………...……………...….18

Supplementary Fig. S3. Zeta potential analysis.……………………………….....……….19

Supplementary Fig. S4. Core size distribution of AuQCs and NADC……………..……….20

Supplementary Fig. S5. Native PAGE analysis of NADC……….……………………….21

Supplementary Fig. S6. The MALDI-TOF mass spectrum analysis…………………..…..22

Supplementary Fig. S7. Characterization of Anti-NGF-CF647 by covalent labeling…….23

Supplementary Fig. S8. Fluorescence phantom imaging of anti-NGF-CF647…………...24

Supplementary Fig. S9. Time-course in vivo NIRF imaging of rats with CYP-induced cystitis after intravesical administration of NADC or anti-NGF……………………………..25

Supplementary Fig. S10. Time-course in vivo NIRF imaging of healthy rats after intravesical administration of NADC or anti-NGF……………………………………..……...26

Supplementary Fig. S11. Immunofluorescence images of bladder tissues of IC rat models with varying CYP induction doses…………………………………………………….…….27

Supplementary Fig. S12. 3D Z-stack volumetric CLSM images of IC rat bladder tissue sections post-intravesical administration of AuQCs or NADC……………………………...28

Supplementary Fig. S13. Fluorescence images of NADC uptake by SV-HUC-1 cells…..29

Supplementary Fig. S14. Phantoms imaging by the dual-energy CT……………………..30

Supplementary Fig. S15. TEER measurement of SV-HUC-1 urothelium……………...….31

Supplementary Fig. S16. Evaluation of mucosal drug transport of NADC across the SV-HUC-1 urothelium mimicry by Ussing chamber tests.…………………………....……….32

Supplementary Fig. S17. Evaluation of mucosal drug transport of NADC across healthy rat bladder tissues by Ussing chamber tests……………..………………………………33

Supplementary Fig. S18. Evaluation of mucosal drug transport of NADC across rat bladder tissues with interstitial cystitis by Ussing chamber tests…………………………..34

Supplementary Fig. S19. Evaluation of mucosal drug transport of NADC across human bladder tissues by Ussing chamber tests……………………………………………..………...35

Supplementary Fig. S20. The penetration depth of NADC in the bladder.….…………..……36

Supplementary Fig. S21. The mRNA expression profiles of pro- and anti-inflammatory cytokines by RT-qPCR in NADC-treated SV-HUC-1 cells with LPS/ATP-induced inflammation……………………………………………………………………………………..37

Supplementary Fig. S22. ROS scavenging and enzyme-mimicking activities of AuQCs in NADC ………………………………………………………………………………….………38

Supplementary Fig. S23. In vitro DHODH enzyme inhibition assay.…………………….…...39

Supplementary Fig. S24. Comparison of NADC with anti-NGF-AuQCs and anti-NGF-DHODHi in treating chronic IC in rats.………….……………………………………………40

Supplementary Fig. S25. Changes in the mechanical withdrawal threshold over one-month duration.……………………………………………………………………………41

Supplementary Fig. S26. Histopathology of bladder tissue sections from AIC rats with intravesical NADC treatment.…………………………………...…..……………………41

Supplementary Fig. S27. Additional data of NADC as a prophylactic agent for AIC in rats.…………………………………...…..……………………………………………………42

Supplementary Fig. S28. Heatmaps of significantly altered key effector genes in the NF-κB, IL-17, and JAK-STAT pathways in rat bladder tissues between the control and IC groups…………………………………………………………………………………..…43

Supplementary Fig. S29. Protein-protein interaction (PPI) network of differentially expressed proteins between the control and IC groups……………..……………..……44

Supplementary Fig. S30. PPI network analysis of differentially expressed proteins in IC rat bladder tissues with or without NADC treatment.…………………………….....……...45

Supplementary Fig. S31. Western blots of molecules involved in inflammation pathways in bladder tissues of IC rats……..…………………………………………...…………...46

Supplementary Fig. S32. Western blots of key molecules in NF-κB and JAK-STAT signaling pathways……………..…………………………………………………………47

Supplementary Fig. S33. Clinical chemistry analysis…………………….…..……………..48

Supplementary Fig. S34. Complete blood count with automated differentials………...…49

### Supplementary Methods

#### Chemicals

High-purity α-LA with a low calcium content (<0.055%) was acquired from Agropur. Gold chloride trihydrate ($HAuCl_4 \cdot 3H_2O$), cyclophosphamide (CYP), and lipopolysaccharides (LPS) were purchased from Sigma-Aldrich. Vidofludimus, sulfosuccinimidyl-4-(*N*-maleimidomethyl)cyclohexane-1-carboxylate (sulfo-SMCC), adenosine 5′-triphosphate (ATP), *N*-(3-dimethylaminopropyl)-*N*′-ethylcarbodiimide hydrochloride (EDC), and *N*-hydroxysuccinimide (NHS) were purchased from MedChemExpress. The anti-NGF therapeutic monoclonal tanezumab was purchased from Absin Bioscience. 2-Thiolanimine hydrochloride (Traut's reagent) was obtained from Shanghai Aladdin.

#### Characterization

The morphology and size of NADC were characterized using a Philips CM-200 TEM at 200 kV. UV-vis absorption spectra were recorded with a UV-2600 spectrophotometer (Shimadzu). Fluorescence measurements were recorded with a Tecan Infinite M200 plate reader. Dynamic light scattering (DLS), zeta potentials, and PDI were measured by Zetasizer Nano ZSE (Malvern Panalytical). CD spectra were monitored with an Aviv 62DS circular dichroism spectrometer (Aviv Biomedical), and secondary structure compositions were determined using the beta structure selection (BeStSel) approach.

#### Native PAGE

The chemical conjugation of AuQCs and vidofludimus to the tanezumab antibody was preliminarily analyzed using native polyacrylamide gel electrophoresis (PAGE) following standard protocols under non-reducing conditions. The bicinchoninic acid (BCA)-calibrated samples were electrophoresed on 10% polyacrylamide gels in reference to an unstained protein ladder and then visualized by Coomassie Brilliant Blue staining. Gel images were taken using the Bio-Rad ChemiDoc MP imaging system.

#### MALDI-TOF-MS

Matrix-assisted laser desorption/ionization time-of-flight mass spectrometry (MALDI-TOF-MS) was conducted with a Bruker UltrafleXtreme III MALDI-TOF/TOF high-performance mass spectrometer operating in the positive ion mode. Ionization was achieved with a smartbeam-I UV laser (355 nm). Samples for measurement were resuspended in sterile water and mixed with a matrix solution consisting of 5 mg $mL^{-1}$ 2,5-dihydroxybenzoic acid (DHB) dissolved in 30% acetonitrile. The matrix-to-sample ratio was maintained at 1:1, and a total volume of 2 μL mixture was applied to an AnchorChip probe (Bruker Daltonics), followed by air-drying. The mass spectra primarily exhibited $[M+H]^+$, $[M+Na]^+$, and $[M+K]^+$ ion signals arising from the protonation and formation of sodium and potassium adducts.

#### In vitro NADC uptake in SV-HUC-1 cells with induced inflammation

SV-HUC-1 cells were plated in tissue culture dishes (Jet Biofil, China) at a density of $3 \times 10^4$ cells $mL^{-1}$ in 1 mL complete growth medium. Seeded cells were incubated overnight and then stimulated with 10 μg $L^{-1}$ LPS and 2.5 mmol $L^{-1}$ ATP for 12 h. After stimulation,

50 μL 50 μM AuQCs or 10 μM NADC was added and further incubated for 2 h. After that, treated cells were washed thoroughly three times with 1× PBS. Finally, the cells were observed and imaged using a confocal laser scanning microscope (Carl Zeiss) equipped with filters (Ex(λ) 488 nm, Em(λ) 700 nm). Image analysis and processing were conducted with Carl Zeiss LSM software (version 2.3).

### TEER measurement of uroepithelium

The human normal urothelial cell line SV-HUC-1 used in this study was from the Cell Resource Center of the Chinese Academy of Sciences (Shanghai, China). The integrity and permeability of cellular monolayers were tested with immortalized human uroepithelial SV-HUC-1 cells by seeding into the upper chambers of 12-well plates at a density of 5 × $10^4$ cells per well. Over time, TEER was periodically measured with the RE1600 Epithelial Volt/Ohm Meter (Beijing Jingshi Technology). After continuous growth of cells to reach stabilized TEER, the upper chamber membranes with epithelial cells were used for Ussing chamber assays(1). TEER is calculated as follows:

$$TEER = (R1 - R0)/0.33$$

Where 0.33 is the surface area of the upper membrane, R0 is the TEER value of a cell-free insert, and R1 is the TEER value of an insert with cells forming stabilized tight junctions.

### Ussing chamber assays

To evaluate the mucosal drug transport of NADC across SV-HUC-1 mimicry, rat, and human bladder urothelial tissues, we carried out the Ussing chamber assays in a setup consisting of two fluid-filled, heated, and aired chambers divided into the apical and basolateral sides. Rat bladders were excised from sacrificed rats and extensively washed with cold PBS. Stripped human and rat tissues and the membrane of tight-junction SV-HUC-1 cells from above with an effective exposed area of 0.1 $cm^2$ were settled in modified Ussing chambers(2). Krebs-Henseleit Buffer Modified (KHB) was filled in the feeding and reception tanks. The system was continuously oxygenated with $O_2/CO_2$ (95/5%) and maintained at 37 °C. Next, 100 nM (Au content equivalence) AuQCs or NADC was added to the reception tank. For fluorescence measurement at each time point using a Tecan M200 plate reader, 100 μL solution was taken every 30 min from the feeding and reception tanks and supplemented with identical volumes of fresh solution on both sides.

### Cellular mRNA expressions by RT-qPCR

SV-HUC-1 cells were cultured at 37 °C in Ham's F-12K medium (Gibco) containing 10% fetal bovine serum (FBS) in a humidified atmosphere with 5% $CO_2$. Typically, 2×$10^6$ SV-HUC-1 cells were seeded in 6-well plates into six groups: control, LPS/ATP, LPS/ATP + anti-NGF antibody (10 μg $mL^{-1}$ tanezumab), LPS/ATP + AuQCs (180 nM), LPS/ATP + DHODHi (300 nM), and LPS/ATP + NADC (10 μg $mL^{-1}$ antibody basis). After 2 h treatment, cells were further stimulated with 10 μg $L^{-1}$ LPS and 2.5 mmol $L^{-1}$ ATP for another 12 h, with total RNA extracted using an RNAprep Pure Cell kit (TianGen Biotechnology). RNA concentrations were calibrated by a NanoDrop 2000 spectrophotometer (Thermo Fisher Scientific). Further, cDNA was synthesized from 1 μg total RNA with a FastKing RT Kit (TianGen Biotechnology) and subject to RT-qPCR. PCR amplification was performed with a RealStar Green Fast Mixture (GenStar) containing hot-start Taq DNA polymerase and SYBR Green I to monitor DNA amplification. Sense and antisense strands of primers were synthesized at Shanghai Sangon Biotechnology using sequences as in Supplementary Table 2. Melting curves were constantly monitored to exclude nonspecific products, and expression was calculated from the average of triplicates using $2^{-\Delta\Delta Ct}$ with reference to the glyceraldehyde-3-phosphate dehydrogenase (GAPDH) or β-actin controls.

### ROS scavenging of NADC

To evaluate the total antioxidant capacity (T-AOC) of NADC against ROS, we used the ABTS chromogenic assay using a T-AOC kit (Beyotime) and strictly followed the manufacturer's instructions. 10 μL NADC at various concentrations, 20 μL peroxidase, and 180 μL freshly prepared ABTS working solution were sequentially added to each test tube and incubated for 5 min, after which the absorbance was measured at 734 nm using UV-vis spectroscopy. The absorbance at 734 nm was also continuously monitored for 5 min to study dynamics. Samples of AuQCs, tanezumab, and vidofludimus were included as controls at molar concentrations equivalent to the corresponding ratio in NADC, with Trolox as a positive control.

### Measurement of enzyme-mimetic activities of NADC

The catalase (CAT)-like activity of NADC was assessed using the Catalase Assay Kit (Beyotime) following the manufacturer's instructions. The assay involved 520 nm absorbance measurement for both the purified catalase (1 μg) as a positive control and samples, including PBS, tanezumab (1 μM), AuQCs (5 μM), vidofludimus (3 μM), and NADC (1 μM), in the presence of 20 μM $H_2O_2$. Results were expressed in units per milligram (U $mg^{-1}$). One unit of activity is defined as the amount of CAT that catalyzes the decomposition of 1 μM $H_2O_2$ per min at 25 °C. The relative CAT activity was correlated with the catalase standard. Similarly, the superoxide dismutase (SOD)-like activity of NADC was measured using the total SOD assay kit with WST-8 (S0101, Beyotime). The WST-8/enzyme working solution and reaction initiation solution were prepared according to instructions. In a 96-well plate, 160 μL WST-8/enzyme working solution and 20 μL reaction initiation solution were added, followed by the addition of 20 μL of various samples, including SOD (1 μg), PBS, tanezumab (1 μM), AuQCs (5 μM), vidofludimus (3 μM), and NADC (1 μM). In addition, 20 μL SOD assay buffer was added instead of samples as the blank control. The plate was incubated at 37 °C for 30 min, with absorbance measured at 450 nm. The glutathione peroxidase (GPx)-like activity of NADC was also evaluated using the GPx assay kit (Beyotime). The GPx assay working solution was prepared by combining 15 μL GPx assay buffer, 2 μL 62.5 mM NADPH, 2 μL 75 mM reduced glutathione (GSH), 1 μL glutathione reductase (GR), and 10 μL 30 mM cumene hydroperoxide (Cum-OOH). The GPx assay working solution was then mixed with 20 μL PBS, tanezumab (1 μM), AuQCs (5 μM), vidofludimus (3 μM), or NADC (1 μM). The resulting mixture was incubated, and absorbance at 340 nm was monitored to assess the GPx-like activity.

### ESR spectroscopy

ESR spectroscopy was used to further study NADC's scavenging activities against specific radicals. Hydroxyl radicals ($OH^•$) were generated through the Fenton reaction by mixing 5 mM $FeCl_2$ and 25 mM $H_2O_2$ with 10 μL 50 mM 5,5-dimethyl-1-pyrroline-*N*-oxide (DMPO) spin trap. ESR spectra were recorded in the presence of 10 μL samples of PBS, tanezumab (1 μM), AuQCs (5 μM), vidofludimus (3 μM), or NADC (1 μM) with a JES-FA200 ESR spectrometer (JEOL) at 1 mW microwave power and 1 G modulation amplitude. Likewise, $O_2^{•-}$ radicals were generated using the ammonium persulfate-*N*,*N*,*N'*,*N'*-tetramethylethylenediamine (AP-TEMED) method. Solution A was prepared by dissolving 9.5 g of Tris in 12 mL 0.1 M HCl, followed by adding 0.1 mL TEMED and diluting the final volume to 25 mL. In parallel, Solution B was prepared by dissolving 0.28 g AP in 100 mL water. The reaction mixture of 20 μL Solution A and 100 μL Solution B was allowed to proceed for 1 min. Subsequently, a 10 μL sample of PBS, tanezumab (1 μM), AuQCs (3 μM), vidofludimus (5 μM), or NADC (1 μM) was added, and then 10 μL 50 mM 5-diethoxyphosphoryl-5-methyl-1-pyrroline-*N*-oxide (DEPMPO) was included as the spin trap.

### DHODH enzyme inhibition assay

The biochemical detection of the DHODH enzymatic activity relies on the oxidation of *L*-dihydroorotic acid (*L*-DHO) and the reduction of 2,6-dichloroindophenol (DCIP) and decylubiquinone (DUQ), leading to a decrease in absorbance. The inhibitory effects of vidofludimus and NADC on DHODH were evaluated by coupled zymographic spectrophotometry. 25 ng DHODH enzyme (Abmart) was dissolved in an assay buffer consisting of 50 mM pH 8 Tris HCl, 150 mM KCl, and 0.8% Triton X-100 and added to a 96-well plate containing gradient concentrations of DHODHi or NADC. Next, a substrate mixture of 20 mM *L*-DHO, 2 mM DUQ, and 2 mM DCIP was added to initiate the redox reaction. After 30 min, the decrease in absorbance at 600 nm was measured. $IC_{50}$ values for the DHODH inhibitors were determined using GraphPad Prism.

### In vivo near-infrared fluorescence imaging

Female Sprague-Dawley (SD) rats weighed 220 ± 20 g were regularly kept in a constant environment at 24 °C with a 12-h diurnal cycle. 500 μL PBS, AuQCs (15 μM), or NADC (5 μM) was instilled by intravesical delivery using a 20-gauge catheter needle. Non-invasive in vivo NIRF imaging was conducted using an IVIS Spectrum system (PerkinElmer). Rats were fasted for 4 h and shaved at the abdominal region before imaging. After intravesical administration of AuQCs or NADC, NIRF images were taken at various time points post-injection under anesthesia and processed by multispectral unmixing to distinguish AuQCs-specific signals from autofluorescence using the Living Image software (PerkinElmer)(3, 4). Similarly, NIRF images were taken at fixed time points after intravesical injection of anti-NGF-CF647 or NADC. The average fluorescence signal intensity of the region of interest (ROI) in the bladder area was calculated following the identification from bright-field images.

### The von Frey test

As the hallmark syndrome for IC, evoked bladder pain was assessed by the mechanical withdrawal responses with von Frey filaments(5). Rats were placed on a broad-gauge wire mesh surface and acclimatized to the chamber environment for 30 min before testing. Different calibrated von Frey filaments with gradually increasing forces from a 1-g filament were administrated to the lower abdominal areas and poked onto the skin until minor bending was reached under the 50% withdrawal threshold technique. Withdrawal responses of rats, such as sudden retracting, lower abdomen licking, and jumping, were observed after reaching their threshold. Likewise, filaments from 1-26 g were applied sequentially to the plantar surface of the rabbit's hind paw in ascending order. If a twitch response was observed following 1 g filament, filaments ranging from 0.6 to 0.008 g were applied in descending order until no additional response was detected. Each paw was minimally tested five times to ensure specific filament responses, with the force consistently triggering withdrawal responses recorded. In the absence of a withdrawal response to the 26 g von Frey filament, the paw was classified as non-responsive and assigned a default threshold value of 60 g. The mean withdrawal force from the bilateral hind paws was computed for subsequent statistical evaluation.

### Conscious cystometry for bladder function testing

The bladder catheter was linked through a three-way stopcock to a BL-420N (information) biological signal acquisition and processing system (Chengdu Thai unita software) with a microinjection pump at the same height level as the bladder. Repeated bladder contractions were induced with 6 mL $h^{-1}$ saline infusion at room temperature(6). After an initial stabilization period of 30 min, the fluidic pressure and volume in the bladder were measured during filling, storage, and voiding cycles, with a minimum of four cycles recorded. Data on intersystolic and maximal voiding pressures, voiding pressure threshold, and baseline resting pressure were derived from the curves. Cystometry for rabbits was conducted similarly. Rabbits were sedated by intramuscular injection of 0.7 mL $kg^{-1}$ Zoletil 20/xylazine hydrochloride (15/5 mg $kg^{-1}$). In the supine position, a 6-French double-lumen transurethral catheter was inserted into the bladder and connected to a pressure transducer. The bladder was perfused with normal saline at room temperature at a flow rate of 2 mL $min^{-1}$, with intravesical pressure recorded simultaneously.

### Open field test

Abdominal pain has been shown to affect general activity levels, gross locomotor activity, and exploration behaviors, impacting patients' quality of life(7). We employed sensorimotor OFT to study the influence of IC on rodent exploration footage. Rats with AIC receiving various treatments were placed in a clean chamber and allowed to move freely for a while, with an overhead video camera that covered the entire arena for recording. After that, the rat footage was recorded for 6 min at 30 frames $s^{-1}$ (approximately 10,800 total frames) and analyzed using the Manual Tracking Manager plugin in ImageJ 1.8.0 (National Institutes of Health). The tail base of each rat in all frames was clicked and set to track the movement. Data were displayed in the rat's coordinates (X, Y) in each frame. The following formula is used to calculate the movement distance in each frame:

$$Distance = \sqrt{(x_2 - x_1)^2 + (y_2 - y_1)^2}$$

Where ($x_1$, $y_1$) dictates the position in the first frame and ($x_2$, $y_2$) represents the position in the second frame. The cumulative distance is quantified by adding the distance between each frame over a pre-defined period.

### Macroscopic analysis of bladder gross pathology

Rats with IC were euthanized at pre-defined time points with bladders excised, weighed, and examined macroscopically for hemorrhage and edema. Bladder tissues were pathologically scored according to the following criteria: 3 points as severe (severe edema of the bladder wall with significant bleeding), 2 points as moderate (moderate edema of the bladder wall with minor bleeding), 1 point as mild (minor edema with no bleeding), and 0 points as no effects(8, 9).

### Histology

For histopathological examination, freshly resected bladder tissues were fixed in 4 % methanol-free paraformaldehyde, embedded in paraffin blocks, and sectioned into 8 μm in thickness. After staining by hematoxylin and eosin (H&E), the bladder sections were evaluated for the severity of inflammation according to the following grading criteria: 0 as normal, 1 as subepithelial inflammatory infiltration (focal and multifocal), 2 as edema and subepithelial inflammatory cell infiltration (diffuse), 3 as marked necrosis and neutrophils in and on the surface of the bladder mucosal epithelium with subepithelial inflammatory cells, 4 as inflammatory cell infiltration extending into muscles, and 5 as loss of superficial epithelial cells in addition to grade 3(10, 11).

### Immunofluorescence analysis

We stained NGF with a labeled anti-NGF monoclonal antibody to track NADC in bladder mucosa and correlate with NGF expression. Typically, 130 nM CF488A in *N*-hydroxysuccinimide (NHS) ester (Biotium) was mixed with 2 μg anti-NGF antibodies in 1 mL PBS under stirring for 1 h at room temperature. Purification was achieved using a 3 kDa MWCO ultrafiltration tube with the retentate collected and stored at 4°C in the dark. Similarly, 130 nM CF647 NHS ester (Biotium) was added to 2 μg $mL^{-1}$ anti-NGF antibody and then purified to derive anti-NGF-CF647. After being anesthetized with 7% chloral hydrate, rats with CYP-induced cystitis were administered 500 μL AuQCs or NADC for 12 h by an intravesical catheter. Bladders were then isolated from rats, rinsed with PBS, and embedded in OCT for fresh cryosections. 8 μm thick sections were cut, blocked with 2% BSA for 1 h at room temperature, and stained with CF488A labeled anti-NGF antibodies (2 μg $mL^{-1}$) for 1 h at room temperature, with DAPI counterstaining (Beyotime)(12, 13). After mounting, confocal imaging was performed using an LSM880 confocal laser scanning microscope (Zeiss) for colocalization analysis.

### Western blot analyses

Harvested bladder tissues were homogenized in the lysis buffer (20 mM HEPES–KOH, pH 7.5, 10 mM KCl, 1.5 mM $MgCl_2$, 0.2 mM EDTA, 0.1% Triton-X100), with protein concentrations calibrated with a BCA kit (Pierce)(14). 20 μg proteins were loaded onto sodium dodecyl-sulfate polyacrylamide gel electrophoresis (SDS-PAGE) gels and blotted onto polyvinylidene fluoride (PVDF) membranes(15). After blocking, membranes were stained at 4 °C with the following primary antibodies at 1:1000 dilutions: tumor necrosis factor alpha (TNF-α, ab66579, Abcam), interleukin-1 beta (IL-1β, ab9722, Abcam), IL-6 (DF6087, Affinity Biosciences), NGF (ab52918, Abcam), IL-17A (ab79056, Abcam), tropomyosin receptor kinase A (TrkA, DF6822, Affinity Biosciences), phospho-TrkA (AF3072, Affinity Biosciences), nuclear factor kappa B (NF-κB, AF5006, Affinity Biosciences), phospho-NF-κB (AF2006, Affinity Biosciences), inhibitory-κB kinase alpha (IKK-α, AF6012, Affinity Biosciences), phospho-IKK-α (AF3012, Affinity Biosciences), IKK-β (AF6009, Affinity Biosciences), phospho-IKK-β (AF3010, Affinity Biosciences), Janus kinase-1 (JAK-1, AF5012, Affinity Biosciences), phospho-JAK-1 (AF2012, Affinity Biosciences), signal transducer and activator of transcription 1 (STAT1, AF6300, Affinity Biosciences), phospho-STAT1 (AF3300, Affinity Biosciences), STAT2 (AF6342, Affinity Biosciences), phospho-STAT2 (AF3342, Affinity Biosciences), and β-actin (T0022, Affinity Biosciences). After further staining with goat anti-rabbit or goat anti-mouse matching secondary antibodies for 1 h, the blots were developed using Clarity ECL Western Blotting Substrate (Bio-Rad) for chemiluminescence. Band signals of proteins were analyzed with ImageJ.

**Supplementary Table S1: Marker genes of the 20 clusters in the bladder dataset.**

| Cluster | Cell type | Marker gene |
|---|---|---|
| 0 | Epithelial cell-1 | KRT19, KRT18, KRT17 |
| 1 | Fibroblast-1 | VIM, TAGLN, COL1A2 |
| 2 | T cell-1 | CD3D, CD3E, CD2 |
| 3 | Immune cell | IL6, IL1B, CXCL8 |
| 4 | Macrophage-1 | CD68, CD163, CD14 |
| 5 | Plasma cell | IGHG3, JCHAIN, HLA-DPA1 |
| 6 | Fibroblast-2 | COL1A2, COL6A1, COL6A2 |
| 7 | B cell-1 | MS4A1, CD19, CD79A |
| 8 | T cell-2 | CD3E, CD3D, CD2 |
| 9 | Epithelial cell-2 | KRT7, KRT13, KRT19 |
| 10 | Smooth muscle cell | ACTA2, MYL9, MYH11 |
| 11 | Endothelial cell | PECAM1, VWF, EMCN |
| 12 | B cell-2 | IGHG1, CD38, CD79A |
| 13 | Neutrophil | S100A8, S100A9, FCGR3B |
| 14 | T cell-3 | CD3D, CD3E, CXCR4 |
| 15 | Mast cell | TPSB2, TPSAB1, CPA3 |
| 16 | Macrophage-2 | CD68, C1QB, C1QA |
| 17 | Fibroblast-3 | COL6A1, MMP2, FN1 |
| 18 | Schwann cell | S100B, PLP1, MPZ |
| 19 | B cell-3 | MS4A1, CD79A, POU2AF1 |

**Supplementary Table S2: List of primers for RT-qPCR analysis.**

| Genes | Sequences (5'→3') |
|---|---|
| *hIL-1β* forward | CCACAGACCTTCCAGGAGAATG |
| *hIL-1β* reverse | GTGCAGTTCAGTGATCGTACAGG |
| *hTNF-α* forward | CTCTTCTGCCTGCTGCACTTTG |
| *hTNF-α* reverse | ATGGGCTACAGGCTTGTCACTC |
| *hIL-6* forward | AGACAGCCACTCACCTCTTCAG |
| *hIL-6* reverse | TTCTGCCAGTGCCTCTTTGCTG |
| *hIL-10* forward | TCTCCGAGATGCCTTCAGCAGA |
| *hIL-10* reverse | TCAGACAAGGCTTGGCAACCCA |
| *hIL-4* forward | CCGTAACAGACATCTTTGCTGC |
| *hIL-4* reverse | GAGTGTCCTTCTCATGGTGGCT |
| *hIL-13* forward | ACGGTCATTGCTCTCACTTGCC |
| *hIL-13* reverse | CTGTCAGGTTGATGCTCCATACC |

**Supplementary Table S3: Bacterial 16S rRNA-targeting PCR primers for sequencing.**

| Name | Primers |
|---|---|
| 338F | ACTCCTACGGGAGGCAGCAG |
| 806R | GGACTACHVGGGTWTCTAAT |

H: A/T/C. V: G/A/C. W: A/T

**Supplementary Table S4: Analysis of 16S rRNA sequencing data quality.**

| | Input | Filtered | DenoisedF | DenoisedR | Merged | Nonchim |
|---|---|---|---|---|---|---|
| Control 1 | 311159 | 254502 | 253068 | 251845 | 247936 | 196528 |
| Control 2 | 310374 | 256173 | 254598 | 251807 | 246075 | 194994 |
| Control 3 | 317420 | 254733 | 253489 | 251590 | 243267 | 190992 |
| Control 4 | 318242 | 258871 | 257722 | 254732 | 248269 | 183220 |
| Control 5 | 299463 | 249351 | 248512 | 246539 | 243122 | 193688 |
| IC+PBS 1 | 277375 | 184357 | 182805 | 182583 | 177404 | 136917 |
| IC+PBS 2 | 330106 | 266076 | 264790 | 264507 | 262839 | 110454 |
| IC+PBS 3 | 279762 | 177375 | 176152 | 176045 | 174290 | 77847 |
| IC+PBS 4 | 344315 | 277354 | 273105 | 275052 | 264465 | 192774 |
| IC+PBS 5 | 284356 | 236082 | 235625 | 235049 | 233625 | 181786 |
| IC+Anti-NGF 1 | 278031 | 199143 | 196757 | 197554 | 193869 | 156931 |
| IC+Anti-NGF 2 | 280831 | 240291 | 239580 | 238400 | 236450 | 192533 |
| IC+Anti-NGF 3 | 304277 | 247229 | 245595 | 244824 | 236505 | 189971 |
| IC+Anti-NGF 4 | 292487 | 246020 | 245456 | 244271 | 242821 | 197616 |
| IC+Anti-NGF 5 | 290498 | 239958 | 239167 | 238179 | 235427 | 188404 |
| IC+AuQCs 1 | 281329 | 181477 | 179808 | 179196 | 175693 | 129751 |
| IC+AuQCs 2 | 274884 | 228036 | 226485 | 225258 | 222119 | 175008 |
| IC+AuQCs 3 | 256316 | 177115 | 175371 | 174920 | 167986 | 130235 |
| IC+AuQCs 4 | 286475 | 236674 | 235679 | 233375 | 229240 | 176564 |
| IC+AuQCs 5 | 248890 | 162285 | 161486 | 161520 | 160908 | 129177 |
| IC+DHODHi 1 | 309606 | 252431 | 251748 | 251104 | 250022 | 159918 |
| IC+DHODHi 2 | 312577 | 250586 | 246594 | 248510 | 243037 | 192678 |
| IC+DHODHi 3 | 299893 | 248885 | 247374 | 246166 | 240101 | 194002 |
| IC+DHODHi 4 | 301351 | 207343 | 206231 | 206029 | 203801 | 150542 |
| IC+DHODHi 5 | 361602 | 291535 | 290377 | 288372 | 278253 | 219054 |
| IC+NADC 1 | 279637 | 217962 | 216507 | 215186 | 209636 | 166298 |
| IC+NADC 2 | 272994 | 227858 | 226211 | 225478 | 221338 | 173387 |
| IC+NADC 3 | 308998 | 252296 | 249342 | 249542 | 241610 | 191301 |
| IC+NADC 4 | 323864 | 269014 | 268103 | 267009 | 263593 | 207376 |
| IC+NADC 5 | 322157 | 265567 | 264563 | 262985 | 257840 | 207310 |

Input: the number of reads present in the initial FASTQ file. Filtered: the count of reads subsequent to preliminary quality filtration. Denoised (F/R): the number of reads following error rate inference and denoised. Merged: the total of paired reads that have been merged. Nonchim: the count of reads merged post the elimination of chimeric sequences.

### Supplementary Table S5: Linear regression modeling of the abundance of individual taxa.

| | Rank | Formula | Estimate | Std.error | Statistic | *P*.value | *P*.adj.BH.rank |
|---|---|---|---|---|---|---|---|
| Control vs. IC+PBS | Class | Cyanobacteriia | -5.0635157 | 2.0896802 | -2.4231056 | 0.041648 | 0.43611915 |
| | | Chlamydiae | -7.0282977 | 3.00725372 | -2.337115 | 0.04763 | 0.43611915 |
| | Order | Rickettsiales | -6.1167537 | 2.59265875 | -2.3592591 | 0.046011 | 0.44956694 |
| | | Bifidobacteriales | -3.841077 | 1.1376995 | -3.3761789 | 0.009696 | 0.44956694 |
| | | Chlamydiales | -7.0282977 | 3.00725372 | -2.337115 | 0.04763 | 0.44956694 |
| | Family | Rickettsiales Order | -5.5099006 | 2.06380257 | -2.6697809 | 0.028372 | 0.49736178 |
| | | Burkholderiaceae | -10.027092 | 2.87393573 | -3.4889757 | 0.008211 | 0.49736178 |
| | | JG30-KF-CM45 | -2.3371314 | 0.97917869 | -2.3868282 | 0.044073 | 0.49736178 |
| | | Cellulomonadaceae | -9.7739299 | 3.57114182 | -2.7369201 | 0.025571 | 0.49736178 |
| | | Bifidobacteriaceae | -3.841077 | 1.1376995 | -3.3761789 | 0.009696 | 0.49736178 |
| | | Prolixibacteraceae | -2.8302881 | 1.20061535 | -2.3573646 | 0.046148 | 0.49736178 |
| | Genus | Rickettsiales Order | -5.5099006 | 2.06380257 | -2.6697809 | 0.028372 | 0.4682981 |
| | | JG30-KF-CM45 Family | -2.3371314 | 0.97917869 | -2.3868282 | 0.044073 | 0.4682981 |
| | | Aquabacterium | -5.4307213 | 1.31799118 | -4.1204534 | 0.003342 | 0.4682981 |
| | | Cellulomonadaceae Family | -5.6321437 | 1.54572184 | -3.6436981 | 0.006554 | 0.4682981 |
| | | Bifidobacterium | -3.841077 | 1.1376995 | -3.3761789 | 0.009696 | 0.4682981 |
| | | Prolixibacteraceae Family | -2.8302881 | 1.20061535 | -2.3573646 | 0.046148 | 0.4682981 |
| | | Patulibacter | -4.7408693 | 1.74067881 | -2.723575 | 0.026104 | 0.4682981 |
| | | Hyphomicrobium | -4.2469681 | 1.8052412 | -2.3525766 | 0.046494 | 0.4682981 |
| | | Ilumatobacter | -4.5200523 | 1.87342778 | -2.4127177 | 0.042328 | 0.4682981 |
| | | Proteiniclasticum | -4.0838601 | 1.70578756 | -2.39412 | 0.043575 | 0.4682981 |
| | | Roseomonas | -4.5105921 | 1.84451094 | -2.4454136 | 0.040223 | 0.4682981 |
| | | Thermomonas | -4.8762702 | 1.92377422 | -2.5347414 | 0.034996 | 0.4682981 |
| | | Paraclostridium | -3.2760683 | 1.34904968 | -2.4284267 | 0.041303 | 0.4682981 |
| | | Ileibacterium | -3.4672245 | 1.46207603 | -2.3714393 | 0.045145 | 0.4682981 |
| | | Xanthobacteraceae Family | -3.7713311 | 1.58868155 | -2.3738748 | 0.044974 | 0.4682981 |
| Control vs. IC+NADC | Phylum | ASV161 | 4.91882112 | 2.00951604 | 2.44776405 | 0.040076 | 0.39315084 |
| | Class | ASV161 | 4.91882112 | 2.00951604 | 2.44776405 | 0.040076 | 0.40376357 |
| | Order | ASV161 | 4.91882112 | 2.00951604 | 2.44776405 | 0.040076 | 0.44091959 |
| | | Oscillospirales`~DiseaseState | -8.6162083 | 2.87996585 | -2.9917745 | 0.017287 | 0.44091959 |
| | | Christensenellales`~DiseaseState | -3.4510905 | 1.41510423 | -2.4387536 | 0.040643 | 0.44091959 |
| | Family | Enterobacteriaceae`~DiseaseState | -7.8259109 | 3.32381825 | -2.3544942 | 0.046355 | 0.48430253 |
| | | Enterococcaceae`~DiseaseState | -4.2193915 | 1.4520846 | -2.9057477 | 0.019717 | 0.48430253 |
| | | ASV161 | 4.91882112 | 2.00951604 | 2.44776405 | 0.040076 | 0.48430253 |
| | | Cellulomonadaceae`~DiseaseState | -11.58912 | 3.28606356 | -3.5267486 | 0.007769 | 0.48430253 |
| | | Microscillaceae`~DiseaseState | -7.1319076 | 3.04117845 | -2.3451132 | 0.047039 | 0.48430253 |
| | | Christensenellaceae`~DiseaseState | -3.4510905 | 1.41510423 | -2.4387536 | 0.040643 | 0.48430253 |
| | Genus | Escherichia-Shigella`~DiseaseState | -7.2463087 | 1.55720069 | -4.6534199 | 0.001637 | 0.47656607 |
| | | Enterococcus`~DiseaseState | -4.2193915 | 1.4520846 | -2.9057477 | 0.019717 | 0.47656607 |
| | | ASV161 | 4.91882112 | 2.00951604 | 2.44776405 | 0.040076 | 0.47656607 |
| | | Luteimonas`~DiseaseState | -4.5283227 | 1.77815763 | -2.5466374 | 0.034354 | 0.47656607 |
| | | Shinella`~DiseaseState | -4.2621188 | 1.83890819 | -2.3177442 | 0.049092 | 0.47656607 |

| | | | | | | | |
|---|---|---|---|---|---|---|---|
| | | Actinotalea`~DiseaseState | -5.4316148 | 1.49949751 | -3.62229 | 0.00676 | 0.47656607 |
| | | Ohtaekwangia`~DiseaseState | -4.2335336 | 1.78737303 | -2.3685786 | 0.045347 | 0.47656607 |
| | | Ilumatobacter`~DiseaseState | -4.5200523 | 1.87342778 | -2.4127177 | 0.042328 | 0.47656607 |
| | | Proteiniclasticum`~DiseaseState | -4.0838601 | 1.70578756 | -2.39412 | 0.043575 | 0.47656607 |
| | | Flavobacteriaceae Family`~DiseaseState | -4.1740549 | 1.24360286 | -3.3564211 | 0.009985 | 0.47656607 |
| | | Lactobacillus`~DiseaseState | -3.1626929 | 1.31418624 | -2.4065789 | 0.042736 | 0.47656607 |
| | | Paraclostridium`~DiseaseState | -3.2760683 | 1.34904968 | -2.4284267 | 0.041303 | 0.47656607 |
| | | Christensenellaceae R-7 group`~DiseaseState | -3.4510905 | 1.41510423 | -2.4387536 | 0.040643 | 0.47656607 |
| | | Vibrionimonas`~DiseaseState | 2.90177939 | 1.23188704 | 2.35555639 | 0.046278 | 0.47656607 |
| IC+PBS vs. IC+NADC | Phylum | ASV161 | 4.85138199 | 1.9820038 | 2.44771579 | 0.040079 | 0.4257303 |
| | Class | ASV161 | 4.85138199 | 1.9820038 | 2.44771579 | 0.040079 | 0.4475983 |
| | | Thermoleophilia`~DiseaseState | 9.89175335 | 3.670526 | 2.69491439 | 0.027288 | 0.4475983 |
| | | KD4-96`~DiseaseState | 3.46841323 | 1.48286432 | 2.33899567 | 0.04749 | 0.4475983 |
| | Order | ASV161 | 4.85138199 | 1.9820038 | 2.44771579 | 0.040079 | 0.48465437 |
| | | Alphaproteobacteria Class`~DiseaseState | 5.7115867 | 1.57544011 | 3.62539118 | 0.00673 | 0.48465437 |
| | | KD4-96 Class`~DiseaseState | 3.46841323 | 1.48286432 | 2.33899567 | 0.04749 | 0.48465437 |
| | Family | Enterobacteriaceae`~DiseaseState | -17.369303 | 5.28625502 | -3.2857482 | 0.011092 | 0.49283749 |
| | | Enterococcaceae`~DiseaseState | -6.741478 | 2.91117893 | -2.315721 | 0.049247 | 0.49283749 |
| | | Rickettsiales Order`~DiseaseState | 4.89135124 | 2.05570668 | 2.37940134 | 0.044587 | 0.49283749 |
| | | Caulobacteraceae`~DiseaseState | 4.28486235 | 1.32974592 | 3.22231659 | 0.012197 | 0.49283749 |
| | | Corynebacteriaceae`~DiseaseState | -1.5820229 | 0.64276869 | -2.4612631 | 0.039241 | 0.49283749 |
| | | ASV161 | 4.85138199 | 1.9820038 | 2.44771579 | 0.040079 | 0.49283749 |
| | | Intrasporangiaceae`~DiseaseState | 4.36514543 | 1.7147563 | 2.54563604 | 0.034408 | 0.49283749 |
| | | Comamonadaceae`~DiseaseState | 12.7521462 | 4.80950877 | 2.65144462 | 0.02919 | 0.49283749 |
| | | Alphaproteobacteria Class`~DiseaseState | 5.7115867 | 1.57544011 | 3.62539118 | 0.00673 | 0.49283749 |
| | | JG30-KF-CM45`~DiseaseState | 1.81497891 | 0.54996825 | 3.30015218 | 0.010856 | 0.49283749 |
| | | KD4-96 Class`~DiseaseState | 3.46841323 | 1.48286432 | 2.33899567 | 0.04749 | 0.49283749 |
| | Genus | Escherichia-Shigella`~DiseaseState | -10.548622 | 2.95464618 | -3.5701813 | 0.007292 | 0.45331895 |
| | | Enterococcus`~DiseaseState | -6.741478 | 2.91117893 | -2.315721 | 0.049247 | 0.45331895 |
| | | Rickettsiales Order`~DiseaseState | 4.89135124 | 2.05570668 | 2.37940134 | 0.044587 | 0.45331895 |
| | | Corynebacterium`~DiseaseState | -1.5820229 | 0.64276869 | -2.4612631 | 0.039241 | 0.45331895 |
| | | ASV161 | 4.85138199 | 1.9820038 | 2.44771579 | 0.040079 | 0.45331895 |
| | | Terrabacter`~DiseaseState | 3.68115134 | 1.57699909 | 2.33427613 | 0.047841 | 0.45331895 |
| | | Pelomonas`~DiseaseState | 3.73896685 | 1.34258744 | 2.78489635 | 0.023745 | 0.45331895 |
| | | Alphaproteobacteria Class`~DiseaseState | 5.7115867 | 1.57544011 | 3.62539118 | 0.00673 | 0.45331895 |
| | | Faecalibaculum`~DiseaseState | -5.2810225 | 1.40331511 | -3.7632478 | 0.005519 | 0.45331895 |
| | | JG30-KF-CM45 Family`~DiseaseState | 1.81497891 | 0.54996825 | 3.30015218 | 0.010856 | 0.45331895 |
| | | Microbacterium`~DiseaseState | -4.9514005 | 1.83944363 | -2.6917924 | 0.02742 | 0.45331895 |
| | | Shinella`~DiseaseState | -4.5781895 | 1.90091318 | -2.4084159 | 0.042613 | 0.45331895 |
| | | Pedobacter`~DiseaseState | 4.62871199 | 1.28588127 | 3.59964183 | 0.006986 | 0.45331895 |
| | | KD4-96 Class`~DiseaseState | 3.46841323 | 1.48286432 | 2.33899567 | 0.04749 | 0.45331895 |

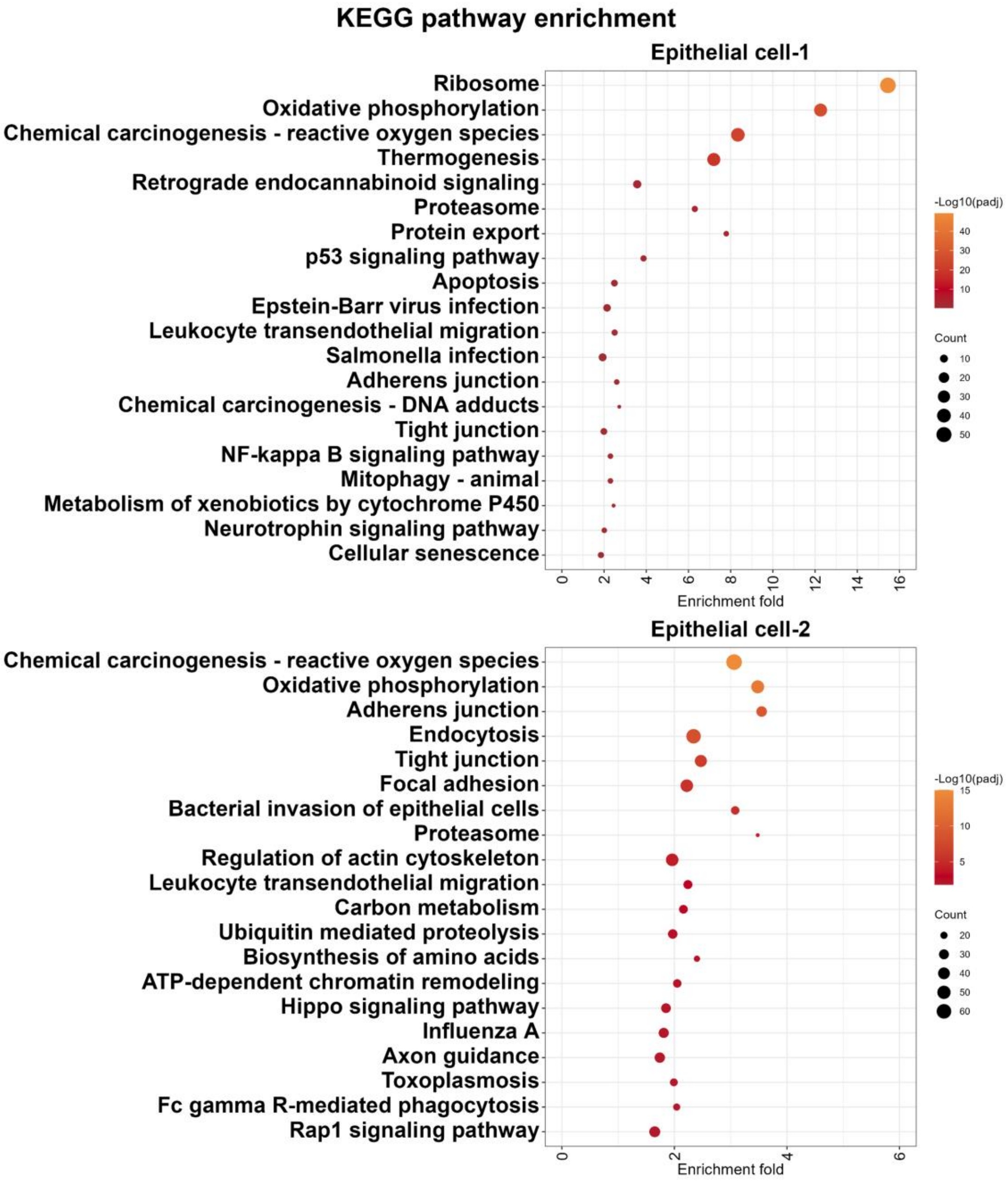


**Fig. S1. KEGG pathway analysis of epithelial cell-1 and epithelial cell-2.** The top 20 enriched pathways with the smallest p.adjust values are shown. Pathway circles are colored from red to yellow to indicate increasing significance, with circle sizes representing the number of enriched genes.

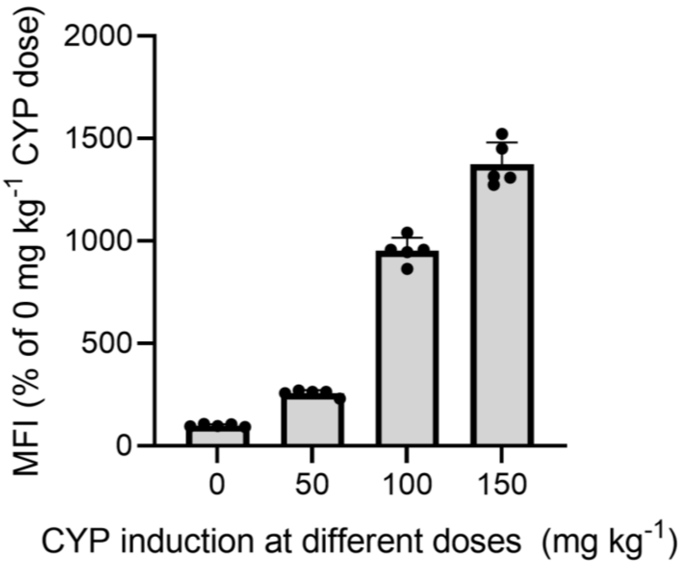


**Fig. S2. Signal quantification of bladder NGF expression from confocal fluorescence microscopic images.** Signal intensities were quantified for images taken by confocal fluorescence microscopy in Fig. 2g. As the CYP induction dose increased, the NGF expression in the bladder mucosa denoted by fluorescence was progressively elevated (n = 5).

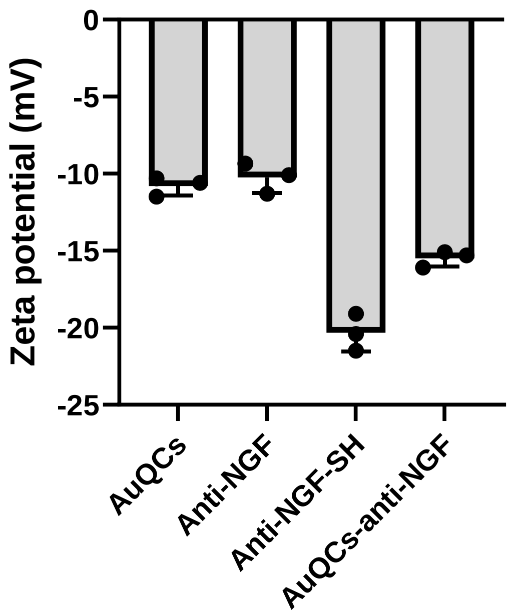


**Fig. S3. Zeta potentials of stepwise synthesized antibody conjugates.** While zwitterionic AuQCs and the unmodified anti-NGF antibody, tanezumab, exhibited a low zeta potential below ~ -10 mV, lysine side chain-based (isoelectric point=9.74) thiolation, which reduces positive charges and deprotonates, raises the zeta potential to a larger negative value.

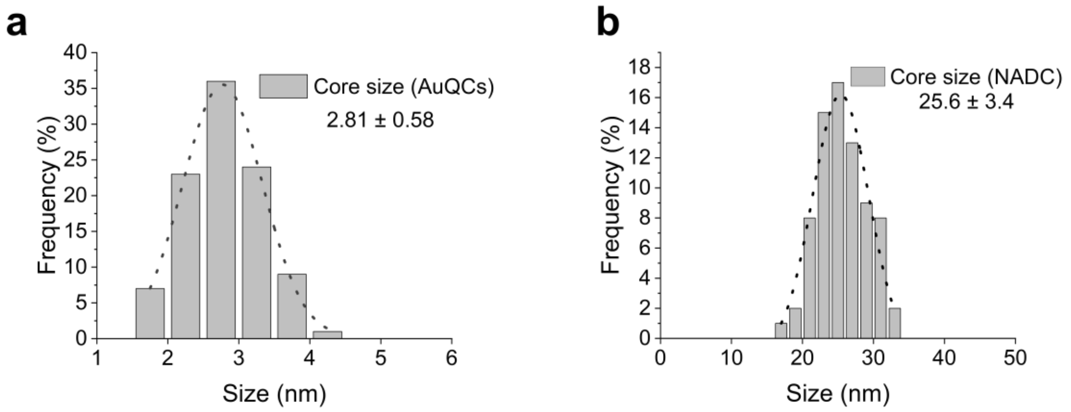


**Fig. S4. Core size distribution of AuQCs and NADC.** (a) AuQCs are ultrasmall < 3 nm. (b) The diameter of NADC reaches ~25 nm.

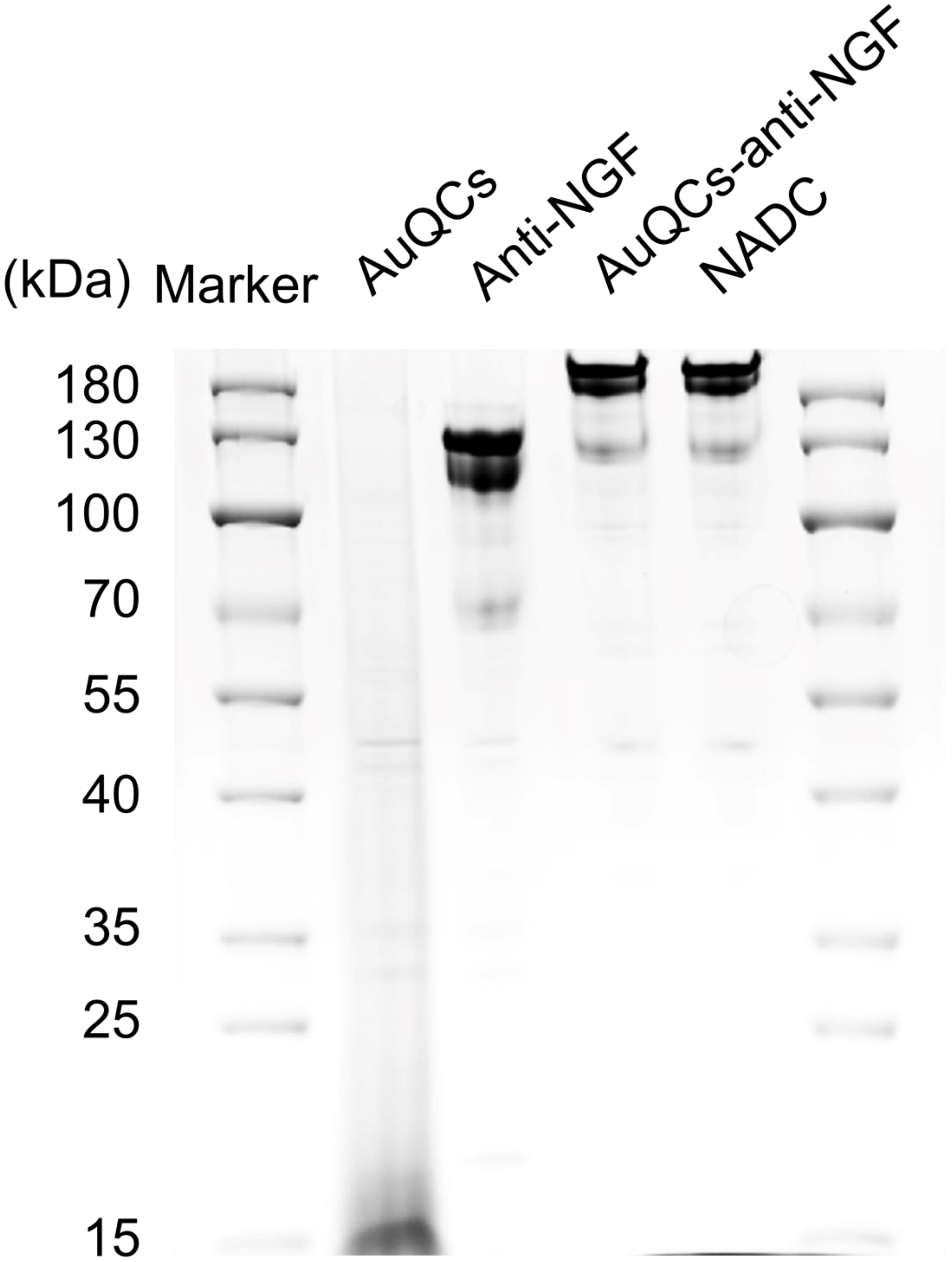


**Fig. S5. Native PAGE analysis of NADC.** A native PAGE gel was run with 10% polyacrylamide under the non-reducing condition (no β-mercaptoethanol and no heating). The gel was then stained with Coomassie Brilliant Blue and imaged with native gel molecular weight ladders.

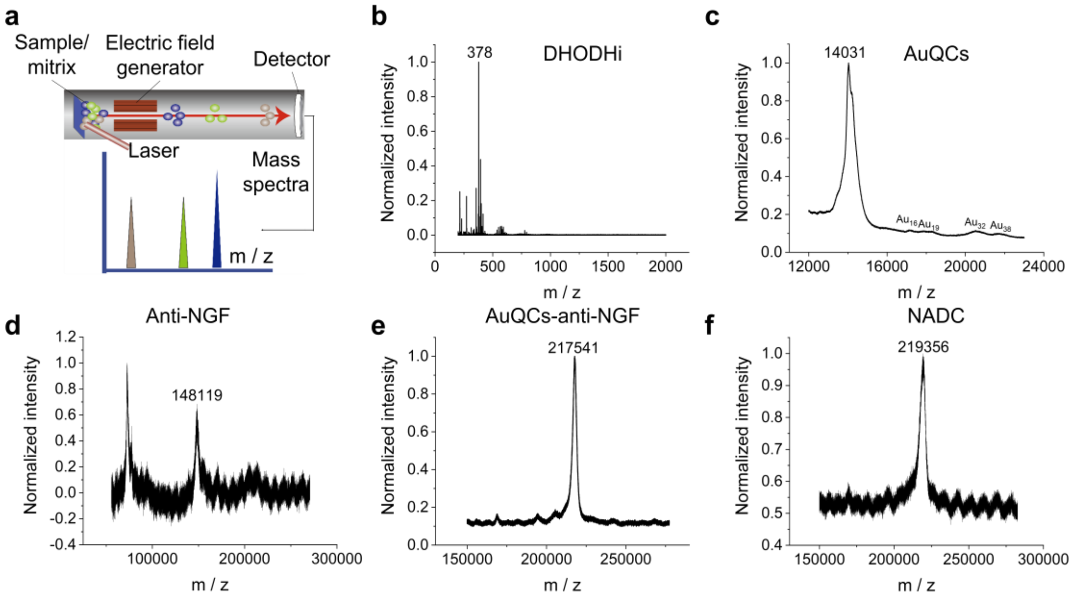


**Fig. S6. The MALDI-TOF mass spectrum analysis.** (a) Schematic representation showing the MALDI-TOF mass spectrometric measurement. MALDI-TOF-MS spectra of (b) DHODHi, (c) AuQCs, (d) anti-NGF, (e) anti-NGF-AuQCs, and (f) NADC. The number of α-LA/AuQCs ligands calculated from the difference between anti-NGF-AuQCs and tanezumab is ~4.95. The added molecular weight from the crosslinker is 219.1, resulting in calculated ~3.2 DHODHi molecules by the m/z shift in the NADC final product. The MALDI data agree well with the experimental NDAR ratio.

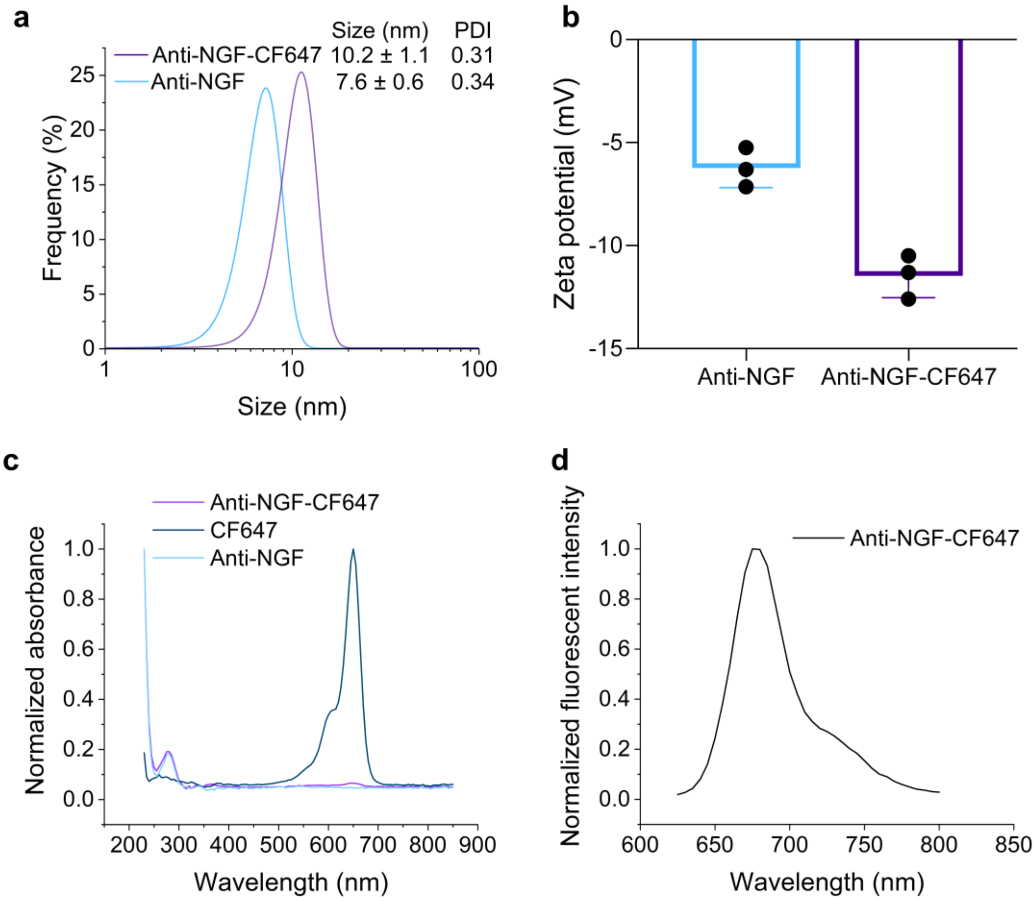


**Fig. S7. Characterization of anti-NGF-CF647 by covalent labeling.** (a) DLS analysis for the hydrodynamic size distribution and (b) zeta potential measurement of unlabeled anti-NGF and labeled anti-NGF-CF647 antibodies (n = 3). (c) Normalized UV-vis absorption spectra showing feature peaks of antibody molecules and CF647 dyes. (d) Normalized fluorescence emission spectra of anti-NGF-CF647 excited at 550 nm.

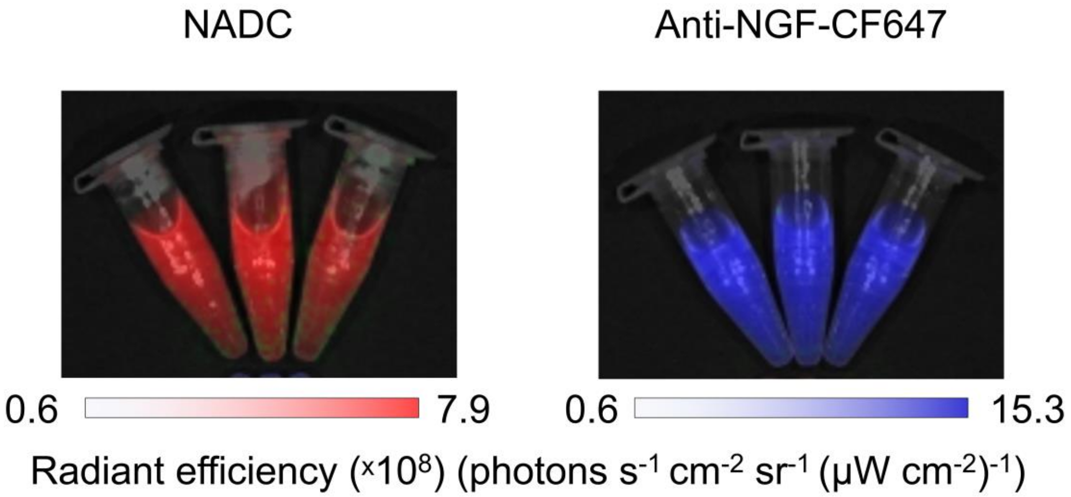


**Fig. S8. Fluorescence phantom imaging of NADC and anti-NGF-CF647.**

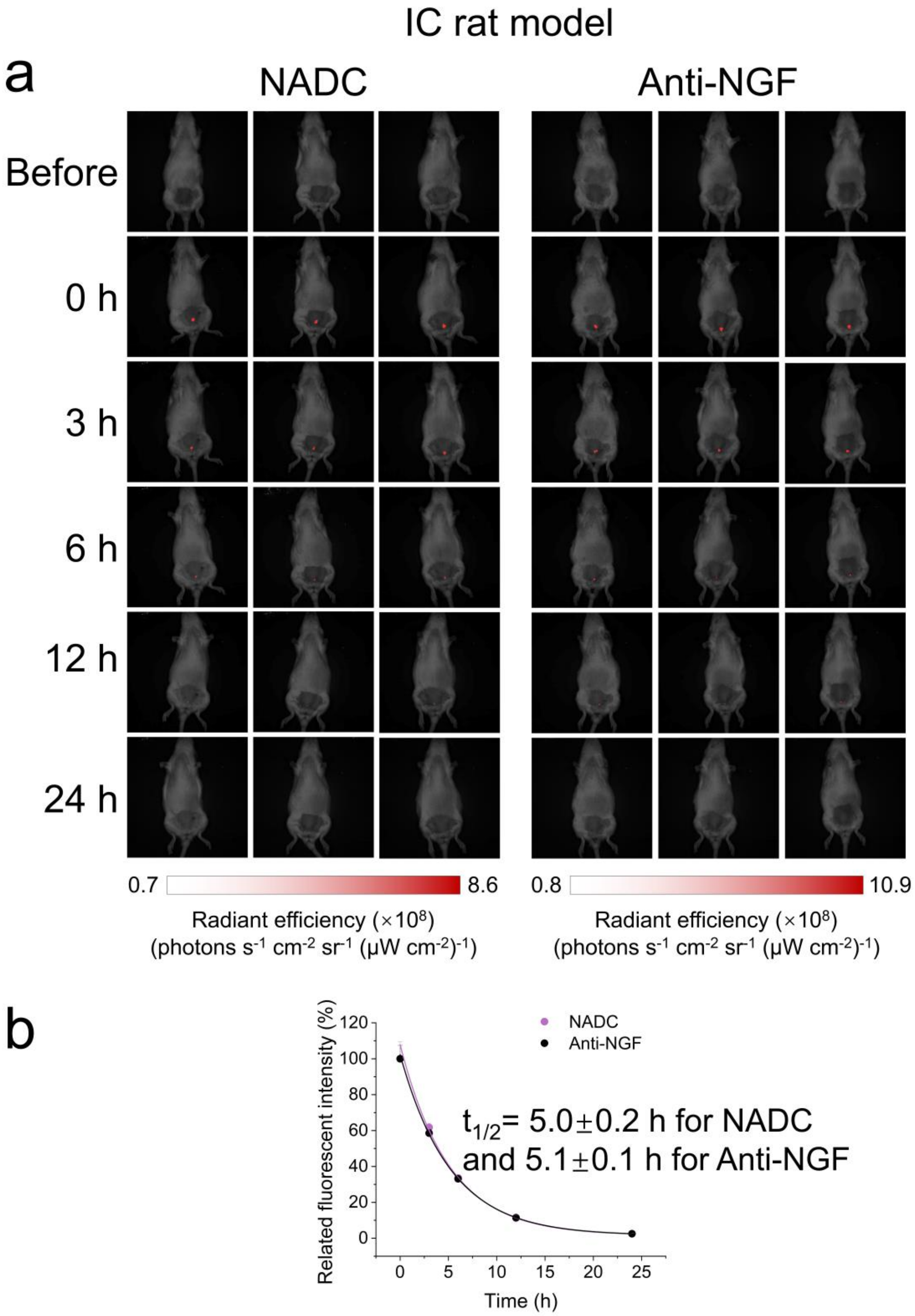


**Fig. S9.** Time-course in vivo NIRF imaging of rats with CYP-induced cystitis after intravesical administration of NADC or anti-NGF (n = 3 animals per group). (a) Corresponding NIRF images at indicated time points after intravesical injection. No statistically significant difference in the bladder retention half-lives of IC rats was perceived between NADC ($t_{1/2}$= 5.0 ± 0.2 h) and anti-NGF antibodies labeled with CF647 ($t_{1/2}$= 5.1 ± 0.1 h).

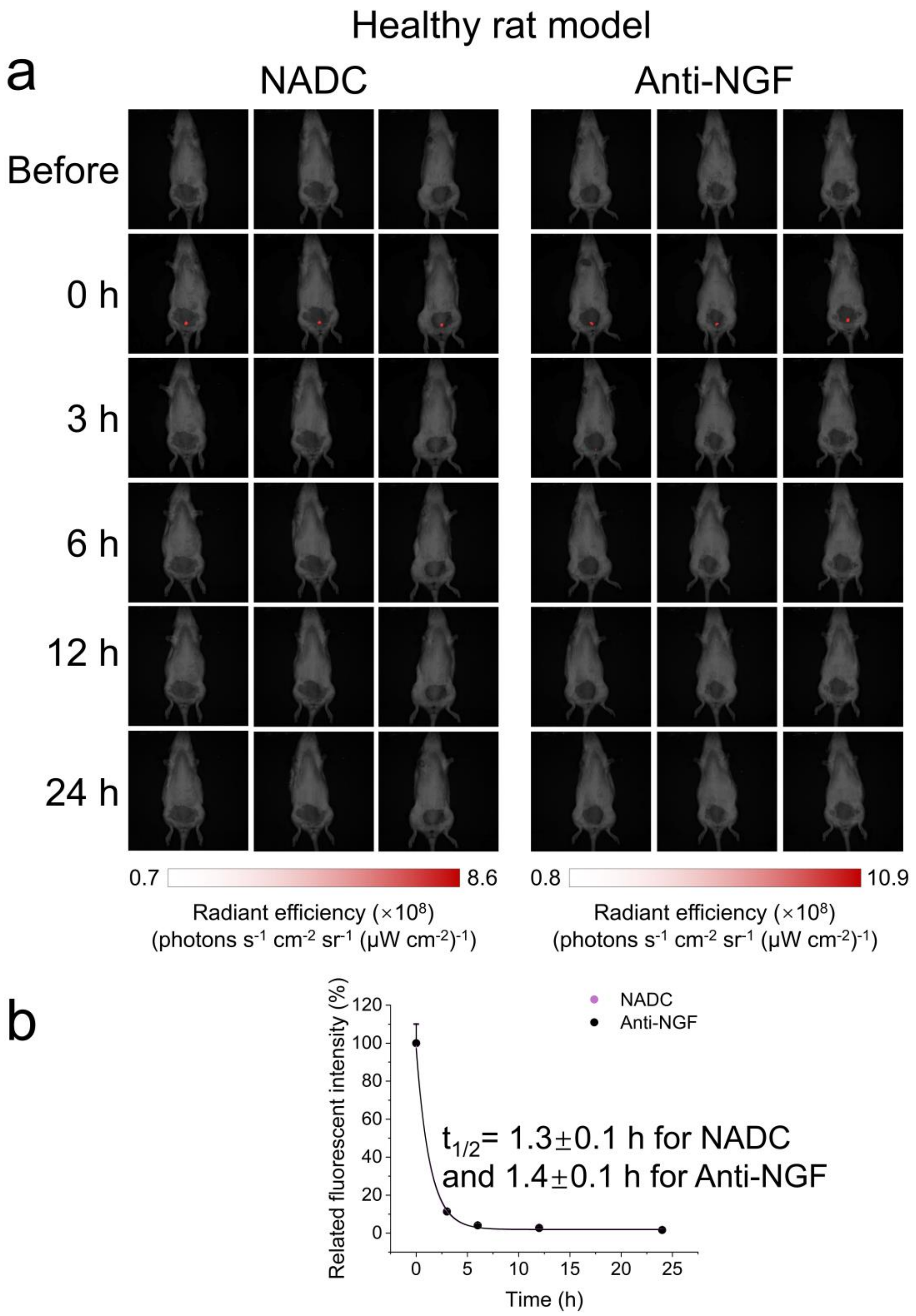


**Fig. S10.** Time-course in vivo NIRF imaging of healthy rats after intravesical administration of NADC or anti-NGF (n = 3 animals per group). (a) Corresponding NIRF images at indicated time points after intravesical injection. No statistically significant difference in the bladder retention half-lives of healthy rats was perceived between NADC ($t_{1/2}$= 1.3 ± 0.1 h) and Anti-NGF ($t_{1/2}$= 1.4 ± 0.1 h).

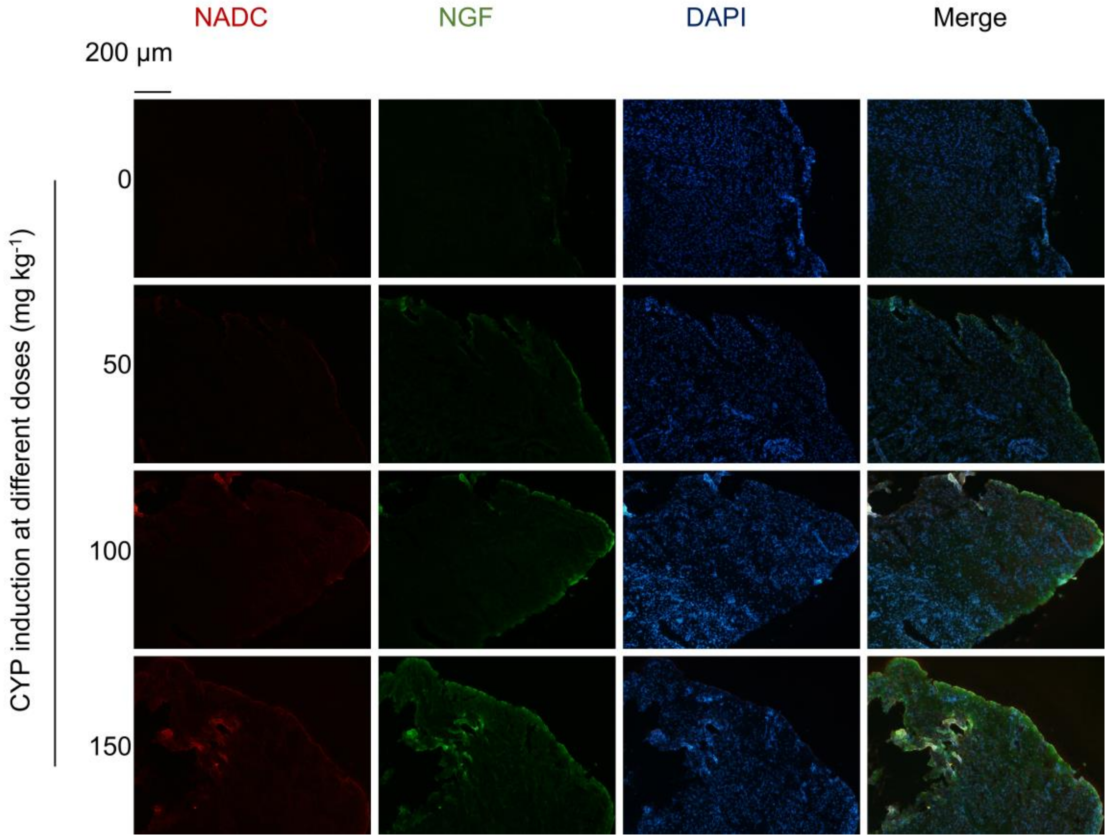


**Fig. S11. Immunofluorescence images of bladder tissues of IC rat models with varying CYP induction doses.** As the CYP induction dose increases, the NGF expression in the bladder mucosa is more pronounced. Additionally, NADC fluorescence shares a high extent of colocalization with NGF fluorescence, indicating specific targeting of NADC at lesions with high NGF expression in bladder tissues.

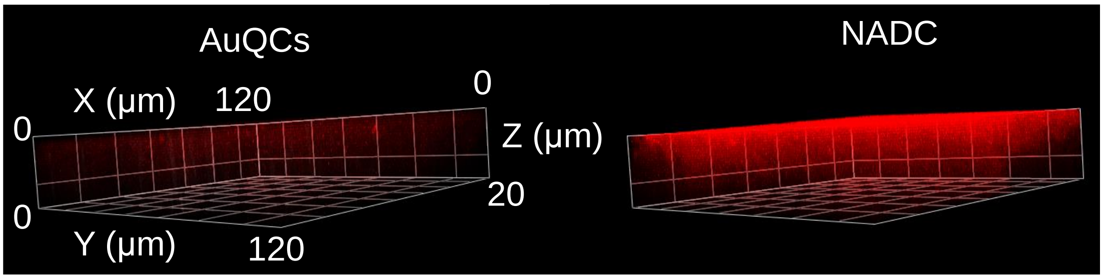


**Fig. S12. 3D Z-stack volumetric CLSM images of IC rat bladder tissue sections post-intravesical administration of AuQCs or NADC.** A more intense, broader, and deeper NIRF signal was observed for NADC.

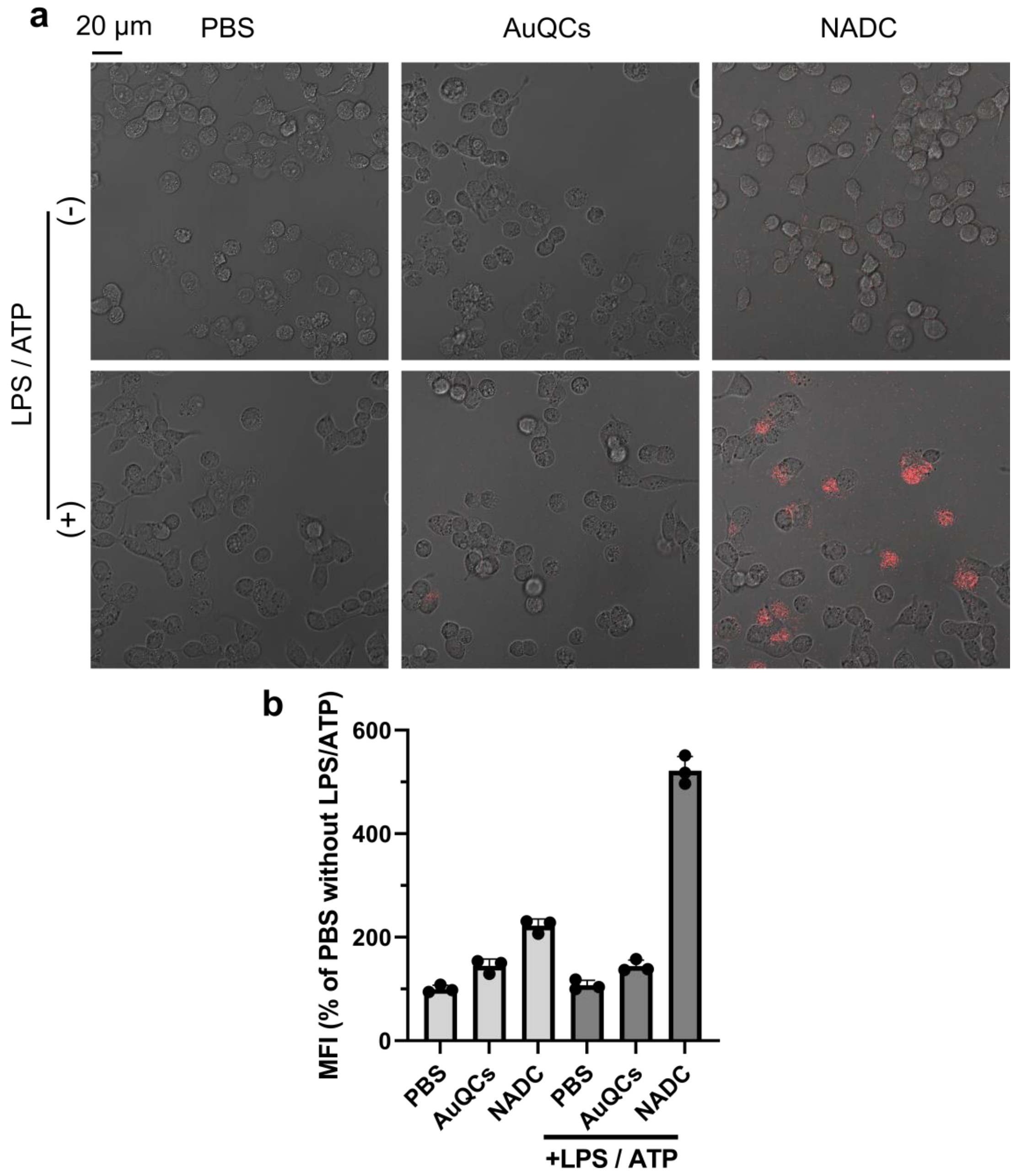


**Fig. S13. Fluorescence imaging of NADC in SV-HUC-1 cells following LPS/ATP-induced inflammation.** (a) Cells were treated by PBS, AuQCs, or NADC, with or without LPS/ATP stimulation. (b) Quantitative fluorescence analysis of acquired images (n = 3).

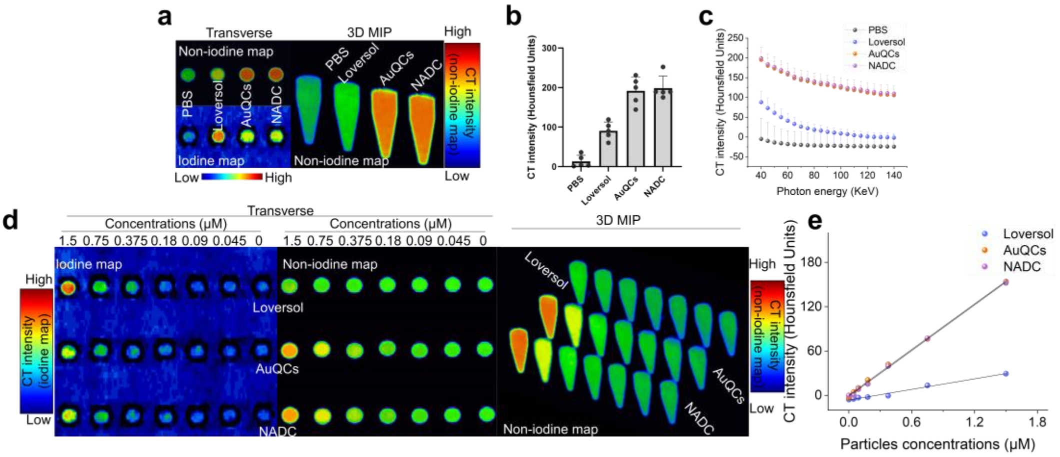


**Fig. S14. Phantom imaging by the dual-energy CT.** (a) Spectral CT images of AuQCs, NADC, and loversol imaging phantoms, reconstructed with the Gemstone Spectral Imaging software. (b) Quantification for the signal intensity of the non-iodine map (n = 5). (c) CT spectral curves of attenuation as a function of monochromatic energies spanning from 40-140 keV. (d) Spectral CT phantom images of loversol, AuQCs, and NADC at gradient concentrations. (e) The correlation between the concentration of contrast agents and the CT signal intensity.

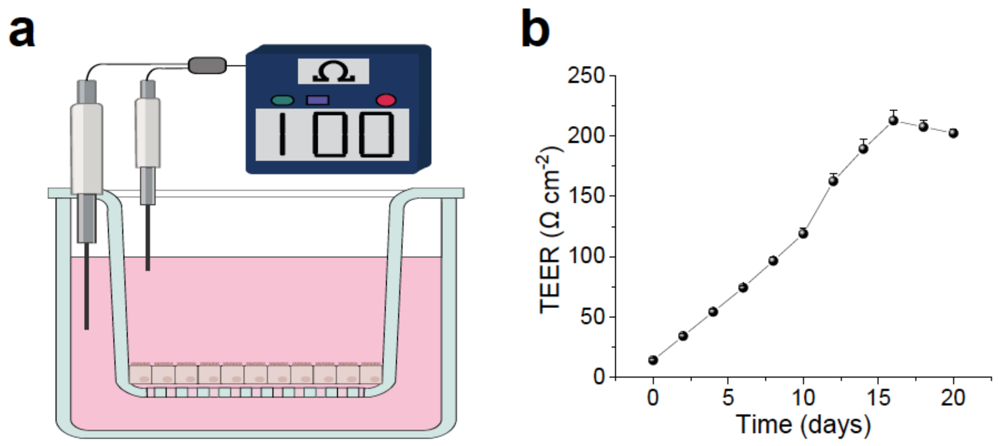


**Fig. S15. TEER measurement of SV-HUC-1 urothelium.** (a) A schematic diagram showing the setup of TEER measurement. (b) TEER values of the SV-HUC-1 monolayer urothelium mimicry over a period of 20 days reaching high confluence and saturation (n = 4). The SV-HUC-1 cells were continuously cultured to establish a cell monolayer, emulating the development of the bladder mucosal barrier. The emergence of tight junctions among the cells leads to an increment in resistance, which eventually stabilizes.

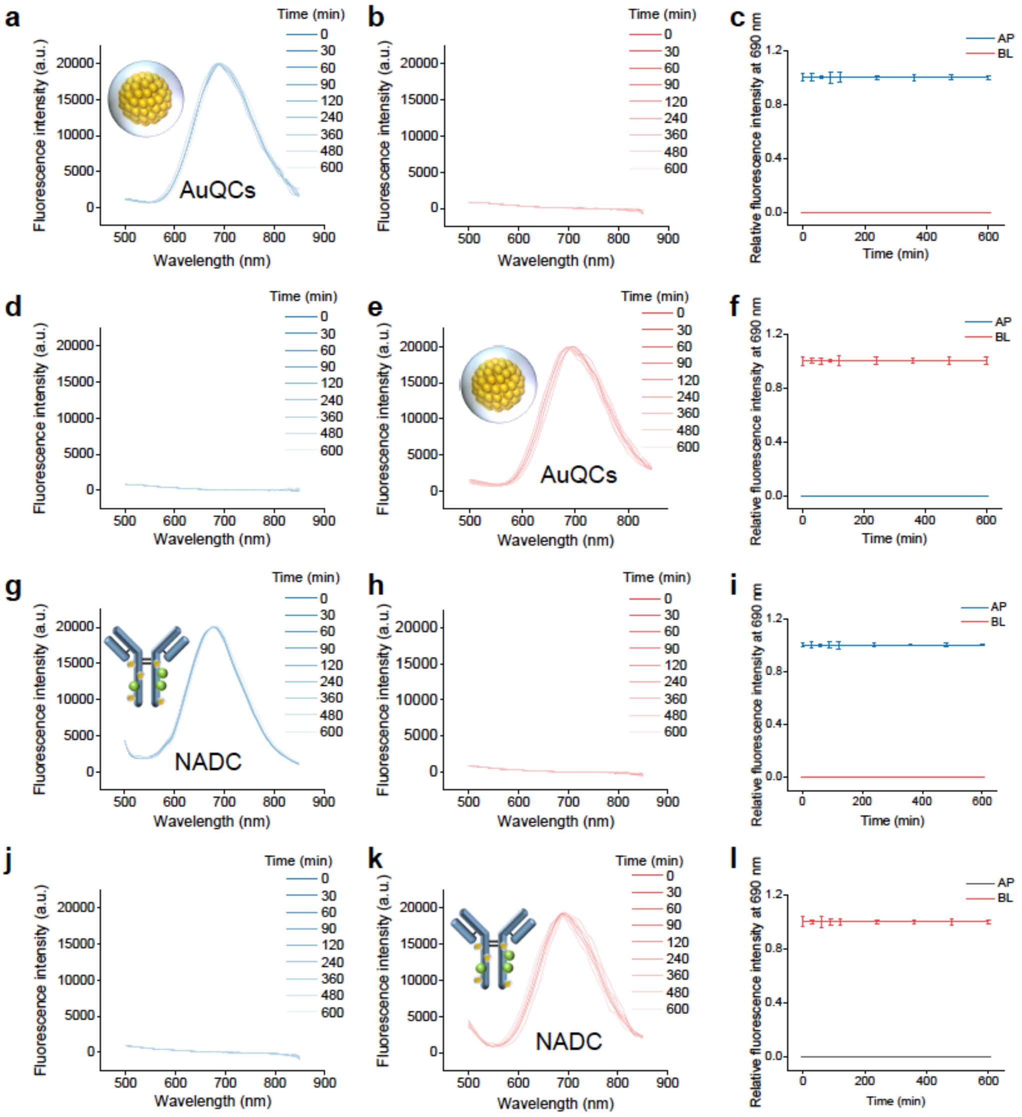


**Fig. S16. Evaluation of mucosal drug transport of NADC across the SV-HUC-1 urothelium mimicry by Ussing chamber tests.** (a-f) AuQCs were added to (a-c) the apical (AP) side (blue) or (d-f) the basolateral (BL) side (red), with samples collected at predetermined intervals to measure fluorescence spectra on both sides. Fluorescence spectra on (a, d) the AP side and (b, e) the BL side. (c, f) NIRF intensities on both sides as a function of time. (g-l) Similarly, NADC was introduced to either (g-i) the AP side or (j-l) the BL side. Fluorescence spectra on (g, j) the AP side and (h, k) the BL side. (i, l) NIRF intensities on both sides as a function of time.

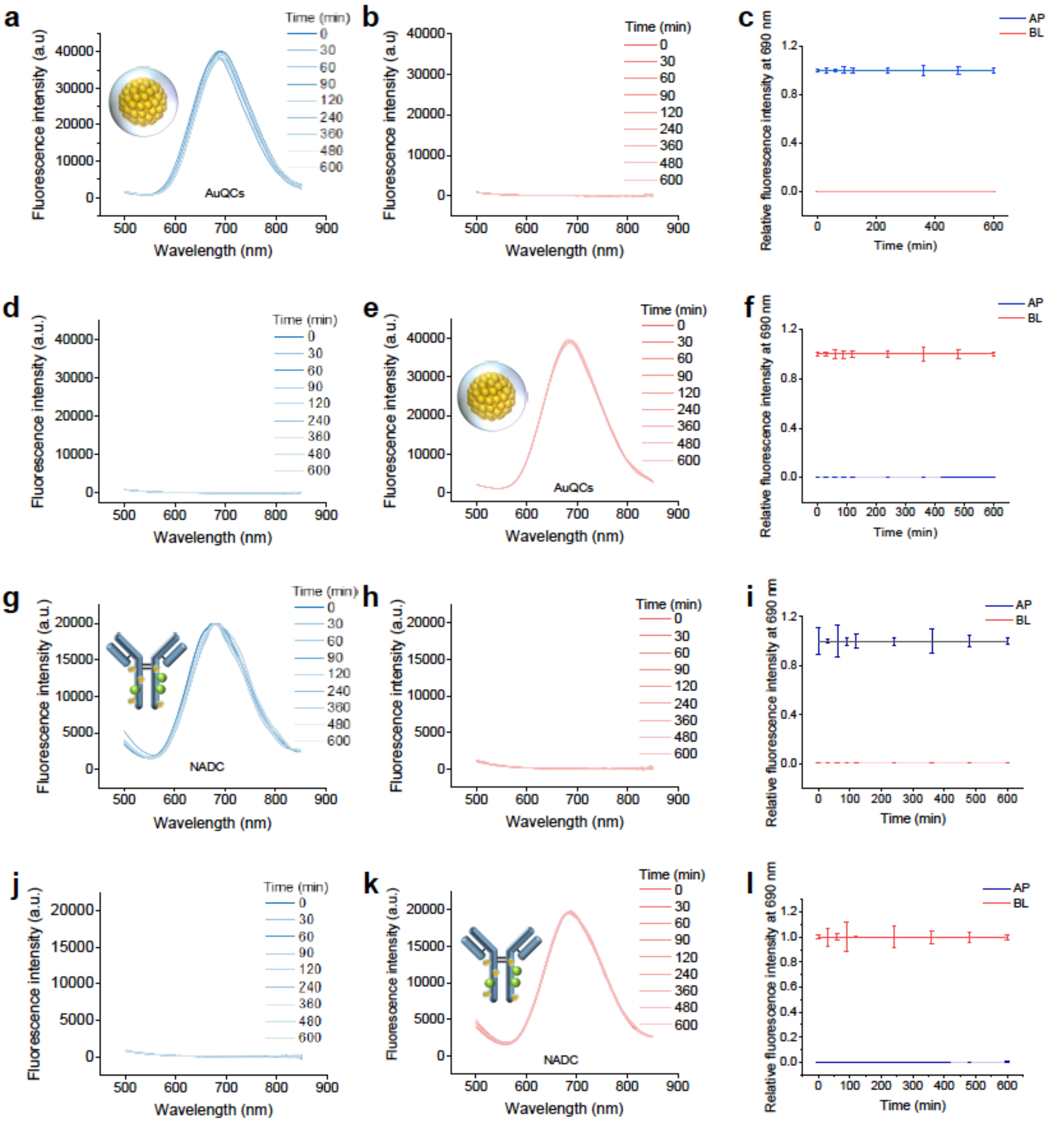


**Fig. S17. Evaluation of mucosal drug transport of NADC across healthy rat bladder tissues by Ussing chamber tests.** (a-f) AuQCs were added to (a-c) the apical (AP) side (blue) or (d-f) the basolateral (BL) side (red), with samples collected at predetermined intervals to measure fluorescence spectra on both sides. Fluorescence spectra on (a, d) the AP side and (b, e) the BL side. (c, f) NIRF intensities on both sides as a function of time. (g-l) Similarly, NADC was introduced to either (g-i) the AP side or (j-l) the BL side. Fluorescence spectra on (g, j) the AP side and (h, k) the BL side. (i, l) NIRF intensities on both sides as a function of time.

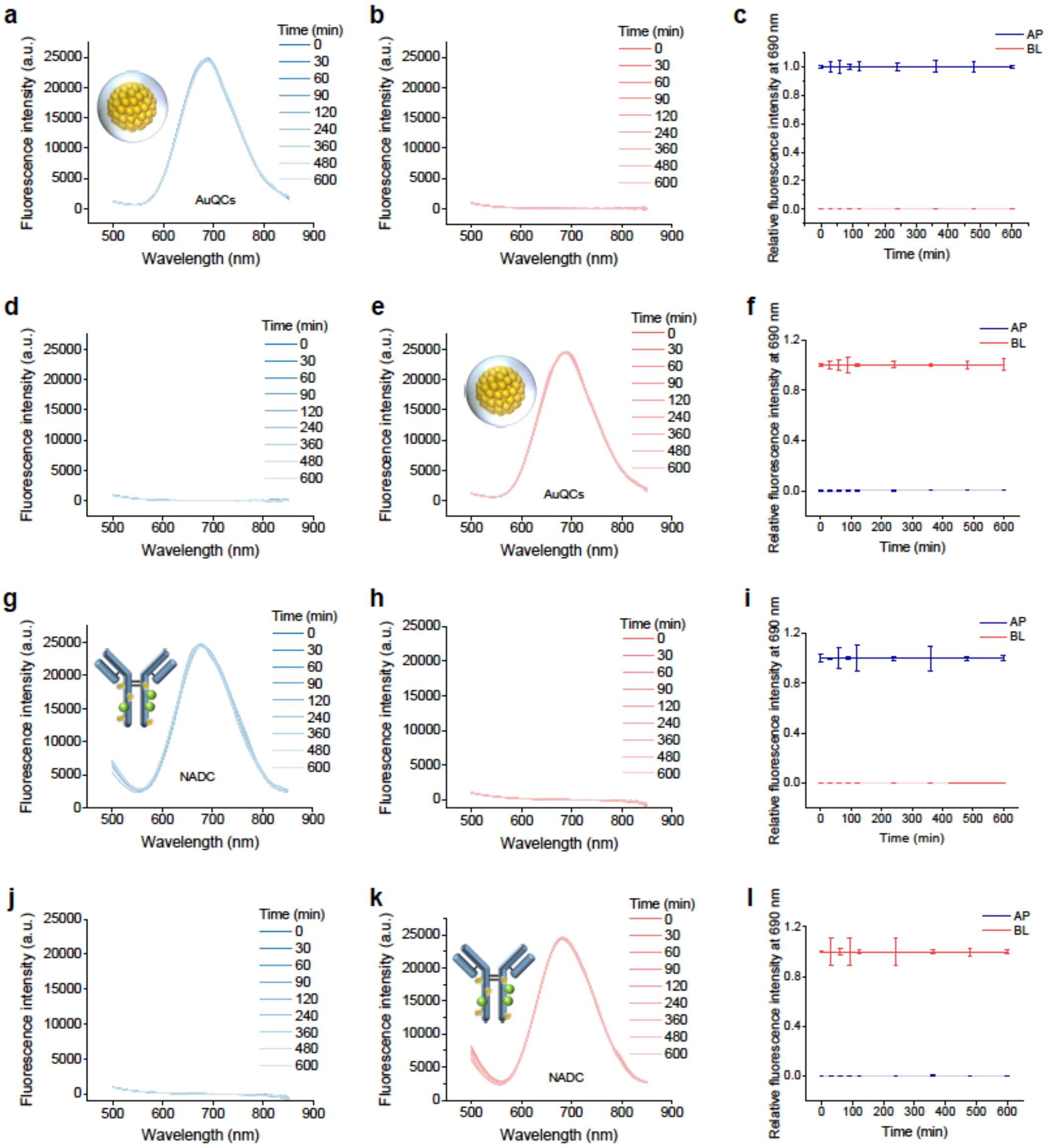

**Fig. S18. Evaluation of mucosal drug transport of NADC across rat bladder tissues with interstitial cystitis by Ussing chamber tests.** (a-f) AuQCs were added to (a-c) the apical (AP) side (blue) or (d-f) the basolateral (BL) side (red), with samples collected at predetermined intervals to measure fluorescence spectra on both sides. Fluorescence spectra on (a, d) the AP side and (b, e) the BL side. (c, f) NIRF intensities on both sides as a function of time. (g-l) Similarly, NADC was introduced to either (g-i) the AP side or (j-l) the BL side. Fluorescence spectra on (g, j) the AP side and (h, k) the BL side. (i, l) NIRF intensities on both sides as a function of time.

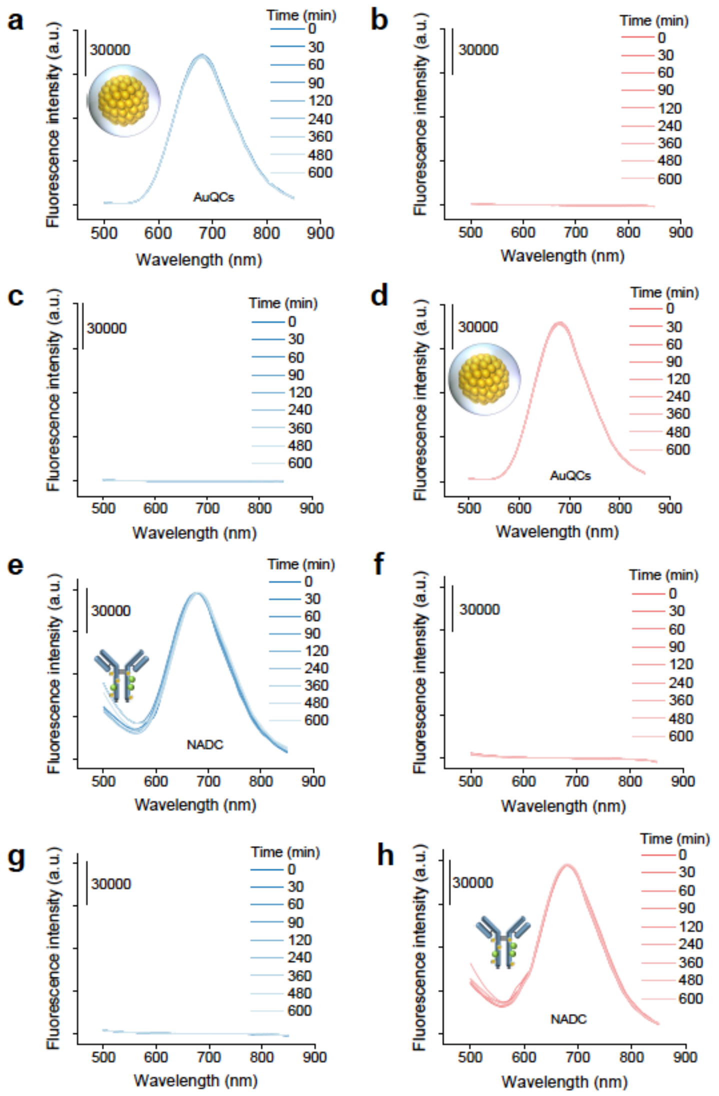


**Fig. S19. Evaluation of mucosal drug transport of NADC across human bladder tissues by Ussing chamber tests.** AuQCs were added to (a-b) the apical (AP) side (blue) or (c-d) the basolateral (BL) side (red), with samples collected at predetermined intervals to measure fluorescence spectra on both sides. (e-h) Similarly, NADC was introduced to either (e-f) the AP side or (g-h) the BL side.

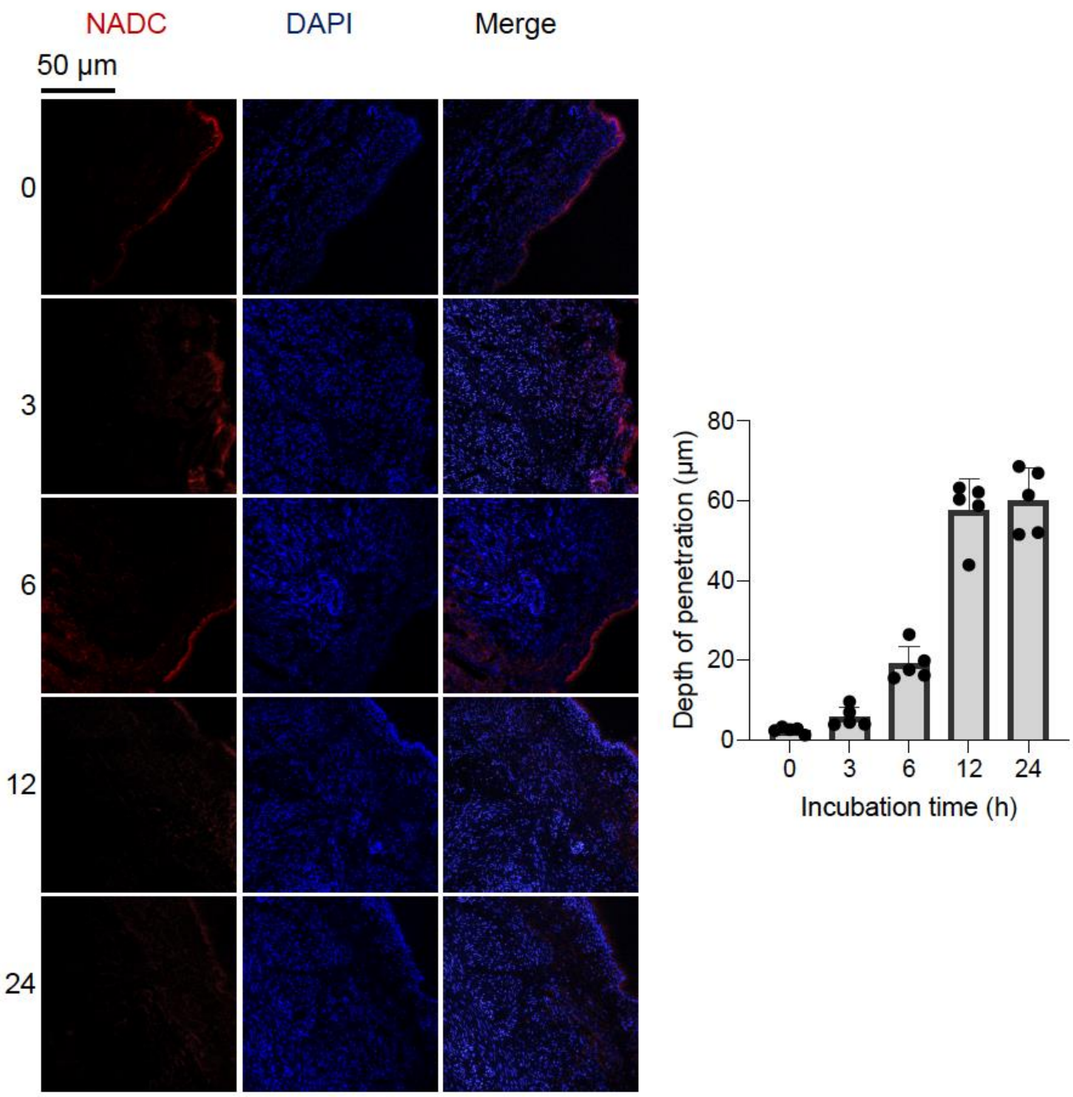


**Fig. S20. The penetration depth of NADC in the bladder.** The penetration depth of NADC in the bladder was assessed at various time points following intravesical instillation of a consistent NADC dose. Rats with IC were euthanized, with bladder tissues harvested, freeze-embedded for longitudinal sections, and stained with DAPI counterstain to highlight nuclei. NADC gradually penetrated the bladder mucosal tissue over time, reaching a maximum depth of nearly 60 microns beneath the mucosa in 12 h (n = 5).

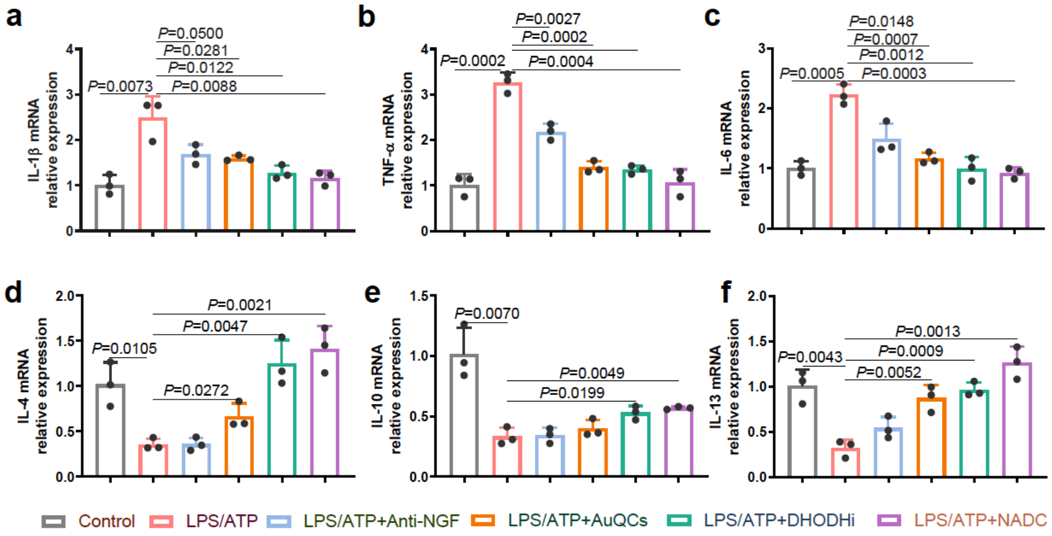


**Fig. S21. The mRNA expression profiles of pro- and anti-inflammatory cytokines by RT-qPCR in NADC-treated SV-HUC-1 cells with LPS/ATP-induced inflammation.** The relative expression levels of (a) IL-1β, (b) TNF-α, (c) IL-6, (d) IL-4, (e) IL-10, and (f) IL-13 (n = 3). Co-stimulation of SV-HUC-1 cells with LPS/ATP simultaneously led to an increase in the expression of pro-inflammatory cytokines like IL-1β, TNF-α, and IL-6 and a decrease in the expression of anti-inflammatory cytokines like IL-4, IL-10, and IL-13, which were notably reversed towards healthy control levels through NADC treatment.

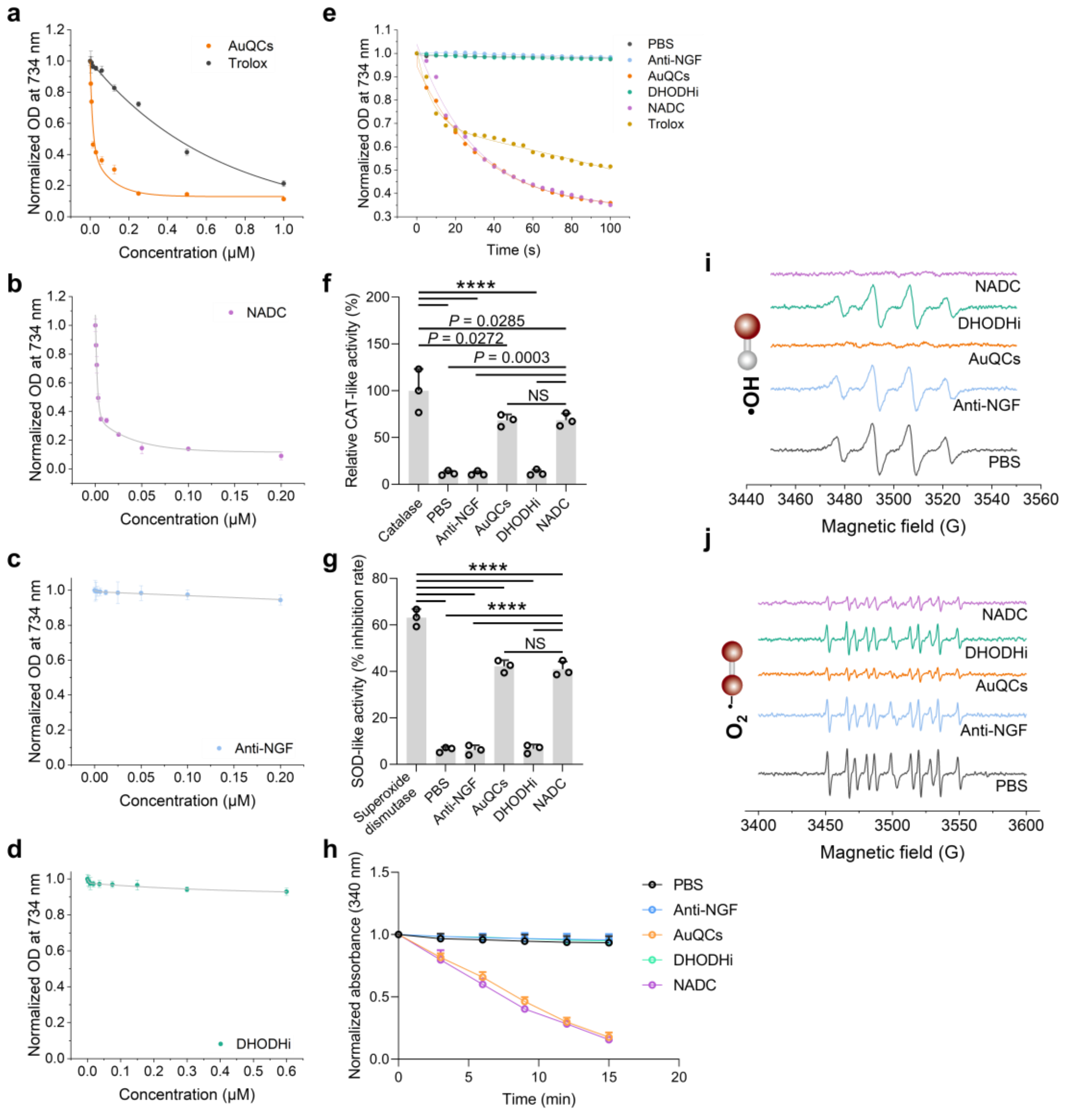


**Fig. S22. ROS scavenging and enzyme-mimicking activities of AuQCs in NADC.** (a-d) Total antioxidant capacity (T-AOC) colorimetric assays of gradient concentrations of AuQCs, Trolox, NADC, anti-NGF, and DHODHi against the $ABTS^{\bullet +}$ radical cation. (e) Dynamic scavenging curves of T-AOC for indicated agents by measuring time-dependent changes in absorbance at 734 nm. (f-g) CAT-, SOD-, and GPx-mimicking activities. Electron spin resonance (ESR) spectra of various samples with (i) DMPO or (j) DEPMPO spin traps for hydroxyl radicals (·OH) and superoxide anion ($O_2^{\cdot -}$), respectively.

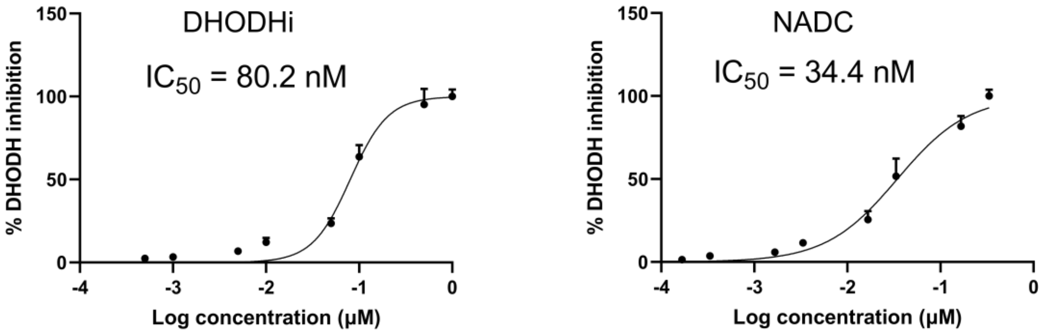


**Fig. S23. In vitro DHODH enzyme inhibition assay.** A substrate mixture of *L*-DHO, DUQ, and DCIP was added to initiate the enzymatic reaction and evaluate the inhibition by DHODHi or NADC. Decreased absorbance at 600 nm was measured in response to the reduction of DCIP.

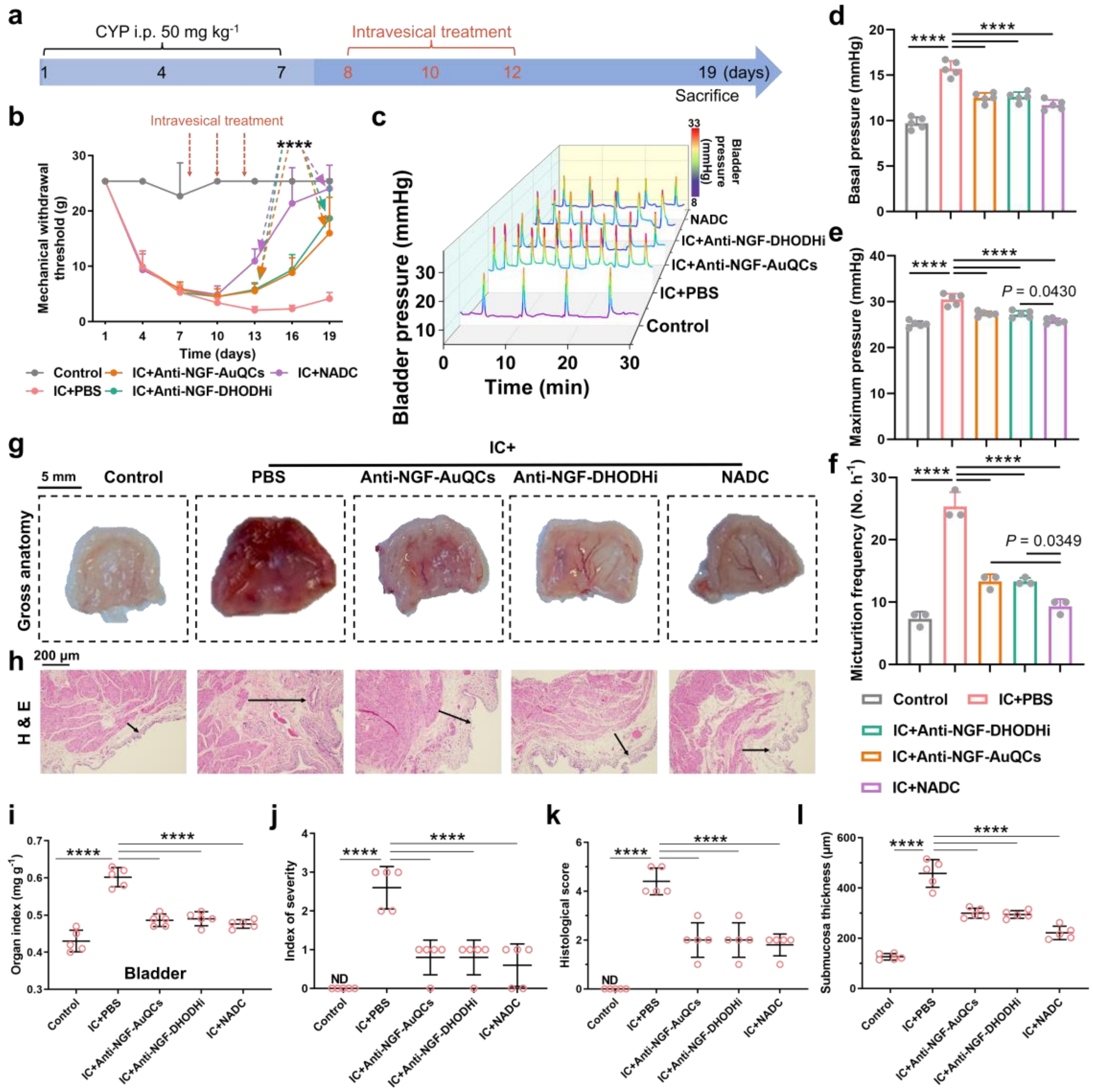


**Fig. S24. Comparison of NADC with anti-NGF-AuQCs and anti-NGF-DHODHi in treating chronic IC in rats.** (a) Schematic workflow of chronic IC induction and therapeutic interventions in rats. (b) Time-course evaluation of mechanical allodynia thresholds in IC rats treated with indicated therapies. All statistical analyses between groups in $P$ values were compared with the IC+PBS group (n=5). (c) Representative real-time bladder urodynamic curves of rats with chronic IC across treatment groups. (d) The basal pressure, (e) maximum pressure, and (f) micturition frequency quantified from urodynamic analyses. (g) Gross morphological appearance of bladders following intravesical therapy. (h) Hematoxylin and eosin (H&E)-stained bladder sections. Scale bar: 200 μm. Arrows denote mucosal edema extent with length proportional to severity. (i) Bladder organ index defined by the bladder-to-body weight ratio, (j) index of severity, (k) histological scores, and (l) submucosa thickness (n=5). Statistical significance levels: ****$P \leq 0.0001$.

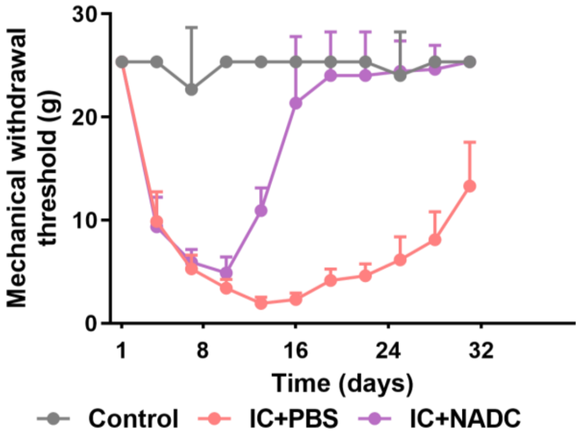


**Fig. S25. Changes in the mechanical withdrawal threshold over one-month duration.** Intravesical NADC interventions effectively relieved pain symptoms in rats with chronic IC and capably sustained the efficacy for over a month without recurrence, while the symptomatic relief with PBS treatment was insignificant in comparison.

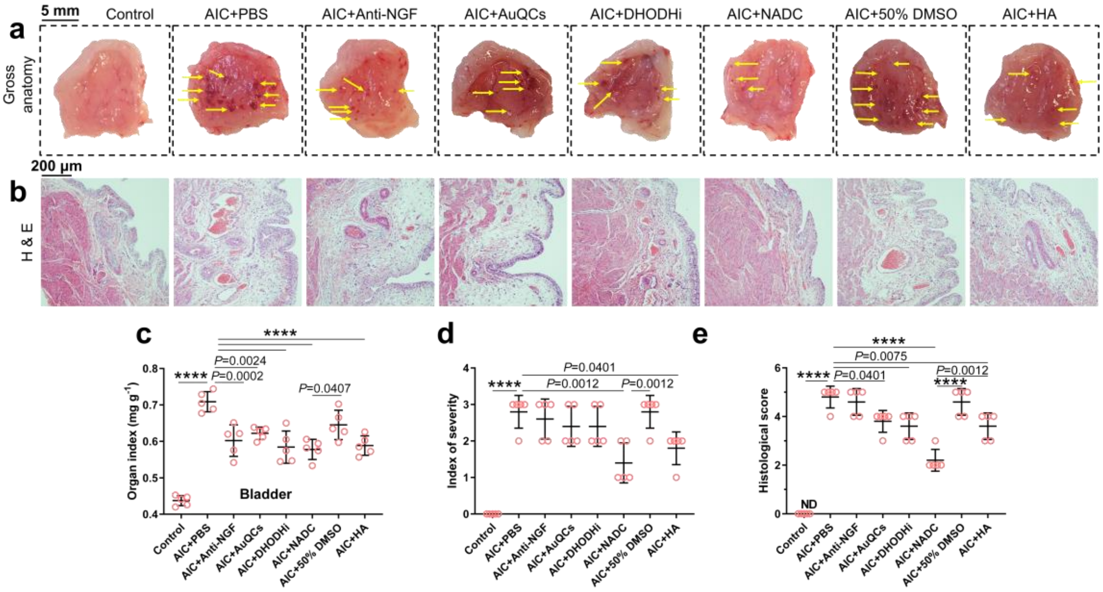


**Fig. S26. Histopathology of bladder tissue sections from AIC rats with intravesical NADC treatment.** (a) Gross anatomy of AIC rat bladders harvested at the therapeutic endpoint of intravesical infusion. NADC significantly mitigated AIC symptoms of congestion, bleeding, and edema, as indicated by yellow arrows. (b) H&E staining of rat bladder sections in indicated treatment groups. (c) The bladder organ index, (d) index of severity, and (e) histological scores. Statistical significance levels (n = 5): ****$P \leq 0.0001$.

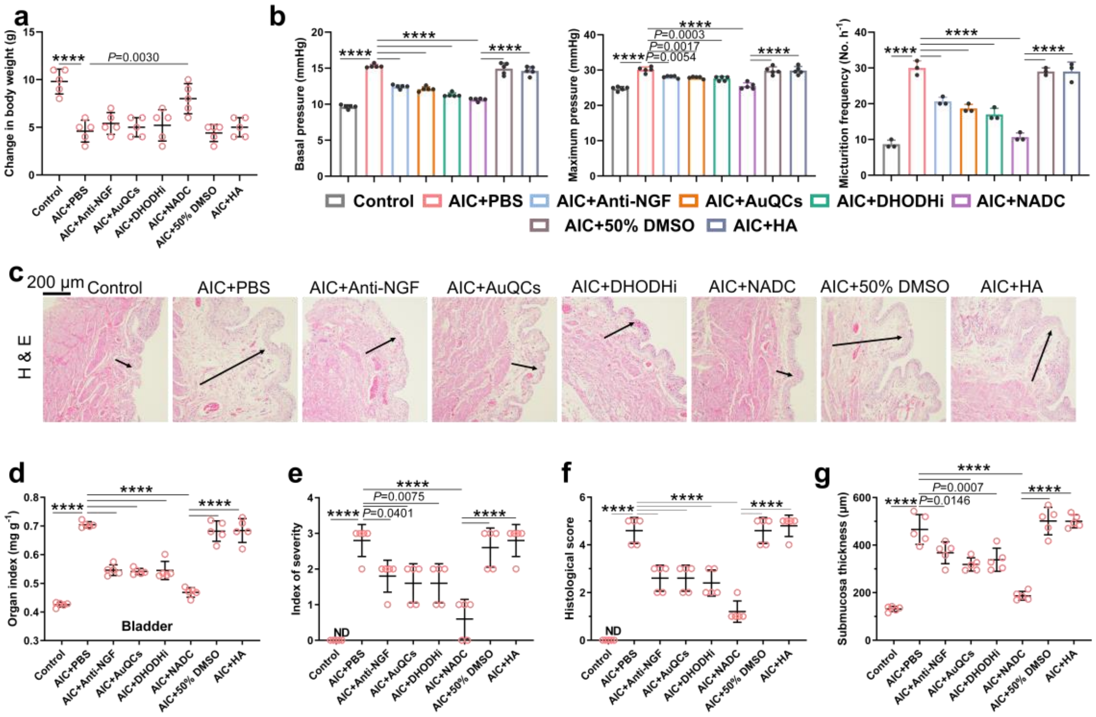


**Fig. S27. Additional data of NADC as a prophylactic agent for AIC in rats.** (a) Body weight changes of AIC rats receiving prophylactic treatment at the study endpoint (n=5). (b) Real-time bladder urodynamic curves of AIC rats with prophylactic treatment. The basal pressure, maximum pressure, and micturition frequency of bladder urodynamics in AIC rats receiving pre-administration of intravesical prophylaxis (n=3 to 5 animals per group). (c) Histopathology of resected bladders in AIC rats with pre-administered prophylactic treatment by H&E staining. (d) Bladder organ index, (e) index of severity, (f) histological scores and (g) submucosa thickness (n=5). Statistical significance levels: ****$P \leq 0.0001$.

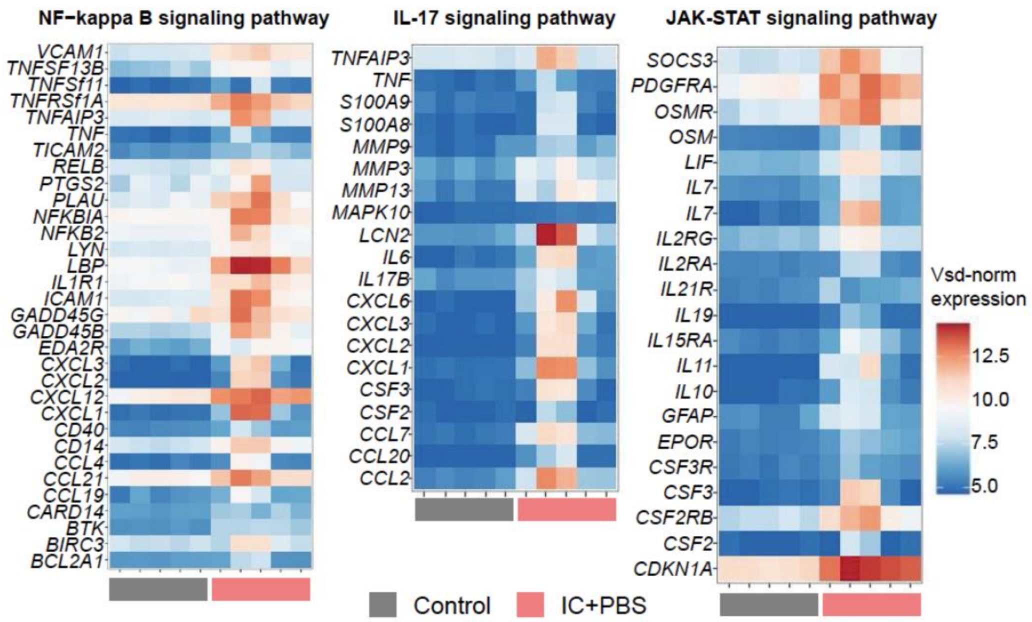


**Fig. S28. Heatmaps of significantly altered key effector genes in the NF-κB, IL-17, and JAK-STAT pathways in rat bladder tissues between the control and IC groups.** Expression values from high to low are displayed in color gradients ranging from red to blue (n = 5).

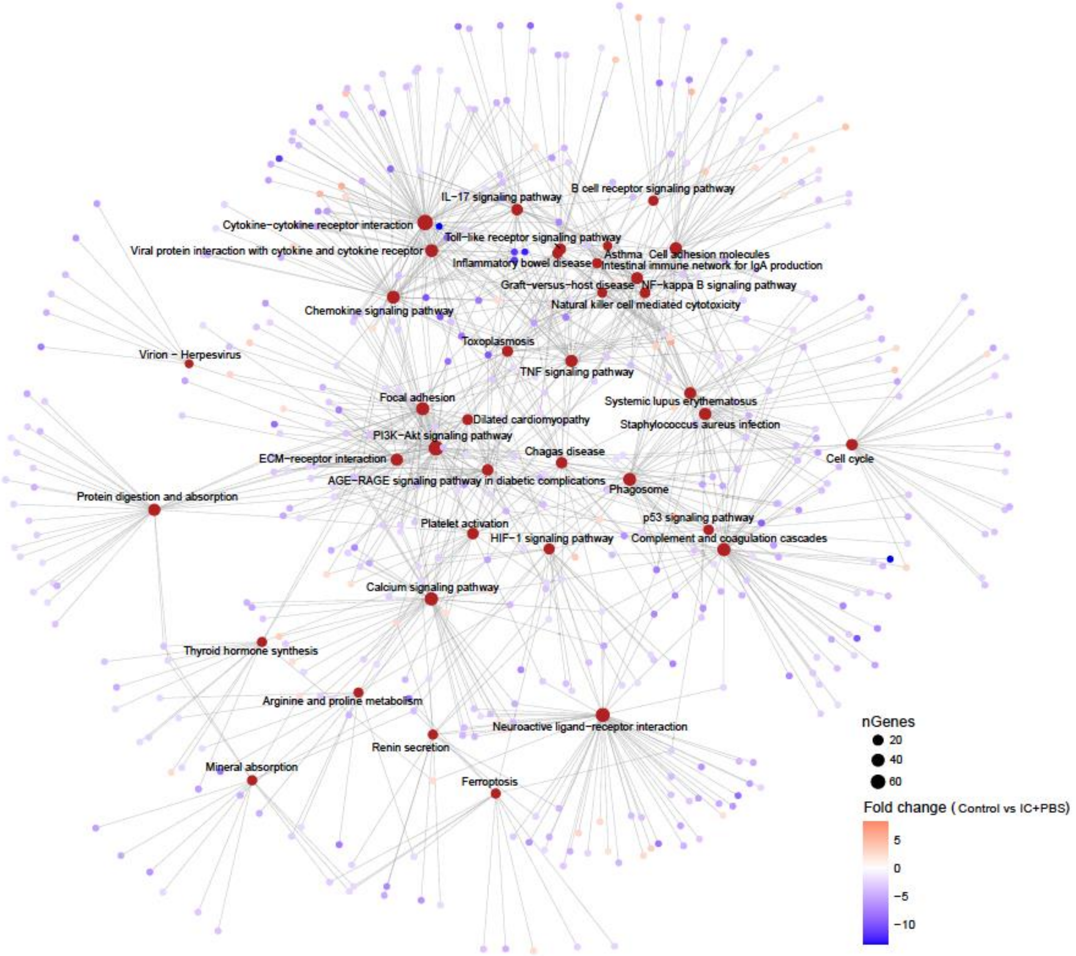


**Fig. S29. Protein-protein interaction (PPI) network of differentially expressed proteins between the control and IC groups.** Key nodes in the complex network of biological functions are highlighted in colors denoting fold changes. A node's size corresponds to its betweenness centrality (BC) value.

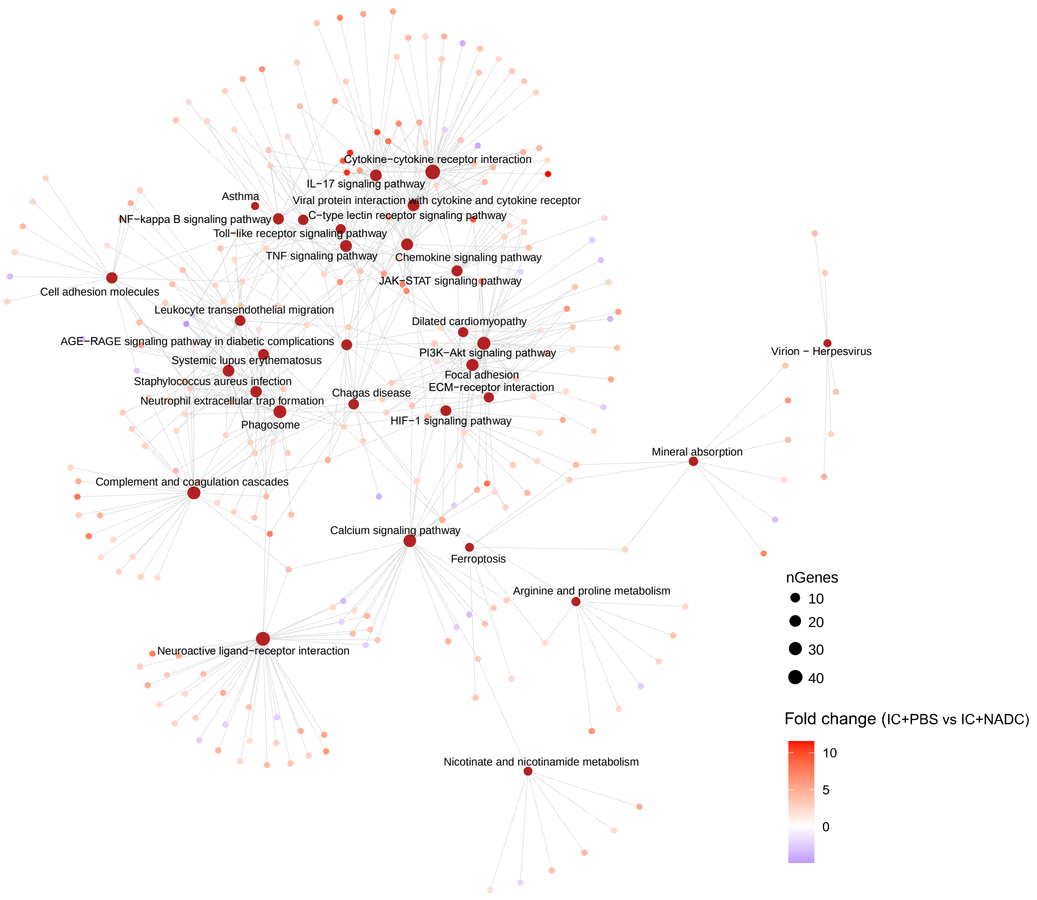


**Fig. S30. PPI network analysis of differentially expressed proteins in IC rat bladder tissues with or without NADC treatment.** Key nodes are highlighted with colors and sizes denoting fold changes and BC values, respectively.

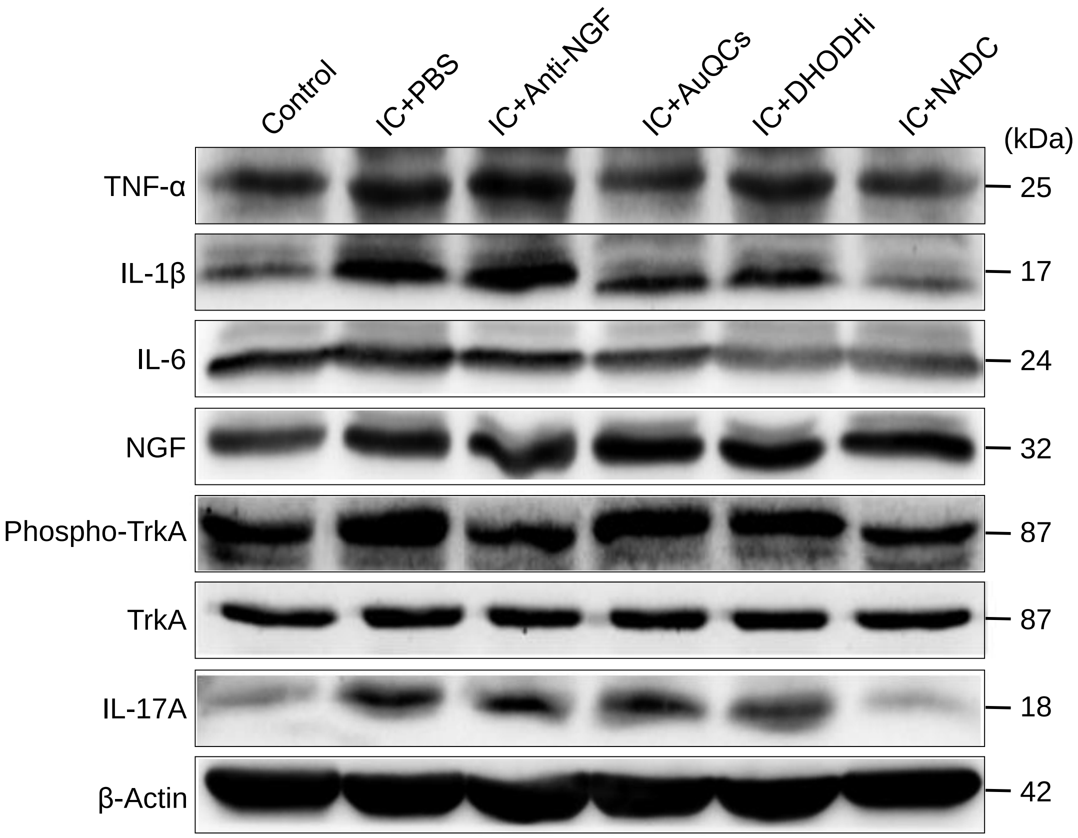


**Fig. S31. Western blots of molecules involved in inflammation pathways in bladder tissues of IC rats.** In bladder tissues of IC rats, pro-inflammatory cytokines were upregulated in contrast to the healthy control, including TNF-α, IL-1β, and IL-6. Notably, aberrant NGF overexpression in response to IC occurrence in rat bladders was not inhibited by any agents, including the NGF-targeting tanezumab and NADC. The phosphorylation of the transmembrane NGF catalytic receptor and the nociceptive mediator tropomyosin receptor kinase A (TrkA) experienced an increase in IC rat models but was inhibited by both tanezumab and NADC. As an upstream regulator, IL-17A was a known target of vidofludimus that inhibited IL-17A expression. β-Actin served as an internal control for loading, with molecular mass markers denoted on the right, expressed in kilodaltons (kDa).

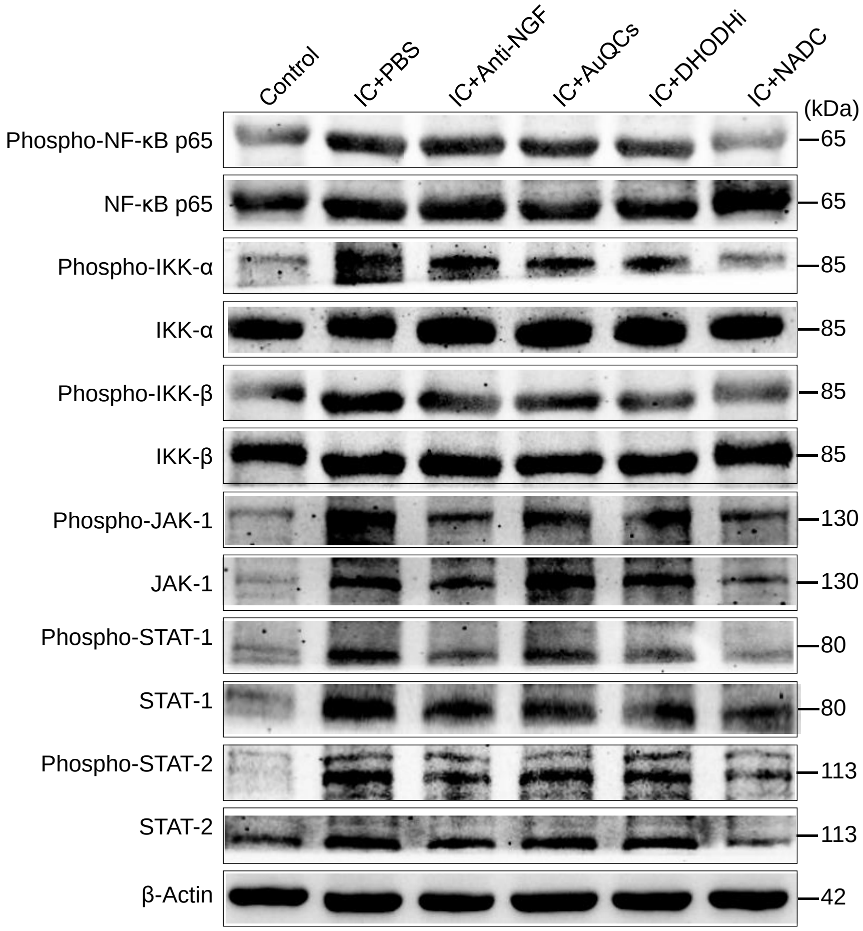


**Fig. S32. Western blots of key molecules in NF-κB and JAK-STAT signaling pathways.** A noted increase was observed in the phosphorylation of two IκB kinase (IKK) catalytic subunits, IKKα and IKKβ, as well as RelA/p65, leading to the activation of the classical NF-κB pathway in rat IC models. The NF-κB activation was effectively attenuated by intravesical NADC. Likewise, in the context of IC, the dysregulation of JAK-STAT regulators broadly implicated in cytokine signaling, including JAK1, STAT1, and STAT2, was significantly mitigated by NADC, demonstrating potent synergy in inhibiting both phosphorylation and expression. β-Actin served as an internal control for loading, with molecular mass markers denoted in kDa.

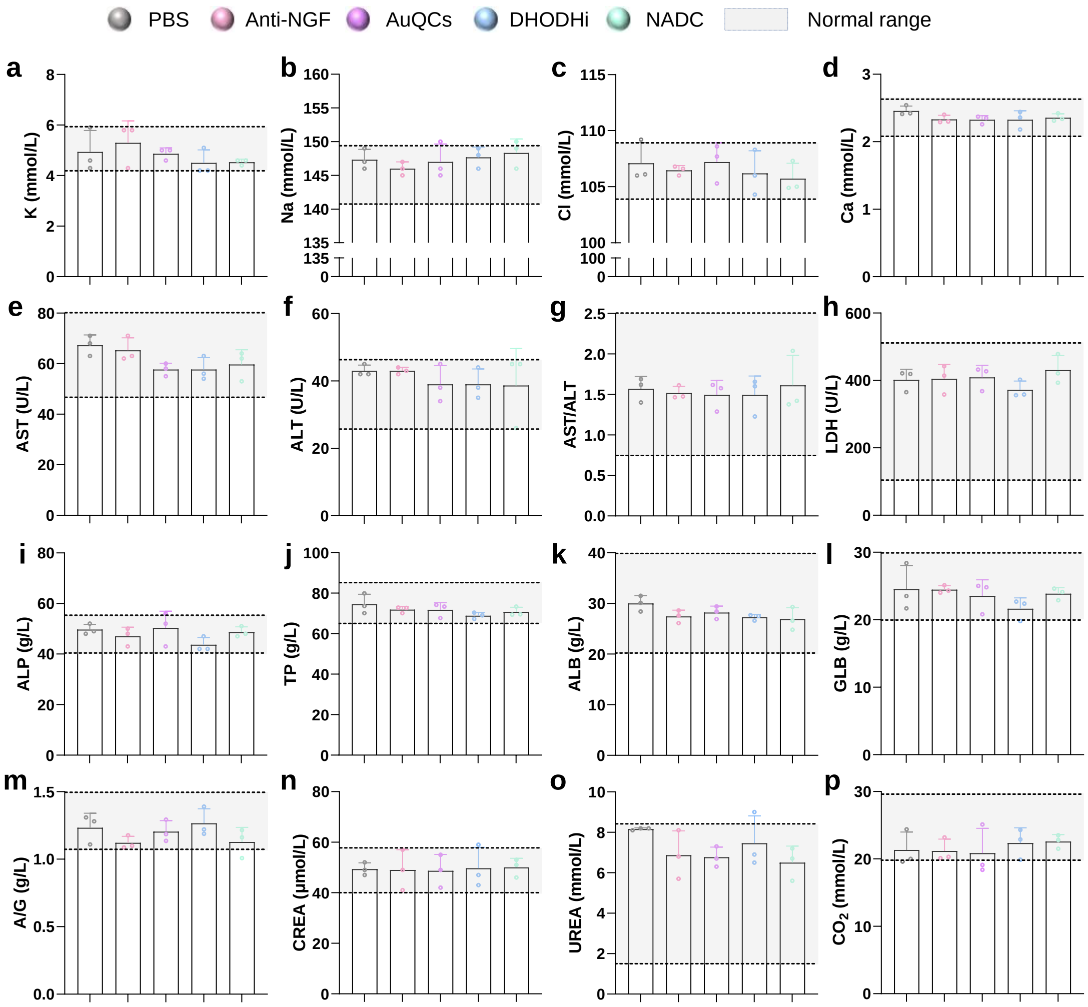


**Fig. S33. Clinical chemistry analysis.** The configuration of indicators is delineated as follows (n=3): (a) potassium, (b) sodium, (c) chloride, (d) calcium, (e) aspartate aminotransferase (AST), (f) alanine aminotransferase (ALT), (g) AST/ALT, (h) lactate dehydrogenase (LDH), (i) alkaline phosphatase (ALP), (j) total protein (TP), (k) albumin (ALB), (l) globulin (GLB), (m) ALB/GLB, A/G, (n) creatinine (CREA), (o) urea nitrogen (UREA), (p) carbon dioxide ($CO_2$). All indicators fell within normal ranges (gray regions) with no statistically significant difference from healthy control rats.

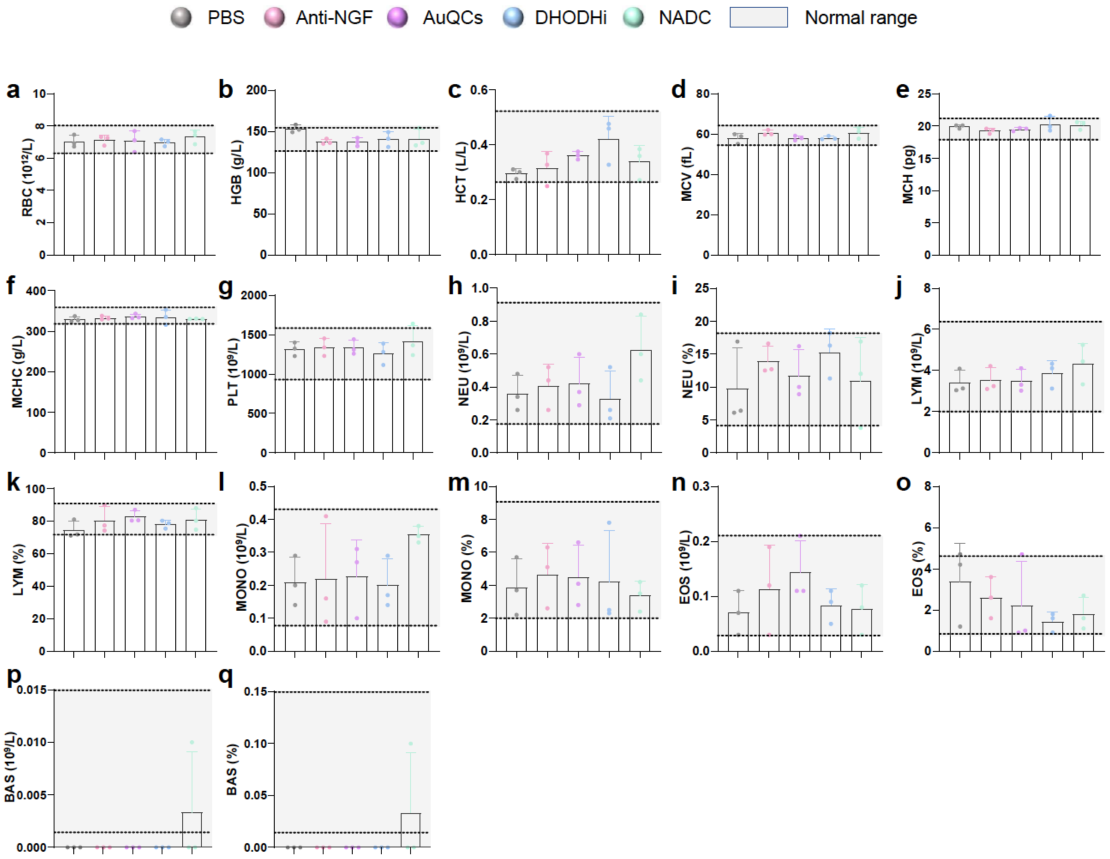


**Fig. S34. Complete blood count with automated differentials.** The configuration of indicators is delineated as follows (n=3): (a) red blood cell count (RBC), (b) hemoglobin (HGB), (c) hematocrit (HCT), (d) mean corpuscular volume (MCV), (e) mean corpuscular hemoglobin (MCH), (f) mean corpuscular hemoglobin concentration (MCHC), (g) platelet count (PLT), (h) neutrophil count (NEU), (i) neutrophil percentage, (j) lymphocyte count (LYM), (k) lymphocyte percentage, (l) monocyte count (MONO), (m) monocyte percentage, (n) eosinophil count (EOS), (o) eosinophil percentage, (p) basophil count (BAS), and (q) basophil percentage. All indicators fell within normal ranges (gray regions) with no statistically significant difference from healthy control rats.
